# Random access DNA memory in a scalable, archival file storage system

**DOI:** 10.1101/2020.02.05.936369

**Authors:** James L. Banal, Tyson R. Shepherd, Joseph Berleant, Hellen Huang, Miguel Reyes, Cheri M. Ackerman, Paul C. Blainey, Mark Bathe

## Abstract

DNA is an ultra-high-density storage medium that could meet exponentially growing worldwide demand for archival data storage if DNA synthesis costs declined sufficiently and random access of files within exabyte-to-yottabyte-scale DNA data pools were feasible. To overcome the second barrier, here we encapsulate data-encoding DNA file sequences within impervious silica capsules that are surface-labeled with single-stranded DNA barcodes. Barcodes are chosen to represent file metadata, enabling efficient and direct selection of sets of files with Boolean logic. We demonstrate random access of image files from an image database using fluorescence sorting with selection sensitivity of 1 in 10^6^ files, which thereby enables 1 in 10^6*N*^ per *N* optical channels. Our strategy thereby offers retrieval of random file subsets from exabyte and larger-scale long-term DNA file storage databases, offering a scalable solution for random-access of archival files in massive molecular datasets.

## INTRODUCTION

While DNA is conventionally the polymer used for storage and transmission of genetic information in biology, it can also be used for the storage of arbitrary digital information at densities far exceeding conventional data storage technologies such as flash and tape memory, at scales well beyond the capacity of the largest current data centers^1,2^. Recent progress in nucleic acid synthesis and sequencing technologies continue to reduce the cost of writing and reading DNA, thereby rendering DNA-based information storage potentially viable commercially in the future^3-6^. Demonstrations of its viability as a general information storage medium include numerous examples including the storage and retrieval of books, images, computer programs, audio clips, works of art, and Shakespeare’s sonnets using a variety of encoding schemes^7-13^, with data size limited primarily by the cost of DNA synthesis. In each case, digital information was converted to DNA sequences composed of ∼100–200 nucleotide (nt) data blocks for ease of chemical synthesis and sequencing. Sequence fragments were then assembled to reconstruct the original, encoded information.

While significant effort in DNA data storage has focused on increasing the scale of DNA synthesis, as well as improving encoding schemes, an additional crucial aspect of a successful molecular data storage system is the ability to efficiently retrieve specific files, or random subsets of files, from a large-scale pool of DNA data on demand, without error, without data destruction, and ideally at low cost for a practical archival data storage and retrieval device. Toward this end, to date research has largely used conventional polymerase chain reaction (PCR)^9,11,13^, which uses up to 20–30 heating and cooling cycles with DNA polymerase to selectively amplify and extract specific DNA sequences from a DNA data pool using primers. Nested addressing barcodes^14-16^ have also been used to uniquely identify a greater number of files, as well as biochemical affinity tags to selectively pull down oligos for targeted amplification^17^.

Major limitations of PCR-based approaches, however, include the length of DNA needed to uniquely label DNA data strands for file indexing, which dramatically reduces the DNA available for data storage. For example, for an exabyte-scale data pool, each file requires at least three barcodes^17^, or up to sixty nucleotides in total barcode sequence length, thereby reducing the number of nucleotides that can be used for data encoding. Further, selective amplification of a specific file using PCR requires access to the entire data pool for each query, which is destructive to the data pool, and intrinsically limited by the finite number orthogonal primers, e.g., 28,000 for previously demonstrated PCR-based random access system^13^, available to amplify target files without strand crosstalk due to non-specific hybridization. Finally, PCR-based approaches do not allow for physical deletion of specific files from a data pool and require numerous heating and cooling cycles with DNA polymerase, which may be prohibitively costly, time-consuming, and impractical for random access memory in exabyte-to-yottabyte-scale data pools. While spatial segregation of data into distinct pools^18^ and extraction of selected DNA using biochemical affinity pulldown have yielded significant improvements in PCR-based file selection strategies, these implementations vastly reduce data density^17^, and cannot access random subsets of files in this direct manner that is required for a truly scalable and deployable archival molecular file storage and retrieval system.

As an alternative to PCR-based approaches, here we focus on archival DNA data storage and retrieval by first encapsulating physically DNA-based files within discrete, impervious silica capsules, which we subsequently label with single-stranded DNA barcodes that enable direct, random access on the entire data pool via barcode hybridization, without need for amplification and without crosstalk with the physically isolated data-encoding DNA, followed by downstream selection that may be optical, physical, or biochemical. Each “unit of information” encoded in DNA we term a *file*, which includes both the DNA encoding the main data as well as any additional components used for addressing, storage, and retrieval. Each file contains a *file sequence*, consisting of the DNA encoding the main data, and *addressing barcodes*, or simply *barcodes*, which are additional short DNA sequences used to identify the file in solution using hybridization. We refer to a collection of files as a *data pool* or *database*, and the set of procedures for storing, retrieving, and reading out files is termed a *file system* (see **Supplementary Section S0** for a full list of terms).

As a proof-of-principle of our archival DNA file system, we encapsulated 20 image files, each composed of a ∼0.1 kilobyte image file encoded in a 3,000-base-pair plasmid, within monodisperse, 6-µm silica particles that were chemically surface-labeled using up to three 25-mer single-stranded DNA (ssDNA) oligonucleotide barcodes chosen from a library of 240,000 orthogonal primers, which allows for identification of up to ∼10^15^ possible distinct files using only three unique barcodes per file^19^ (**Fig. 1**). While we chose plasmids to encode DNA data in order to produce microgram quantities of DNA memory at low cost and to facilitate a renewable, closed-cycle write-store-access-read system using bacterial DNA data encoding and expression^20-22^, our file system is equally applicable to single-stranded DNA oligos produced using solid-phase chemical synthesis^2,7,8,10-13,17^ or gene-length oligos produced enzymatically^23-26^, and larger file sizes on the megabyte to gigabyte scale. And while only twenty icon-resolution images were chosen as our image database, representing diverse subject matter including animals, plants, transportation, and buildings (**Supplementary Fig.1**), our file system equally applies to thousands, billions, or larger sets of images, limited only by the cost of DNA synthesis, rather than any intrinsic property of our file system itself (**Supplementary Fig.1**).

**Figure 1.**
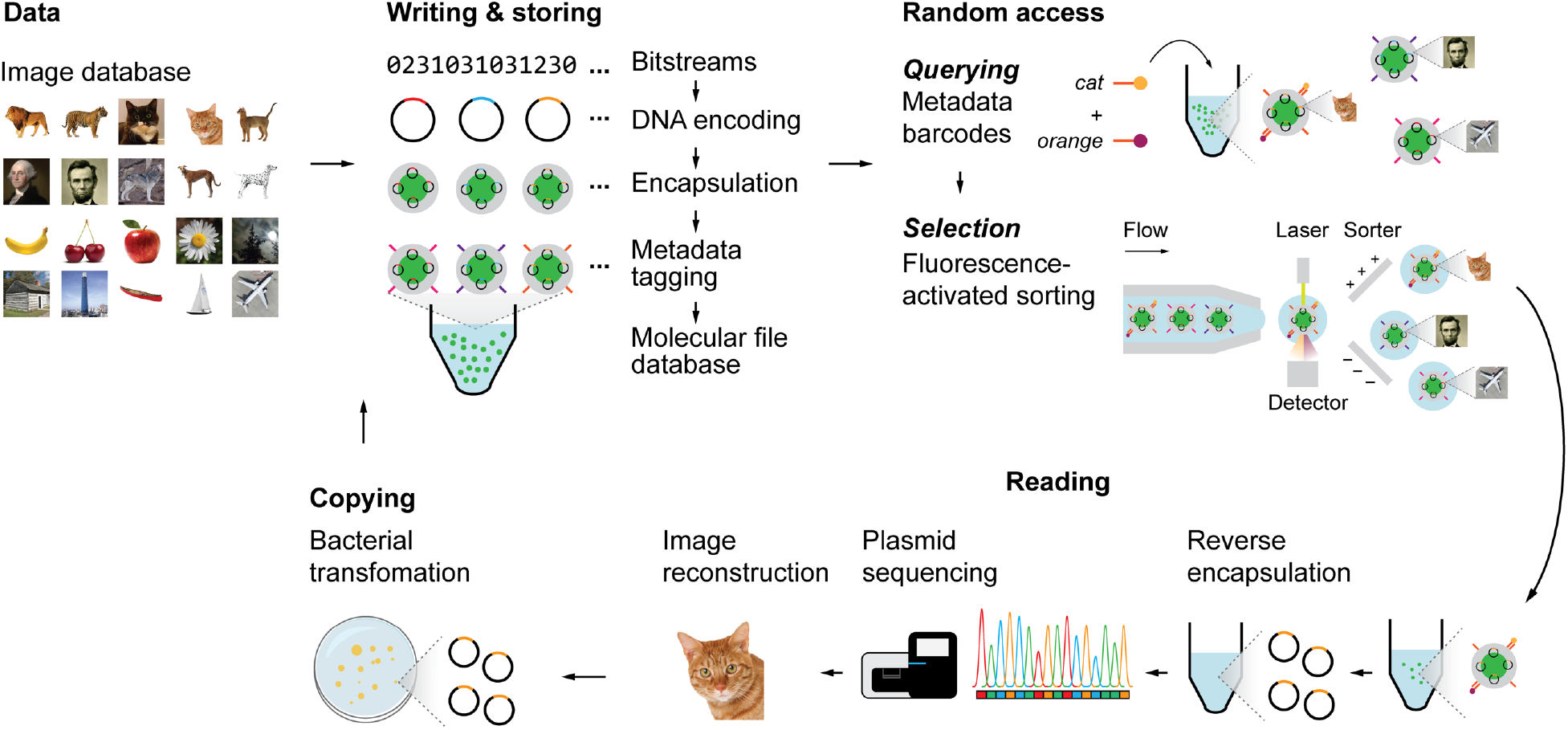
Write-access-read cycle for a content-addressable molecular file system. Colored images were converted into 26 × 26-pixel, black-and-white icon bitmaps. The black-and-white images were then converted into DNA sequences using ternary encoding scheme ^8^. The DNA sequences that encoded the images (file sequences) were inserted into a pUC19 plasmid vector and encapsulated into silica particles using sol-gel chemistry. Silica capsules were then addressed with content barcodes using orthogonal 25-mer single-stranded DNA strands, which were the final forms of the files. Files were pooled to form the molecular file database. To query a file or several files, fluorescently-labelled 15-mer ssDNA probes that are complementary to file barcodes were added to the data pool. Particles were then sorted with fluorescence-activated sorting (FAS) using two to four fluorescence channels simultaneously. Addition of a chemical etching reagent into the sorted populations released the encapsulated DNA plasmid. Sequences for the encoded images were validated using Sanger sequencing or Illumina MiniSeq. Because plasmids were used to encode information, re-transformation of the released plasmids into bacteria to replenish the molecular file database thereby closed the write-access-read cycle.

Fluorescence-activated sorting (FAS) was used to select target subsets of the complete data pool by first annealing fluorescent oligonucleotide probes that are complementary to the barcodes used to address the database^27^, enabling direct retrieval of specific, individual files from a pool of (10^6^)^*N*^ total files, where *N* is the number of fluorescence channels employed, without amplification required for PCR-based approaches, or loss of nucleotides available for data encoding. Further, our system enables direct, complex Boolean AND, OR, NOT logic to select random subsets of files with combinations of distinct barcodes to query the data pool, similar to conventional Boolean logic applied in text and file searches on solid-state silicon devices. And because physical encapsulation separates file sequences from external barcodes that are used to describe the encapsulated information, our file system offers long-term environmental protection of encoded file sequences via silica encapsulation for permanent archival storage^10,28,29^, where external barcodes may be renewed periodically, further protected with secondary encapsulation, or replaced for more sophisticated file operations involving re-labeling of data pools. Taken together, our strategy presents a practical and scalable archival molecular file storage system with random access capability that applies to the exabyte-to-yottabyte scales, limited only by the current cost of DNA synthesis.

### File Synthesis

Digital information in the form of 20 icon-resolution images was stored in a data pool, with each image encoded into DNA and synthesized on a plasmid. We selected images of broad diversity, representative of distinct and shared subject categories, which included several domestic and wild cats and dogs, US presidents, and several human-made objects such as an airplane, boats, and buildings (**Fig. 1** and **Supplementary Fig.1**). To implement this image database, the images were substituted with black-and-white, 26 × 26-pixel images to minimize synthesis costs, compressed using run-length encoding, and converted to DNA (**Supplementary Fig. 1,2**). Following synthesis, bacterial amplification, and sequencing validation (**Supplementary Fig.3**), each plasmid DNA was separately encapsulated into silica particles containing a fluorescein dye core and a positively charged surface^28,29^. Because the negatively charged phosphate groups of the DNA interact with positively charged silica particles, plasmid DNA condensed on the silica surface, after which N-[3-(trimethoxysilyl)propyl]-N,N,N-trimethylammonium chloride (TMAPS) was co-condensed with tetraethoxysilane to form an encapsulation shell after four days of incubation at room-temperature^10,29^ (**Fig. 2a**) to form discrete silica capsules containing the file sequence that encodes for the image file. Quantitative PCR (qPCR) of the reaction supernatant after encapsulation (**Supplementary Fig.4**) showed full encapsulation of plasmids without residual DNA in solution. To investigate the fraction of capsules that contained plasmid DNA, we compared the fluorescence intensity of the intercalating dye TO-PRO when added pre-versus post-encapsulation (**Supplementary Fig.2**). All capsules synthesized in the presence of both DNA and TO-PRO showed a distinct fluorescence signal, consistent with the presence of plasmid DNA in the majority of capsules, compared with a silica particle negative control that contained no DNA. In order to test whether plasmid DNA was fully encapsulated versus partially exposed at the surface of capsules, capsules were also stained separately with TO-PRO post-encapsulation (**Fig. 2b**). Using qPCR, we estimated 10^6^ plasmids per capsule assuming quantitative recovery of DNA post-encapsulation (**Supplementary Fig.5**). Because encapsulation of the DNA file sequence relies only on electrostatic interactions between positively-charged silica and the phosphate backbone of DNA, our approach can equally encapsulate any molecular weight of DNA molecule applicable to MB and larger file sizes, as demonstrated previously^29^, and is compatible with alternative DNA file compositions such as 100-200-mer oligonucleotides that are commonly used ^2,7,8,12,13,17^.

**Figure 2.**
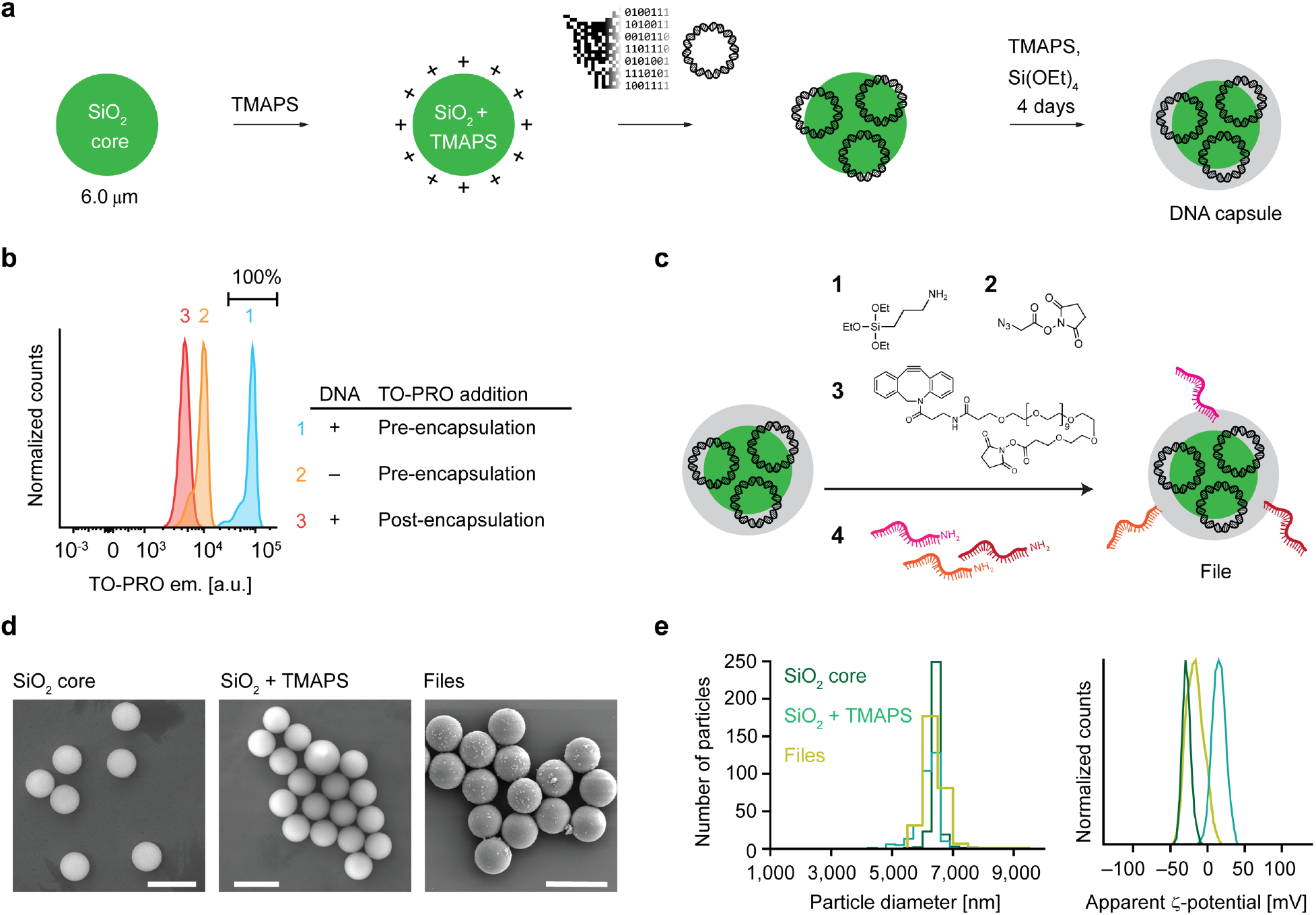
Encapsulation of DNA plasmids into silica and surface barcoding. **a**, Workflow of silica encapsulation ^29^. **b**, Raw fluorescence data from FAS experiments to detect DNA staining of TO-PRO during or after encapsulation. **c**, Functionalization of encapsulated DNA particles. **d**, Scanning electron microscopy images of bare silica particles, silica particles functionalized with TMAPS, and the file. **e**, Distribution of particle sizes determined from microscopy data (left) and zeta potential analyses of silica particles and files.

Next, we chemically attached unique content addresses on the surfaces of silica capsules using orthogonal 25-mer ssDNA barcodes (**Supplementary Fig. 6**) describing selected features of the underlying image for file selection. For example, the image of an orange tabby house cat (**Supplementary Fig.1**) was described with *cat, orange*, and *domestic*, whereas the image of a tiger was described with *cat, orange*, and *wild* (**Supplementary Fig.1** and **Supplementary Table 2**). To attach the barcodes, we activated the surface of the silica capsules through a series of chemical steps. Condensation of 3-aminopropyltriethoxysilane with the hydroxy-terminated surface of the encapsulated plasmid DNA provided a primary amine chemical handle that supported further conjugation reactions (**Fig. 2c**). We modified the amino-modified surface of the silica capsules with 2-azidoacetic acid N-hydroxysuccinimide (NHS) ester followed by an oligo(ethylene glycol) that contained two chemically orthogonal functional groups: the dibenzocyclooctyne functional group reacted with the surface-attached azide through strain-promoted azide-alkyne cycloaddition while the NHS ester functional group was available for subsequent conjugation with a primary amine. Each of the associated barcodes contained a 5’-amino modification that could react with the NHS-ester groups on the surface of the silica capsules, thereby producing the complete form of our file. Notably, the sizes of bare, hydroxy-terminated silica particles representing capsules without barcodes were comparable with complete files consisting of capsules with barcodes attached, confirmed using scanning electron microscopy (**Fig. 2d** and **2e**, left). These results were anticipated given that the encapsulation thickness was only on the order of 10 nm^29^ and that additional steps to attach functional groups minimally increases the capsule diameter. We also observed systematic shifts in the surface charge of the silica particles as different functional groups were introduced onto their surfaces (**Fig. 2e**). Using hybridization assays with fluorescently-labelled probes^30-32^, we estimated the number of barcodes available for hybridization on each file to be on the order of 10^8^ (**Supplementary Fig. 7**). Following synthesis, files were pooled and stored together for subsequent retrieval. Illumina MiSeq was used to read each file sequence and reconstruct the encoded image following selection and de-encapsulation, in order to validate the complete process of image file encoding, encapsulation, barcoding, selection, de-encapsulation, sequencing, and image file reconstruction (**Supplementary Figs. 9, 10**).

### File Selection

Following file synthesis and pooling, we used FAS to select specific targeted files from the complete data pool through the reversible binding of fluorescent probe molecules to the file barcodes (**Supplementary Fig. 6**). All files contained a fluorescent dye, fluorescein, in their core as a marker to distinguish files from other particulates such as spurious silica particles that nucleated in the absence of a core or insoluble salts that may have formed during the sorting process. Each detected fluorescein event was therefore interpreted to indicate the presence of a single file during FAS (**Supplementary Fig. 11**). To apply a query such as *flying* to the image database, the corresponding fluorescently labeled ssDNA probe was added, which hybridized to the complementary barcode displayed externally on the surface of a silica capsule for FAS selection (**Fig. 3a**).

**Figure 3.**
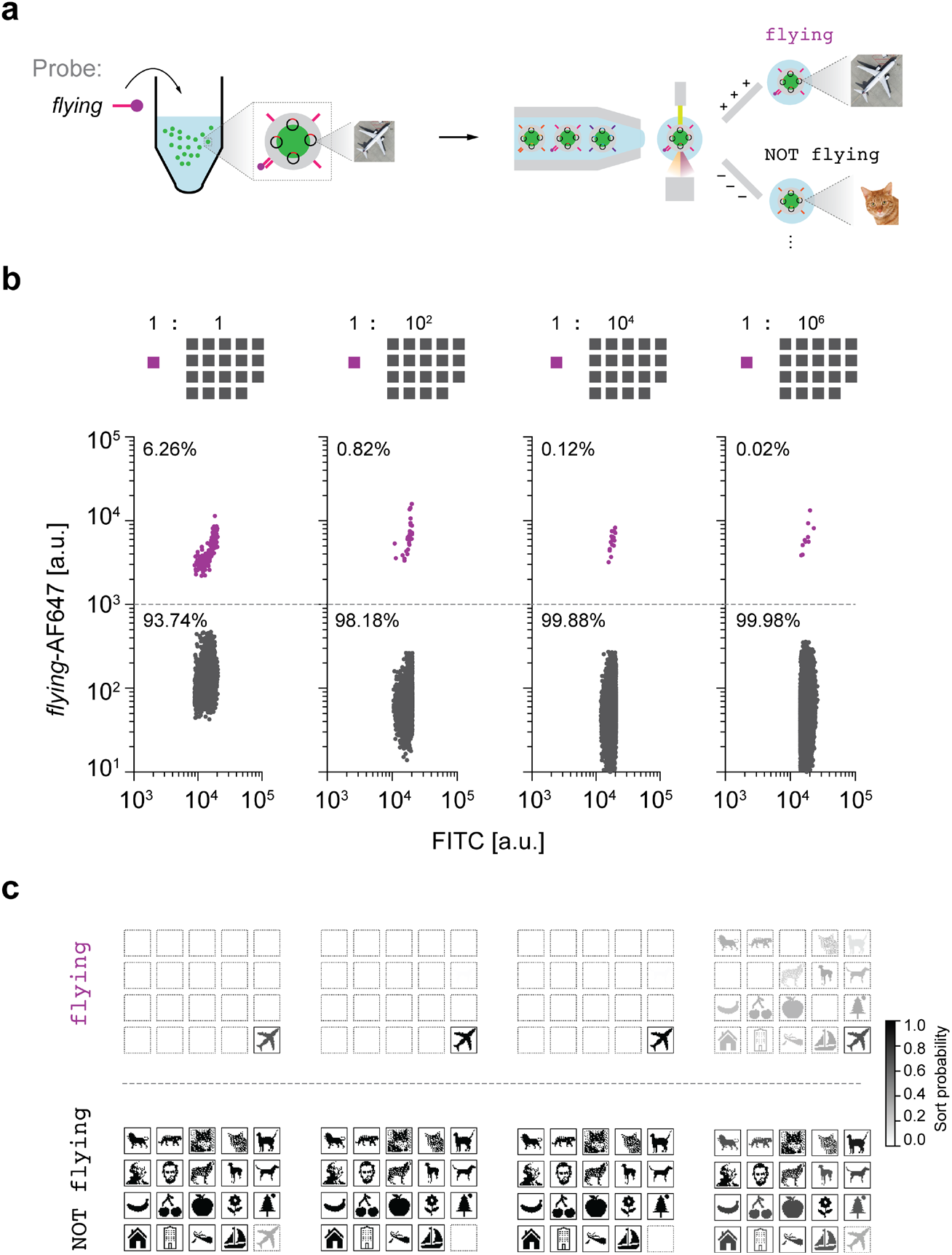
Single-barcode sorting. **a**, Schematic diagram of file sorting using FAS. **b**, Sorting of *Airplane* from varying relative abundance of the other nineteen files as background. Percentages represent the numbers of particles that were sorted in the gate. Colored traces in each of the sorting plots indicate the target population. **c**, Sequencing validation using Illumina MiniSeq. Sort probability is the probability that a file is sorted into one gated population over the other gated populations. Boxes with solid outlines indicate files that should be sorted into the specified gate. Other files have dashed outlines.

We subjected the entire data pool to a series of experiments to test selection sensitivity of target subsets using distinct queries. First, we evaluated single-barcode selection of an individual file, specifically *Airplane*, out of a pool of varying concentrations of the nineteen other files as background (**Fig. 3b**). To select the *Airplane* file, we hybridized an AFDye 647-labelled ssDNA probe that is complementary to the barcode *flying*, which is unique to *Airplane*. We were able to detect and select the desired *Airplane* file through FAS even at a relative abundance of 10^−6^ compared with each other file (**Fig. 3c**). While comparable in sensitivity to a nested PCR barcoding data indexing approach^17^, unlike PCR that requires 20–30 of rounds of heating and cooling to selectively amplify the selected sequence, our approach selects files directly without need for thermal cycling and amplification. This strategy also applies to gating of *N* barcodes simultaneously in parallel optical channels, which offers file selection sensitivity of 1 in 10^6*N*^ total files, where common commercial FAS systems offer up to *N* = 17 channels^33,34^. For example, comparison of the retrieved sequences between the flying gate and the NOT flying gate after chemical release of the file sequences from silica encapsulation revealed that 60–95% of the *Airplane* files were sorted into the flying gate (**Supplementary Figs. 18–21**), where we note that any sort probability above 50% indicates enrichment of *Airplane* within the correct population subset (flying) relative to the incorrect subset (NOT flying), while a sort probability of 100% would indicate ideal performance. Besides single file selection, our approach allows for repeated rounds of FAS selection, as well as Boolean logic, described below.

### Boolean Search

Beyond direct selection of 1 in 10^6*N*^ individual random files directly, without thermal cycling or loss of fidelity due to primer crosstalk, our system offers the ability to apply Boolean logic to select random file subsets from the data pool. AND, OR, and NOT logical operations were applied by first adding to the data pool fluorescently labeled ssDNA probes that were complementary to the barcodes (**Fig. 4**, left). This hybridization reaction was used to distinguish one or several files in the data pool, which were then sorted using FAS. We used two to four fluorescence channels simultaneously to create the FAS gates that executed the target Boolean logic queries (**Fig. 4**, middle). To demonstrate a NOT query, we added to the data pool an AFDye 647-labelled ssDNA probe that hybridized to files that contained the *cat* barcode. Files that did not show AFDye 647 signal were sorted into the NOT cat subset (**Fig. 4a**). An example of an OR gate was applied to the data pool by simultaneously adding *dog* and *building* probes that both had the TAMRA label (**Fig. 4b**). All files that showed TAMRA signal were sorted into the dog OR building subset by the FAS. Finally, an example of an AND gate was achieved by adding *fruit* and *yellow* probes that were labelled with AFDye 647 and TAMRA, respectively. Files showing signal for both AFDye 647 and TAMRA were sorted into the fruit AND yellow subset in the FAS (**Fig. 4c**). For each example query, we validated our sorting experiments by releasing the file sequence from silica encapsulation and sequencing the released DNA with Illumina MiniSeq (**Fig. 4**, right). Sort probabilities of each file for each search query are shown in **Supplementary Figs. S22–S24**.

**Figure 4.**
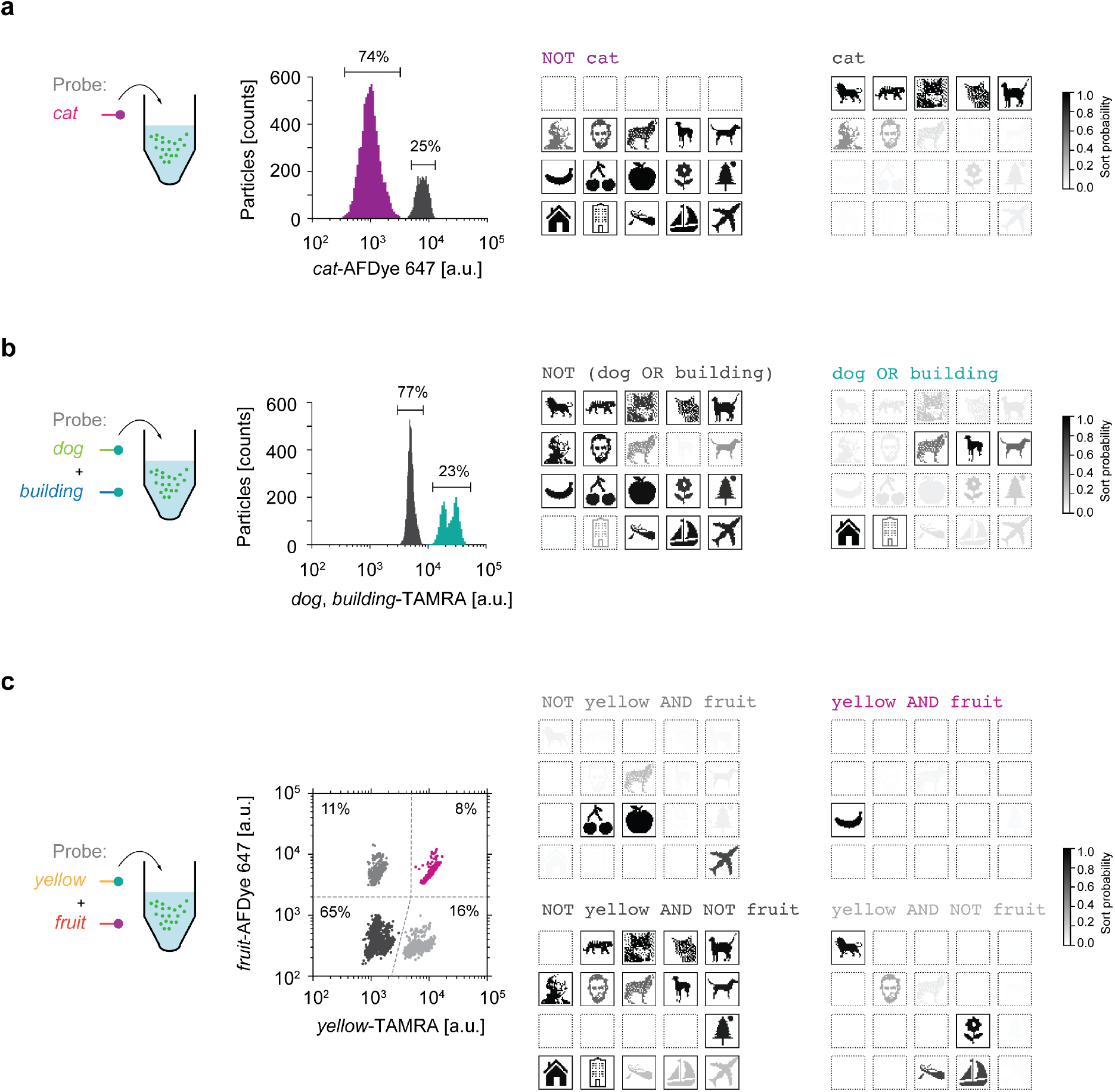
Fundamental Boolean logic gates. **a**, NOT cat selection. Raw fluorescence trace from the FAS system (left) plotted on a 1D sorting plot showing the percent of particles that were sorted in each gate. Sequencing using Illumina MiniSeq tested selection specificity (right). **b**, dog OR building selection. Raw fluorescence trace from the FAS system (left) plotted on a 1D sorting plot showing the percent of particles that were sorted in each gate. Sequencing using Illumina MiniSeq evaluated sorting using the OR gate (right). **c**, A 2D sorting plot to perform a yellow AND fruit gate. Percentages in each quadrant show the percentages of particles that were sorted in each gate. Colored traces in all of the sorting plots indicate the target populations. Sort probability is the probability that a file is sorted into one gated population versus the other gated populations. Boxes with solid outlines indicate files that were intended to sort into the specified gate. Other files have dashed outlines.

The preceding demonstrations of Boolean logic gates enable file sorting with varying specificity of selection criteria for the retrieval of different subsets of the data pool. FAS can also be used to create multiple gating conditions simultaneously, thereby increasing the complexity of target file selection operations, as noted above. To demonstrate increasingly complex Boolean search queries, we selected the file containing the image of Abraham Lincoln from the data pool, which included images of two presidents, George Washington and Abraham Lincoln. The *president* ssDNA probe, fluorescently labeled with TAMRA, selected both *Lincoln* and *Washington* files from the data pool. The simultaneous addition of the *18*^*th*^ *century* ssDNA probe, fluorescently labeled with AFDye 647 (**Fig. 5a**, left), discriminated *Washington*, which contained the *18*^*th*^ *century* barcode, from the *Lincoln* file (**Fig. 5a**, middle). The combination of these two ssDNA probes permitted the complex search query president AND (NOT 18^th^ century). Sequencing analysis of the gated populations after reverse encapsulation validated that the sorted populations matched search queries for president AND (NOT 18^th^ century), president AND 18^th^ century, and NOT president (**Fig. 5a**, right; **Supplementary Fig. 25**).

**Figure 5.**
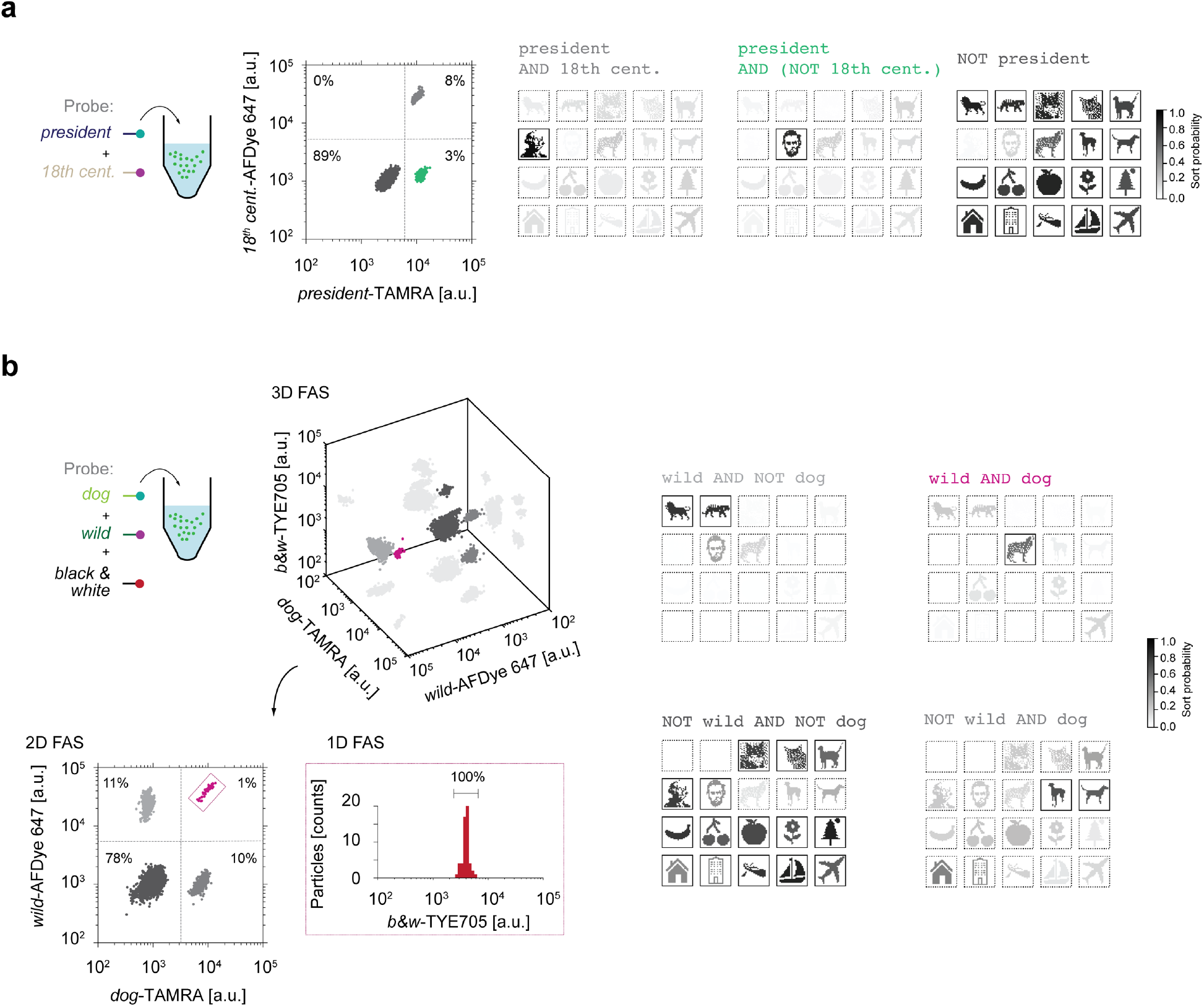
Arbitrary logic searching. **a**, president AND (NOT 18^th^ century) sorting. A 2D sorting plot (middle) was used to sort *Lincoln* by selecting a population that has high TAMRA fluorescence but low AFDye 647 fluorescence. Sequencing using MiniSeq offered quantitative evaluation of the sorted populations. **b**, Multiple fluorescence channels were projected into a 3D FAS plot (left and top). There are three possible 2D plots that can be used for sorting. To select the *Wolf* image using the query wild AND dog, a 2D plot of *wild* versus *dog* was first selected and then populations selected using quadrant gates (left and bottom). One of the quadrants were then selected where the *Wolf* image should belong based on the wild AND dog query in order to test whether only a single population was present in the TYE705 fluorescence channel. Sequencing quantified the sorted populations (right) using Illumina MiniSeq. Sort probability is the probability that a file was sorted into one gated population over the other gated populations. Boxes with solid outlines indicate files that would ideally be sorted into the specified gate. Other files have dashed outlines.

To demonstrate the feasibility of performing Boolean search using more than three fluorescence channels for sorting, we also selected the *Wolf* file from the data pool using the query dog AND wild, and used the *black & white* probe to validate the selected file (**Fig. 5b**, left). Because conventional FAS software is only capable of sorting using 1D and 2D gates, we first selected one out of the three possible 2D plots (**Fig. 5b**, left and bottom): *dog*-TAMRA against *wild*-AFDye 647. We examined the *black & white*-TYE705 channel on members of the dog AND wild subset (**Fig. 5b**, left and bottom). Release of the encapsulated file sequence and subsequent sequencing of each gated population from the *dog* versus *wild* 2D plot validated sorting (**Fig. 5b**, right; **Supplementary Fig. 26**).

In contrast to single-stranded DNA oligos, our use of plasmids as a substrate for encoding information offered the ability to restore files into the data pool after retrieval. In cases where single images were sorted (**Figs. 4c, 5a, b**), we were able to transform competent bacteria from each search query that resulted in a single file (**Supplementary Fig. 27**). Amplified material was pure and ready for re-encapsulation into silica particles, which could be re-introduced directly back into the data pool. Importantly, our molecular file system and file selection process thereby represents a complete write-store-access-read cycle that in principle may be applied to exabyte and larger-scale datasets, with periodic renewal of single-stranded DNA barcodes and bacterial replication of DNA data following reading^20-22^. While sort probabilities were typically below the optimal 100% targeted for a specific file or file subset query, future work may characterize sources of error that could be due to sample contamination or random FAS errors. The latter type of error can be mitigated through repeated cycles of file selection in series. Our technical approach differs significantly from approaches that rely on selective PCR amplification for selection^9,11,13,17,18^, in which repeated amplifications may reduce fidelity of file selection.

## Discussion & Outlook

We introduce a scalable, non-destructive, random access molecular file system for the direct access of arbitrary files and file-subsets from an archival DNA data store. The introduction of our file system overcomes former limitations of indirect, PCR-based file systems for the practical implementation of archival DNA memory systems. This advance now leaves the high cost of DNA synthesis compared with alternative memory storage media as the primary remaining rate-limiting step for translation of this technology. While the overall data density of our file system is considerably lower than the theoretical limit of DNA data density due to the encapsulation of DNA files in silica particles, the physical size of exabyte-scale DNA data stored in our system is still orders of magnitude smaller than conventional archival file storage systems. For example, assuming 2 bits per base, 10^−21^ grams per base, and a density of double-stranded DNA of 1.7 grams per cubic centimeter^4^, PCR-based random access approaches have a theoretical volumetric density limit of 10^27^ bytes per m^3^, compared with our approach of 10^24^ bytes per m^3^ that is 10^3^-fold lower (**Supplementary Section S6**). However, PCR suffers from numerous issues such as enzyme cost, requirement of numerous heating and cooling cycles, and potential crosstalk between file sequences and barcodes^17,18^, which requires spatial segregation of file sequences in electrowetting devices^18^ that reduced data density to ∼10^20^ bytes per m^3^, seven orders of magnitude below the theoretical limit for dry DNA (**Supplementary Section S6**).

In the current implementation of our file system, each file capsule contained 10^6^ DNA plasmids, which could instead store multiple unique file-encoding plasmids or file fragments to increase data density to gigabyte-sized files per capsule, with an overall data density of 10^24^ bytes per m^3^ (**Supplementary Section S6**), which is only three orders of magnitude lower than the theoretical data density limit of dry DNA, and four orders of magnitude higher than published approaches to storing and accessing DNA data with spatial segregation^18^. And equally important to data density per se is the physical size required to store an exabyte-or larger-scale DNA data pool. Using our approach, 10^9^ gigabyte-sized files would still only require 0.2 cm^3^ of total dry volume, without any need for physically separated data pools. Notwithstanding, further increases in data density could be achieved by using nanoparticles ∼100–200 nm in diameter to encode files^10,28,29^ sorted with higher sensitivity FAS systems^35,36^, or multiple layers of encapsulated DNA^37^.

In addition to data pool size and density, another crucial operating feature is the latency or time associated with DNA file retrieval. Because FAS scales linearly with the size of the data pool, retrieval time may still be limiting in an exabyte-scale data pool, even assuming gigabyte-sized files. To further reduce file selection time, future file system implementations may leverage parallel microfluidics-based optical sorting procedures, brighter fluorescent probes to increase selection throughput, alternative barcode implementations^38-42^, or physical sorting strategies such as direct biochemical pulldown^17,43,44^, such as recently implemented using direct magnetic extraction of files labelled with biochemical affinity tags^17^. Additional latency due to chemical deprotection of DNA from silica encapsulation renders our file system ideally suited to long-term, archival DNA storage at the exabyte-to-yottabyte scales.

Indeed, because we view our scalable file system as an alternative to tape-based, ‘cold’ archival data storage systems rather than flash or other ‘hot’ memory, for which latency times may be tolerated on the time frame of several days to weeks, the foregoing latency limitations are of minimal importance compared with the transformative capability offered by our system to store exabyte-to-yottabyte-scale datasets with direct retrieval of arbitrary, random file subsets. Example applications include the retrieval of specific images from archival databases of astronomical image databases^45^, high-energy physics datasets^46^, or high-resolution deep ocean floor mapping^47^.

Finally, because our system is not limited to synthetic DNA, it applies equally to long-term archival storage of bacterial, human, and other genomes for archival sample preservation and retrieval^23,48^, forensic analysis, and retrospective analysis of pandemic outbreaks, as explored in accompanying work^49^. Our demonstrated file system enables complex file search operations on underlying molecular data pools, moving us closer to realizing an economically viable, functional massive molecular file and operating system^27,50,51^.

## Acknowledgments

We gratefully acknowledge fruitful discussions with Charles Leiserson and Tao B. Schardl on the scalability and generalizability of our barcoding approach. We thank Glenn Paradis, Michael Jennings, and Michele Griffin of the Flow Cytometry Core at the Koch Institute in MIT and Patricia Rogers of the Flow Cytometry Facility at the Broad Institute of Harvard and MIT for assistance and fruitful discussions in developing the flow cytometry workflow. We also thank David Mankus of the Nanotechnology Materials Core Facility at the Koch Institute in MIT for assistance in the imaging of the particles using the scanning electron microscope and Alla Leshinsky of the Biopolymer and Proteomics Core at the Koch Institute at MIT for assistance in mass spectrometry characterization.

## Author contributions

J.L.B., T.R.S., and M.B. designed the file labeling and selection scheme. J.L.B, T.R.S., and C.M.A. implemented the file selection scheme using FAS. J.B. and T.R.S. developed the encoding scheme and metadata tagging of the images to DNA. T.R.S. designed the plasmid for encoding imaging. H.H. and T.R.S. performed the cloning, transformation, and purification of the plasmids. J.L.B. synthesized and purified all the TAMRA and AFDye 647-labelled DNA oligonucleotides. J.L.B. characterized the particles. J.L.B. developed the synthetic route to attach DNA barcodes on the surface of the particles. J.L.B. performed the encapsulation, barcoding, sorting, reverse encapsulation of the particles after sorting, and desalting. T.R.S., H.H., and M.R. performed the sequencing. J.B. performed computational validation of the orthogonality of barcode sequences and J.L.B. performed the experimental validation of the orthogonality of barcode and probe sequences. J.B. developed the computational workflow to analyze the sequencing data, including statistical analyses. M.B. conceived of the file system and supervised the entire project. P.C.B. supervised the FAS selection and supervised the sequencing workflow. All authors analyzed the data and equally contributed to the writing of the manuscript.

## Materials and correspondence

Gene sequences and plasmid maps are available from AddGene (https://www.addgene.org/depositing/77231/). Software for sequence encoding and decoding is publicly available on GitHub (https://github.com/lcbb/DNA-Memory-Blocks/). All the data files used to generate the plots in this manuscript are available from M.B. upon request.

### Online content

Any methods, additional references, and supplementary information are available at https://doi.org/10.10XX/XXXXX.

