## Supplementary Information for "Random access DNA memory in a scalable, archival file storage system"

### S0. Glossary of terms

| Terms | Definition |
|---|---|
| file | The most basic unit of a file system, consisting of the DNA encoding the main data (the file sequence), addressing barcodes, and any other components necessary for storage and/or retrieval. In our particular file system, each file is a silica particle that contains the file sequence and displays on its surface DNA barcodes describing features of the data.<br>File names in this paper are italicized and the first letter is capitalized.<br>Example: *Cat2* |
| file sequence | A DNA sequence that encodes the data in the file. |
| barcode/addressing barcode | A 25-mer single-stranded DNA sequence that is used to describe a single feature of the data.<br>Barcode names in this paper are italicized and lower case.<br>Example: *cat* |
| capsule | A silica particle that contains encapsulated DNA; a capsule with barcodes added to the surface constitutes a file. |
| data pool/database | A collection of files. |
| probe | A fluorescently-labelled 15-mer single-stranded DNA sequence that is complementary to a barcode.<br>Probe names in this paper are italicized and lower case.<br>Example: *cat* |
| query | A request for a particular subset of a database.<br>Queries in this paper are written in monospaced Courier font.<br>Example: `cat AND (NOT wild)` |
| file system | A system for storing and organizing data. |

### S1. Materials and methods

**General materials.** All DNA oligonucleotides (oligos), including 5'-amino-modified DNA oligos and TYE705-modified oligos, were purchased from Integrated DNA Technologies (IDT; Coralville, IA) with standard desalting as purification method and were received as dry pellets. Upon receipt of the DNA oligonucleotides, the pellets were dissolved in 1× PBS (catalog number: 79378) from Millipore Sigma (Milwaukee, WI) and kept at 4 °C until further use.

Fluorescein-core silica particles with 5-µm diameter and hydroxyl-terminated surfaces (catalog number: DNG-L034) were obtained from Creative Diagnostics (Shirley, NY). N-hydroxysuccinimide (NHS) ester of TAMRA (catalog number: 1255-25) and AFDye 647 (catalog number: 1121-5) was purchased from Fluoroprobes (Scottsdale, AZ). DBCO-PEG13-NHS ester (catalog number: 1015), and azidoacetic acid NHS ester (catalog number: 1070) were purchased from Click Chemistry Tools (Scottsdale, AZ). TEOS (catalog number: 131903), and APTS (catalog number: 440140) were purchased from Millipore Sigma (Milwaukee, WI). Ammonia in water (28% $NH_3$; catalog number: 338818) was purchased from Millipore Sigma and stored at 4 °C. TMAPS (50% in methanol) was obtained through Alfa Aesar (Haverhill, MA) and stored at 4 °C. Anhydrous organic solvents, dimethyl sulfoxide (DMSO; catalog number: 276855), *N*-methyl-2-pyrrolidone (catalog number: 270458), isopropanol (catalog number: 278475), and ethanol (catalog number: 459836), were purchased from Millipore Sigma (Milwaukee, WI). Activated molecular sieves (3 Å; Millipore Sigma; catalog number: 208574) were added to anhydrous DMSO and DMF upon opening.

Gene sequences and all oligonucleotides were purchased as specified from IDT. Gene sequences and plasmid maps are available from AddGene (https://www.addgene.org/depositing/77231/). Plasmids were verified by IDT using Illumina MiSeq and by Sanger Sequencing by GeneWiz (South Plainfield, NJ) and Illumina MiniSeq. SeaKem agarose was purchased from Lonza (Basel, Switzerland). SybrSafe was purchased from ThermoFisher (Waltham, MA). Luna universal qPCR Master Mix was purchased from New England Biolabs (NEB, Ipswitch, MA). Qiagen (Venlo, Netherlands) HiSpeed Plasmid Midi and Maxi kits were used for plasmid purification after amplification in 100 mL of LB (Sigma; St. Louis, MO) of transformed DH5α *Escherichia coli* cells (NEB). Illustra S-200 HR spin columns (GE Healthcare; Boston, MA) were used for buffer-exchanged salt removal. Qiagen Spin Mini kits were used for small-scale cleanup and concentration of DNAs.

**Characterization of materials.** Surface zeta-potentials were measured using a Malvern Zetasizer Nano ZSP. All samples for surface zeta-potentials were prepared and measured in a standard fluorescence quartz cuvette (catalog number: 3-Q-10) from Starna Cells, Inc. (Atascadero, CA) at a concentration of 0.1 mg $mL^{-1}$ with a volume of 700 µL. A universal 'dip' probe (catalog number: ZEN1002) from Malvern was used to measure zeta potential of particles. Scanning electron microscopy of the particles were performed using a Zeiss Gemini 2 Field Emission Scanning Electron Microscope. Samples were mounted on silicon substrates or glass. For glass-mounted samples, ~5 nm of gold was sputter-coated to make the samples conductive.

### S2. Core memory plasmid sequences

Twenty 26 × 26-pixel, black-and-white icon bitmaps were generated as representations of 20 high-resolution color images (**Supplementary Fig. 1**) encompassing a broad range of subject matters. Each black-and-white icon was converted to a length-676 bitstring in a column-first order, with each black or white pixel encoded as a 0 or 1, respectively. This bitstring was compressed via run length encoding, replacing each stretch of consecutive 0s or 1s with a 2-tuple (value, length) to generate a list of 2-tuples describing the entire bitstring. The maximum length is 15; runs longer than 15 bits are encoded as multiple consecutive runs of the same value. This run length encoding was converted to a sequence of base-4 digits as follows:

1) Begin with an empty string, and set the current run to the first run of the list.
2) Append the value of the current run (0 or 1).
3) Using 2 base-4 digits, append the length of the current run.
4) If the length of this run was 15, encode the next run starting with Step (2). Otherwise, encode starting at Step (3). If no runs remain, the process is complete.

This process produces a quaternary string describing the entire run length encoding of the image. To avoid homopolymer runs and repeated subsequences in the final nucleotide sequence, each digit is offset by a random number generated from a linear congruential random number generator (LCG) beginning with a random seed (i.e. the $i^{th}$ value generated by the LCG is added, modulo 4, to the $i^{th}$ base-4 digit of the quaternary string). We used an LCG of multiplier 5, modulus $2^{31}$-1, and increment 0, although any LCG parameters with period longer than the length of the sequence would be suitable.

The final "randomized" quaternary string is converted to nucleotides by a direct mapping (0 = G, 1 = A, 2 = T, 3 = C). The number used to seed the LCG is prepended to this sequence by converting it into a ternary string of length 20, whose digits are encoded in nucleotides via a base transition table, as done previously by Goldman et al. [1]: (0 = GA, AT, TC, CG; 1 = GT, AC, TG, CA; 2 = GC, AG, TA, CT). The first digit is encoded directly (0 = A, 1 = T, 2 = C).

This sequence was modified with additional flanking sequences added to the beginning and end. A 64-bit wavelet hash of each bitmap was calculated using the whash function provided by the ImageHash Python package, available through the Python Package Index (https://pypi.org/project/ImageHash/). The 64-bit hash was split into two 32-bit halves, each of which was represented in a length-24 ternary string that was subsequently converted to nucleotides through the same process as applied to the seed. The two 24-nt regions were appended to the beginning and end of the sequence (**Supplementary Table 1**, orange text). The sequence containing image hash, seed, and image encoding, was additionally flanked on the 5' and 3' ends by sequences (5′-CGTCGTCGTCCCCTCAAACT-3′ and 5′-GCTGAAAAGGTGGCATCAAT-3′, respectively) that allow amplification from a "master primer" pair that would amplify every sequence in the molecular plasmid database (**Supplementary Table 1**, purple text; **Supplementary Fig. 2**).

The final sequence was checked for problematic subsequences, specifically, GGGG, CCCC, AAAAA, TTTTT, and the restriction enzyme recognition sites GAATTC and CTGCAG. If any of these subsequences were found outside of expected occurrences in the constant flanking regions, the entire sequence was regenerated with a new random seed until no such subsequences appeared.

The generated sequences were cloned on a pUC19-based vector. The software for sequence encoding and decoding is publicly available on GitHub at https://github.com/lcbb/DNA-Memory-Blocks/ and the plasmids are publicly available from AddGene (https://www.addgene.org/depositing/77231/). Each master primer and hash barcode pairs were verified by PCR and agarose gel analysis and PCR bias was checked by qPCR (**Supplementary Fig. 3**).

Generated DNA sequences were ordered as custom genes synthesized and cloned by IDT into pUC19-based plasmids, with sequencing validation provided by IDT. Each clone was designed with flanking single EcoRI and PstI cut sites, sequences which had been designed against in the inserts, enabling future sub-cloning to alternative vectors using standard digestion and ligation. All plasmids were amplified in bacterial cultures and purified using Qiagen HiSpeed Midi or Maxi kits following the protocol provided by the manufacturer. Briefly, DH5α cells were made chemically competent by growing 100 mL of cells to 0.5 and 0.6 at $OD_{600}$. The cells were gently pelleted and incubated on ice in 100 mM $CaCl_2$ for 30 minutes. The pelleted cells were then brought up in 100 mM $CaCl_2$ with 10% glycerol. Transformations from each plasmid were accomplished by incubation of 20 µL chemically competent cells with 1 ng of plasmid DNA for 30 minutes on ice, followed by heating to 42 °C for 45 seconds, placing on ice for 2 minutes, and then addition of 1 mL of pre-warmed SOC media. The mixture was shaken for 1 hour at 250 rpm at 37 °C. A volume of 20 µL of transformed cells were streaked to single colonies on agar plates that were supplemented with 100 µg $mL^{-1}$ ampicillin. Single colonies were chosen for growth in 100 mL LB with 100 µg $mL^{-1}$ ampicillin overnight in disposable baffled flasks and then followed by pelleting cells from the culture. Pelleted cells were lysed and treated with Qiagen purification buffers. Plasmid DNA was then purified on the provided silica support matrix and eluted in 1 mL of TE buffer (10 mM Tris-HCl, 1 mM EDTA, pH 8.0). Plasmid preparations were verified by PCR, Sanger sequencing, and Illumina sequencing as part of the workflow.

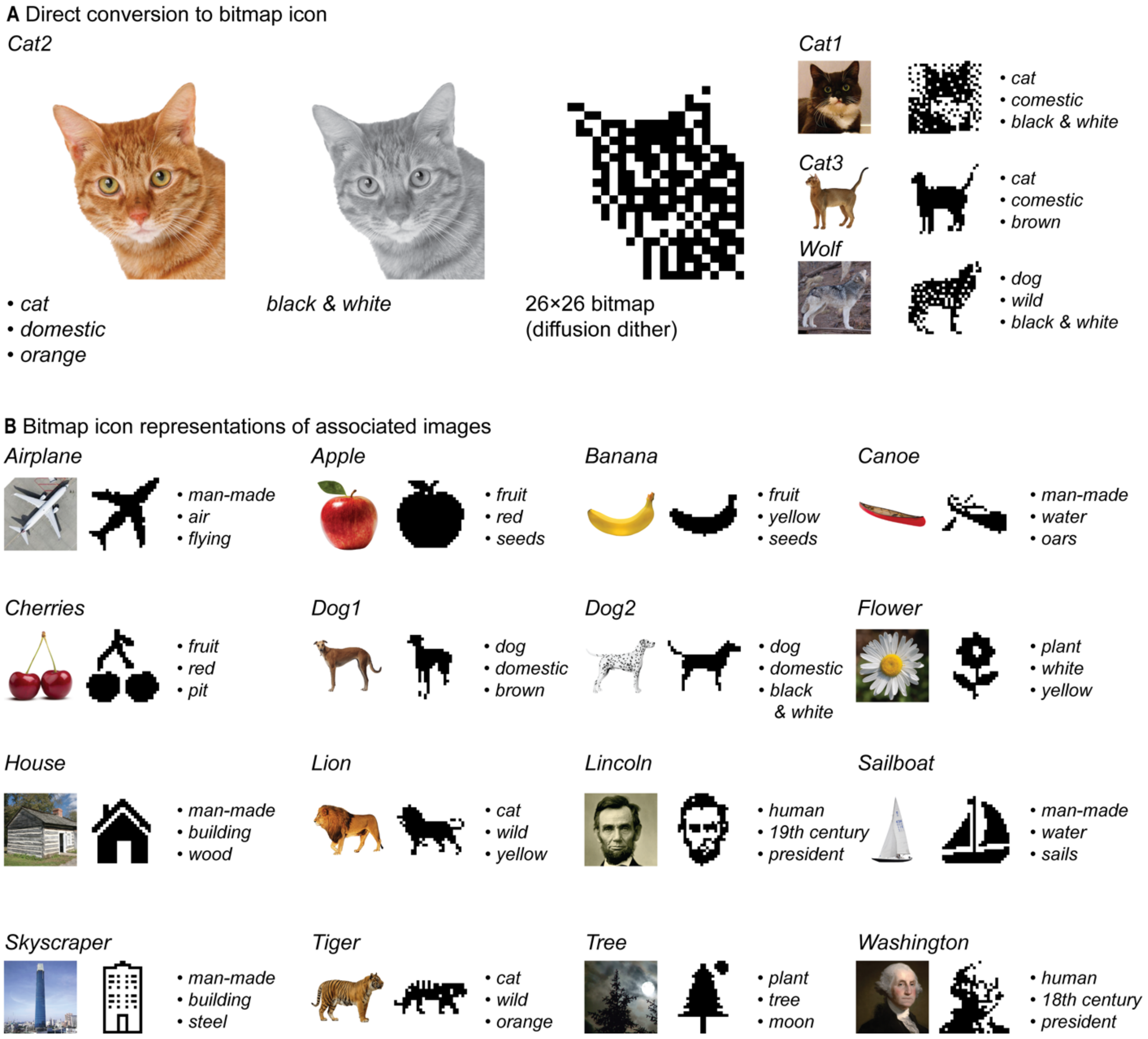

**Supplementary Figure 1. Image database with icon representations and content descriptors of the original images.** (**A**) Three pictures of cats and one picture of a wolf were directly converted to 26 × 26-pixel icons using Adobe Photoshop, by first changing the image to black and white, and then reducing the resolution to 26 pixels per inch with a 1-inch×1-inch image using diffusion dithering. (**B**) Sixteen other images were additionally selected of broad subject matter and icon images were chosen for image representation and reduced to 26 × 26-pixel sizes. Icon images were used to reduce sequence size and therefore cost of synthesis.

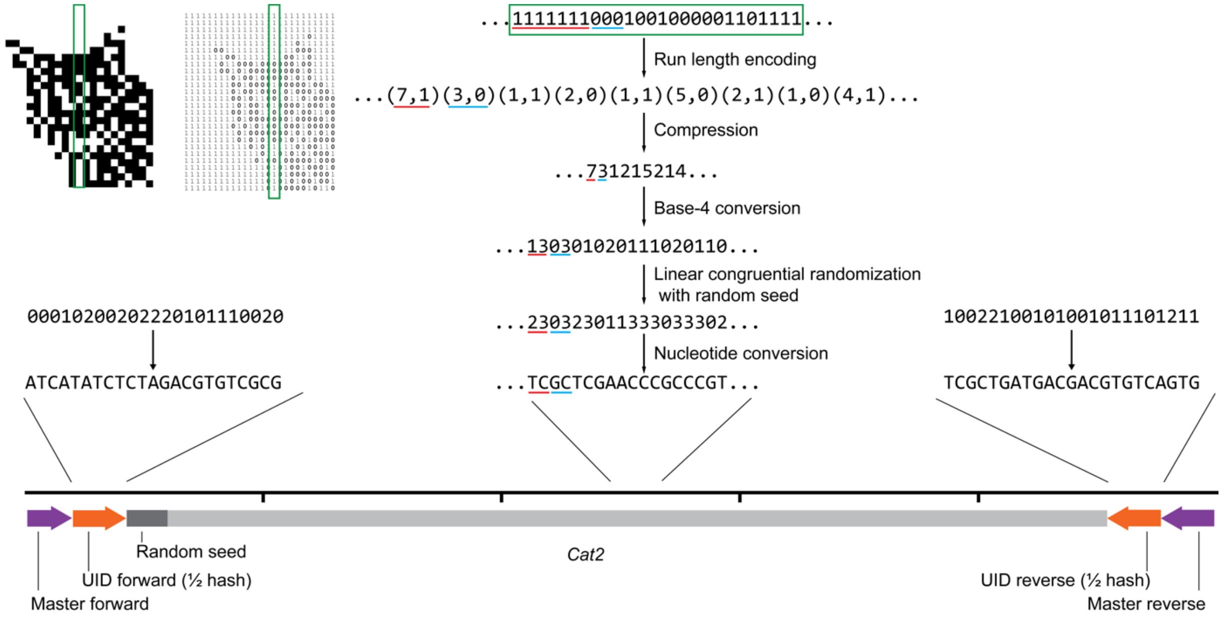

**Supplementary Figure 2. DNA encoding of representative icons.** The black and white 26 × 26-pixel icon is converted to 1 (white) or 0 (black) and the bitmap is generated, followed by run length encoding compression, a base-4 conversion, encryption by simple randomization with a seed (encoded in the DNA), and converted to nucleotides. A binary 64-bit wavelet hash is calculated from the icon image and split in half and converted to DNA to act as a UID primer pair (orange). A master primer pair is added to flank the construct. *Cat2* is shown as an example, but all icon images go through the same workflow.

**Supplementary Table 1. DNA sequences for encoded plasmids.** Purple text indicates master primer regions and orange text indicates hash primer regions.

| Image | Insert sequence (5' to 3' direction) |
|---|---|
| *Airplane* | CGTCGTCGTCCCCTCAAACTATCATGCTACACTGTACAGTGCGCATACTGAGACTGTATCGCACGAAATTTGAGATATTTCGTATC<br>CTTGCCTTTGCTAGGCTGAATTGTTTGGGCAGGCTCCTCGCACCATGATATTCGGGATAGCCTCCAGAATCTTAAGATTAAAATCC<br>AAAGGCTCCAGCCCTGACATCTTCCCAGAGATGCTTCCCTTCCGAGACTCGCTGGTGGTTATAGGCTACCTTTGTTATAGAACGAT<br>CGCTGCTCTGACTGTAGATATCGCTGAAAAGGTGGCATCAAT |
| *Apple* | CGTCGTCGTCCCCTCAAACTATCATATCTCGTCGATCGCTCAGAATGAGCTGTGACGCGTGATCTAACATGATTTTTGGAACGGGA<br>TCGGGAACGGTGGTCAACCTGCTTTTCTCTTCGGCTTAGCATGCAGTCTCATTCCGACGATCATCGCGCTCCCATAGGATATACCA<br>AATTCATATGAACAGTTCCTGTCTATAGGTCCATAATTCAGTTCTTACATGCTGAGAATTGAATGAGGCTTTCACTTTTCTGCGGA<br>TATCGCTGACACTATAGACGTGCTCGCTGAAAAGGTGGCATCAAT |
| *Banana* | CGTCGTCGTCCCCTCAAACTATCATCTGCGCTGACATGTAGCTCTATAGACTACACATGCACACTCCGTGTGTCATTGGAGGTGCT<br>CAGTCCGGTGCAACTTTTCCGAAGTACTTTTTGGTGGGCCAGATTTGACGGAAGCTTGGTGTACACTTAACCTGTGTTTATTGTCT<br>ACCAGGAATTAAACGACTTACGTTGGGACACTAGTCAGTAACATAGCCTCAACAAGTATCAGGCCAAACACATCATGTGAGATCGT<br>ATCTATCTCGCTGAAAAGGTGGCATCAAT |
| *Sailboat* | CGTCGTCGTCCCCTCAAACTATCATATCTCTATCTACTACTGTGTATATACTAGAGCTGCGACTCGCCCTACCGATTAGGAGTGCC<br>ACTGGGAGCAACCGCTAACTGTAAGGTGGGTCACCGCAGGTGCCGACCAAGAGCGAGGGCGGATGGAATATGAGGCATGCCGACTT<br>TGGCTACTGCACAACCGTAGTTTAGCCGCCGCCCGAGGACTCTAGAGTATAGAACCAGTACGACAGAAGCTAGTACTAAGTTTGCC<br>GATATGGCGTAGTGAGCGGTGCAAGGAGACGTCCTGGCTAATCATGCTGAGACGATGTCAGCTCGCTGAAAAGGTGGCATCAAT |
| *Skyscraper* | CGTCGTCGTCCCCTCAAACTATCATATCTCAGCTCATCTACTAGACATATGCATCGTGTCAGATGCGTTGTCCCAGGACTATGAGT<br>AGGTGTTATCACTACGCACACCAACTTGTGGTAGCGGGTAAGACATAGACGCGGATCCTCCTCGGCCTACGTAAAAATTCTCGCTG<br>CGTGACTTAAGACTCCTCGCAAGACGGAGATCTCGCCGTTGCCGGAAGAAGCCGCCGACCGTGGTGGAAAGGTCATCGTTACTAAC<br>GAGGCAGGCGCAAGATGCGTGTCCGTAGGCCGAGTGGATAATGACCGTGCGAGTGATCTACAGTGTCATAGCGGTACTCTGTACCG<br>GCGGTTCCCGATTGGTAAGTTACCACCATTTTGTACACTATCGTCAGCGTAGCATGCGCTATAGCTGAAAAGGTGGCATCAAT |
| *Canoe* | CGTCGTCGTCCCCTCAAACTATCATCACACTAGCGCTGTCGCACACTAGCAGTCGTACGTGTATTCCACTCATATCAACTCATGTT<br>CTGGTGTCGGTCGTAGCTGAGCTACACATGAATACCATTAGACTTTCCGGGCACGAATAAAACGCCCTGAGCACGAGAACCTAAGT<br>TCGTCAAACCCACTTCCAAGTCTGGGAACCAAAAATTATCCAAAACCGCTTCCTCCTTTCGACTGCTAGGATCCAAGCAATGGAAC<br>GCAAATGCCCAATAAGATGTGTACATGTATGCTTGCCATACGCGGTCAGCATCGCTGACAGATCGTCGCTGCTAGCTGAAAAGGTG<br>GCATCAAT |

| Image | Insert sequence (5’ to 3’ direction) |
|---|---|
| *Cat1* | CGTCGTCGTCCCCTCAAACTATCATGTCTATGAGTACACTCGCGATGTGACAGTCACAGATCGTCGCATAAGTGCTCCCACCGTGT<br>CATTGGCGCCAGTAATGTGCCATGTCATTCACTCTAAGAATAAGTGAAAATATGAAGCTGGGAGCGCCCGGAGATTAGTCTGCCAC<br>GACTCAACCTCTAGATCGAACGTCCTCTCTCCGATAGTTAGGCCATACTGGTACGTCATCAAGGCTTCGGGTTTGAATGAGCACCA<br>CAGTCCTCGCCTTTAACTGCGTCCGGCCTCTTGCAGGATTATCTTTAGTTATACTGGAAGTAGGTACTGAAAAGCATCTGCGCCCT<br>AGCAAGGCTTATTTTTAGGATCTCAGTGGGATGGAAAAGCGTATGCCACATGGTAGCGAAAAAGTCGTCTTGCTGTGCCGGAGTCG<br>GCTACGGCCTGATGAGCTGAGGCGGAGGGCTTGCCTTCGAAAGCTCCTACTAACAATTCAATAAATGTGGGAATGCTACATAAATT<br>GTCGTAGTACCGTCAGAGATAGGAGACGGGTCGATTAACACTTCTACGCAGGGTTATAATAGTTTGTGACACTCAGCAATCGCGAC<br>ACTACACGCTACGCACGCTGAAAAGGTGGCATCAAT |
| *Cat2* | CGTCGTCGTCCCCTCAAACTATCATATCTCTAGACGTGTCGCGAAGTCGACACAGCGTACACGCTGTCGGTTACAACGCAGTGAGC<br>TAACATATGCAGATCACGTTTGACGGGAGTACATTATAAACTCACCGCGAGCTTAGTCTCGAGCTCGCCGTGGAGATAGATAGTGT<br>GCCAGCATCGACGCGTCCGTCTTCGTCAGCAGTAACGCGCTCGAAACCAGTCTGAGACACGAAATCGTATATCTGTCATATTCGTC<br>CGATGGGCTATTGGACCGGACAGCTCTCGCTCGAACCCGCCCGTGCTAAAAATTTGCGGACTATTTTAAACAAGAGTCATCCTGTT<br>CTACTACAGTGAAGCCCTGTTGATGGGCGCGTGCTGGAGAGTAGTACGGATGACAACATTAACGACGAGGCCGGTACGAGATATAC<br>TAGGAATCGAGGTCGTCCCGAGCATCGCTGATGACGACGTGTCAGTGGCTGAAAAGGTGGCATCAAT |
| *Cat3* | CGTCGTCGTCCCCTCAAACTATCGCGCTCACACGCGAGACACGAATGCGAGCTATACGACATCTAACACCCGTACCGTCACTCCAG<br>GCGCGATGTAACGTTGTCTGGAGAGTTGCGTGTCTCCTCGCATCGAAGCAGCAGCATAGCTTGTTCTGGTTATCGCGCGTAAACAG<br>AGGCCACGGGCGTGAACCCGCCTCGCCGGCTTGCCCACGAAATCTGTGATTACAAATTCGTCCTCCTTCGTAAGGTGCTATCGCTG<br>AGCAGACGTGCACACTAGCTGAAAAGGTGGCATCAAT |
| *Cherries* | CGTCGTCGTCCCCTCAAACTATCATGCGCTAGCTCACAGAGATCTATATGTGCGTGAGCGAGTCGAGTTGCGCGGGTCACGTGCTA<br>ACCTAGAGTAATATATAGCGAAATTCACAAGTTCCGTACATTAGTTAAATGCATAATCCGATCCGCTACGCCGACTCGGCGAGGAG<br>GCAGAGCTGCTTGATGTTTTCAGGTCGATGCTCAAGGCCAGTACAAGATTGCTTGAGAGTTCCACCGAGCGGGCCGGGTTCGTGGT<br>CCGTCACTTGGCGTAGATCGCACGAGACTATGTAGAGTAGGCTGAAAAGGTGGCATCAAT |
| *Dog1* | CGTCGTCGTCCCCTCAAACTATCATATCACGCTACATCGAGCATACTCGACGAGTGTGCTCGCAAACGGGACGTGCAAGGTCCAAC<br>GTATAAGGTTCCCTATACCACAGAACACGGACGGGTATGCTACACCGGGATTTCTAACGCCCGTTCGGCCGCACATCTGGGACTTA<br>ATTGTGCTAGCAACTTAGCTGTTCATGTGCCTAGTGACGATATGTCACGTTGTCCCGTTGGGATCGCTGACACTATAGACTGCTGC<br>GCTGAAAAGGTGGCATCAAT |
| *Dog2* | CGTCGTCGTCCCCTCAAACTATCGCGCTCTGACGCGTCATCGACAGAGCACGTCGTGTCGCGTCTTCCATAATTAAAACGCAGGAC<br>TCGCGGGAACACCCATCGGATCGCATCCGAAGTATGGATAGGACTAAAGGAAACCGCGTGTGCTTGCAGTGGAACCTCTCCACCTT<br>CGCCAATGTTAAATGACCTAACATTGACAGGAATAGCGCAGCTCTTGCACATGCCCTGTCGAACGCATCAAGCGTGGGCTATCGTA<br>GATATACAGCAGAGCTGCGCTGAAAAGGTGGCATCAAT |
| *Flower* | CGTCGTCGTCCCCTCAAACTATCATATCTCATCTCTATCAGACGACACGCTGAGACGACATATGTAGCGGCTGTGGTTTATCACAG<br>CCTATCATTCAACCTATTATGGGTCAGTCGTCATATGAGCGAACTGTTGATGCGCTCCCAGGCGCTCAAATAACAGCCCAGACAGT<br>GCCCGTTCTCAGATCAGCTCCAGTTTAAACTGTGAGCGGGAGATAATAGCAGGCCAGGTACAGAGGAGCTGTTGACTAATGACTAA<br>TCGATCGACTGAGAGCATGTCGAGCTGAAAAGGTGGCATCAAT |
| *House* | CGTCGTCGTCCCCTCAAACTATCATATCTCACTGCGACTATCGATGAGACATACTCAGCTCGTAACCGTCCGGTGTGCAGGAAGCT<br>CTTTGTATTCGGCGTGTCACGCGGTACAAGGCCGTCATTTATCGGTCATGACACTCGAAAGTGAAAATATATATGGCTGTACCTTA<br>AAATGGCGTAACGCGACCCTTACGTAGCTATGAGCGCAGCAGCCTCTTAGTGTTCCTTGAGCGCCCTCATTACATCTGCCGTACCT<br>ACTATCGTAGTAGACAGCGCACGTCGCGCTGAAAAGGTGGCATCAAT |
| *Lincoln* | CGTCGTCGTCCCCTCAAACTATCATATCAGAGCACACACGCTCGATGATCTGTCGTGCGTCACTCTGATACTCACACTGCGCACGA<br>GGTGGCACTCCGAGATGGTGAGCTCACTGGCTCCGGAACGTGCCCGCCTTACCAGTCCACTTTCTAACCAAAGGTCGCCTCATGCA<br>AGCTATCCTTGTGTCTCGCGGAGCGCGCGAGCGATTGTCAGTTTAGTCTCGTTAGCTCAACACTGGCTATGATTAGTATTCGCACC<br>TTGTGCCGTATAGTAGTTACCCACTGTTAACACCGTGGGCATTGTCGGACAATAGTAAAACATCGACTCTGATATATGATGCTCAG<br>CTGAAAAGGTGGCATCAAT |
| *Lion* | CGTCGTCGTCCCCTCAAACTATCGCTAGCAGCGCTCTGCTCACAATACTCTATGAGCGCTCTACGAAAGTCTCCGTCTCTGAGAGC<br>GACATAAACTGCTCGCTACACGTCTCGGATCGATTTTACATCACACCGCAGCGATGGTTGAGCTCGCAGCTATCAATCTTAGCGAG<br>GAAAGCGTCGACCCTATGCATCACAAGGGCTATTACCGGTGACTCACCTTCAACTGGGCATCGGGCGTTGCATCATAGGTTACAAT<br>CAGTAGATATGTTCATGCTTCGTAATGTCCCATCATCTGCACGATGCAGTCGCTAGCTGAAAAGGTGGCATCAAT |
| *Tiger* | CGTCGTCGTCCCCTCAAACTATCATCTGCGCTGACGATCGTATATGATGACTGTACATAGACATTCCTGTCAAGCGGGCGGACTGC<br>TGGGTTACGAGACAGGTATGGGTTGCACGTCGATATGACCATGCCTAGATAGTCGCTCGTCGGGTGTCGCCAATGTTGATTGCTTG<br>GCTCATTACTTCCGGAGGAATCCCGACTGTATTGTTAAAGGTGGGCGGCATAGCTGTTTAATGCTGCCTATAACCCGAAGGGTTCC<br>GTCCATTGGATCGCTGTCTATCACGAGCTCTACGCTGAAAAGGTGGCATCAAT |
| *Tree* | CGTCGTCGTCCCCTCAAACTATCATATCTCTGCTACTCGCGTCGACTAGTCGTGCAGTCGCTATTCCCAACAACCACATGCCATAG<br>TTTTTATGATGCATACGGAGAGAGTCATGTGAGTTGGGCCGTATTCCGAGACAATCGTTTCATACCTAAGGAAACTGGTCACTTAC<br>ATTCCCTCGGTCTAAGAGCCAACACTTGATACTACACTAGCAACAGGCGGGTAGACCAATATGTATTTTGCATCAAAGTACTTGCG<br>TAACCTGTTATCGATCGATCGACATCTCTGACAGCTGAAAAGGTGGCATCAAT |
| *Washington* | CGTCGTCGTCCCCTCAAACTATCATATCGCTCGCTACGTCAGATTATGAGCAGTCATCACGACAGATGTCCATTCCAATGGTACAC<br>ATAGTGTGTCCTAGATCGACAGTATTCGCGGCGAGTATTATTTAGCGGCTCTGTTTCAGCACCGGAACAATAAGGACGAATAGTAT<br>ACATATGCCTAATACGTTTTCACGTCGGCACAACTCATAGAGAATATCCGGGATGACGCGTGACAATCAGTGATCAATGTAAGCTT<br>TTAACCGATTAGGTAGCGTCGTGGTTAACCGAGACTGTGAGTTACTCCTATGTTCTGCAATAGCCGAGGTCACCTTTGTTCATCAA<br>CTGGGCGGCCAATCGTAAATCCTGCTTATAGCTAGGATCATCGACGCGAGAGCTATCTAGGCTGAAAAGGTGGCATCAAT |

| Image | Insert sequence (5' to 3' direction) |
|---|---|
| *Wolf* | CGTCGTCGTCCCCTCAAACTATCATCTCTAGCGAGCTACAGCACTCACACACGATGACGATCATGTACGAGCTCGGCACTGATATG GAGACTCTCTATCTCAGGCCTCTACTTCTACACTGAGAGATTATCTGGTCTTTCGCCGCAGATTAGCTAAGCTTAAAAATCGTTCA ACGGTAACTAACCCTGCTTTCTTGAGGAACTTTCCGGGCCGTCCCGGTACATCGTACCCGTTTACTACCAACTTACAACACATGGC GCAGATCTAAGCATCGAGTTTCAAAATGCCATTTCTATGATACCCGCTGTACTCCCACCAAAGATACGAAATGCCCACCTTCTAAA AGTTATCTCCACATATAGAGCGACTACGCAGAAGGCAGGAATAACTCCTATATTTAGGTTTTAGCTCGGTTTATCGCTGACGCGCT CAGTACAGCAGCTGAAAAGGTGGCATCAAT |

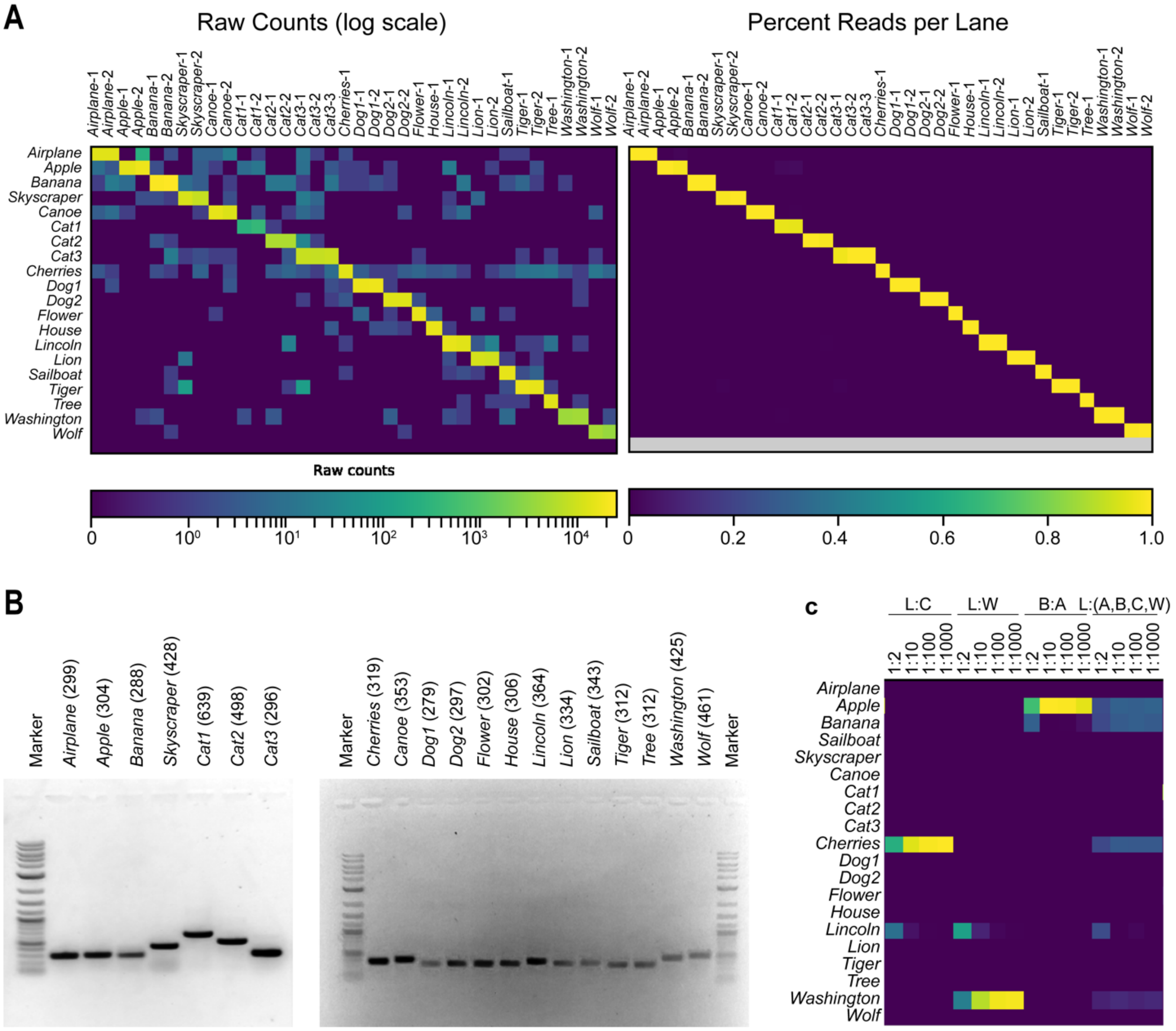

**Supplementary Figure 3. DNA plasmid database characterization.** (**A**) Illumina MiniSeq was used to validate the plasmids after DNA generation, showing baseline purity for each. (**B**) Each plasmid was amplified using the master primers and agarose gel was used to validate sizes, shown in parentheses in nt. (**C**) Dilutions of plasmid databases were generated, *Lincoln* (L) to *Cherries* (C), *Lincoln* to *Washington* (W), *Banana* (B) to *Apple* (A), and *Lincoln* to *Apple*, *Banana*, *Cherries*, and *Washington*. Illumina MiniSeq was used to read out the population after amplification with the master primers and addition of adaptors and sequencing barcodes.

### S3. DNA encapsulation

**Functionalizing 5-μm silica particle with TMAPS.** We adapted a procedure published previously[2]. A volume of 1.0 mL of 50 mg $mL^{-1}$ fluorescein-core 5-μm silica particles was added into a 2.0 mL DNA/RNA LoBind Eppendorf tube. The particles were centrifuged at 1,000 rpm for 10 seconds using a benchtop centrifuge. The particles were redispersed in 1.0 mL anhydrous ethanol with vigorous vortexing. The particles were centrifuged and redispersed in ethanol five times. We then added 50 μL of 50% TMAPS in methanol to the dispersed 5 μm silica particles (50 mg $mL^{-1}$ in ethanol). The mixture was stirred overnight at room temperature using a thermal mixer from Thermo-Fisher (Waltham, MA) at 1,200 rpm. The mixture was centrifuged at 1,000 rpm and washed with ethanol five times to remove any unreacted TMAPS. The functionalized particles were finally redispersed in 1.0 mL DNAse/RNAse-free water. The particles were stored at room-temperature until further use.

**Encapsulating plasmid DNA.** We adapted a procedure published previously[2]. For each data encoding plasmid, a mass of 1.0 mg of TMAPS-functionalized, fluorescent 5 μm particle was added into a 2 mL LoBind Eppendorf tube containing 15 μg of plasmid DNA dissolved in 1 mL of water. The mixture was mixed gently using a tube revolver (Thermo Fisher) at 30 rpm and at room-temperature for 5 minutes. A volume of 10 μL of 50% TMAPS in methanol was then added to mixture and stirred for 10 minutes, at 1000-rpm, and at 25 °C using a thermal mixer. After 10 minutes, a volume of 2 μL of TEOS was added and the mixture is stirred for 24 hours, at 1000 rpm, and at 25 °C using a thermal mixer (Thermo-Fisher). An additional 5 μL of TEOS was then added and the mixture is stirred for 4 days, at 1000 rpm, and at 25 °C which forms our encapsulated DNA particles or DNA capsules. The mixture is centrifuged at 2,000 × g for 3 minutes to sediment the DNA capsules then the supernatant was removed without disturbing the particle sediment pellet. The particles were washed repeatedly for 5 times by re-dispersing the particles with 1 mL of water, sedimenting the particles with a centrifuge at 2,000 ×g for 3 minutes, and removing the supernatant. After the final wash, the DNA capsules were re-dispersed in 1 mL of ethanol with 30 seconds of vortex mixing. A volume of 20 μL of γ-aminopropyltriethoxysilane was then added and the mixture was stirred for 18 hours, at 1000 rpm, and at 25 °C using a thermal mixer. The mixture is centrifuged at 2,000 × g for 3 minutes to sediment the amino-modified, DNA capsules then the supernatant was removed without disturbing the particle sediment pellet. The particles were washed repeatedly for 5 times by re-dispersing the particles with 1 mL of *N*-methyl-2-pyrrolidone, sedimenting the particles with a centrifuge at 2,000 × g for 3 minutes, and removing the supernatant. After the final wash, the DNA capsules were re-dispersed in 1 mL of *N*-methyl-2-pyrrolidone with 30 seconds of vortex mixing and the resulting colloidal suspension was then transferred into a clean 2 mL Eppendorf LoBind tube.

**Encapsulation efficiency.** After four days of encapsulation, all aqueous washes were collected. The amount of DNA that remained in each was estimated using qPCR (**Supplementary Fig. 4**). The washes were amplified using the master primer pair. All of the washes show plasmid concentrations that are below blank, suggesting that our encapsulation efficiency is quantitative.

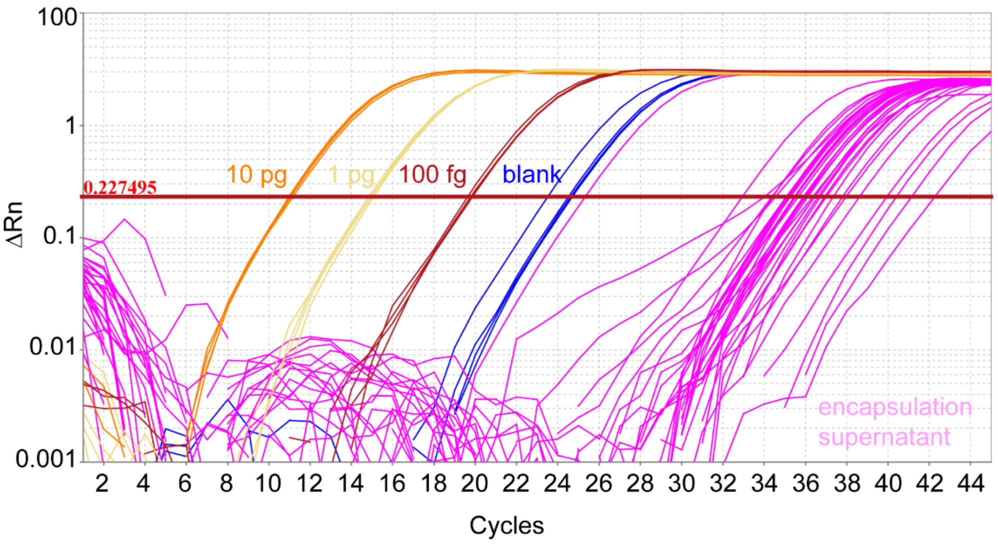

**Supplementary Figure 4.** PCR amplification curves (ΔRn vs Cycle, log scale) shown for the encapsulation supernatant for all plasmids after four days of encapsulation, with 10 pg (orange), 1 pg (yellow), and 100 fg (red) of the *Cat2* plasmid used for calibration. Overlay of 20 plasmids individually amplified with master primers (magenta) and blank (blue).

### S4. Barcoding DNA capsules

**Metadata descriptors DNA hash.** The subject matter of each of the original high-resolution images was associated with metadata. The subject matter included sets of cats and dogs, both wild and domestic, and of a variety of colors: a domestic black-and-white cat (*Cat1*), a domestic orange cat (*Cat2*), domestic brown cat (*Cat3*), a domestic brown dog (*Dog1*), a domestic black-and-white dog (*Dog2*), a wild yellow cat (*Lion*), a wild orange cat (*Tiger*), and a wild black-and-white dog (*Wolf*). Also included in the database were historical US presidents (*Washington*, an 18$^{th}$ century president, and *Lincoln*, a 19$^{th}$ century president); man-made objects (*Canoe*, *Skyscraper*, *Airplane*, *Sailboat*, and *House*); fruits (*Apple*, *Banana*, and *Cherries*); and plants (*Tree* and *Flower*) (**Supplementary Fig. 1**). From these descriptions, each encoded image was annotated with three semantic metadata descriptors (**Supplementary Table 2**) associated with the original image. A table was then generated to associate each descriptor with a unique sequence chosen from a list of 240,000 orthogonal barcode sequences [3].

**Supplementary Table 2. Metadata single-stranded key-value pairs.** Single-stranded DNA sequences were purchased from IDT. /5AmMC6/ denotes an amino hexyl modifier on the 5' end of each ssDNA sequence.

| **Images** | **Barcode 1** | **Sequence** | **Barcode 2** | **Sequence** | **Barcode 3** | **Sequence** |
|---|---|---|---|---|---|---|
| *Airplane* | *man-made* | /5AmMC6/ATGGATGCACGT<br>CCACAAGAAGCAG | *air* | /5AmMC6/TAGAAGCGTTCC<br>GACGAAGTTACCT | *flying* | /5AmMC6/GAGATTATTTCC<br>TCGTTCGGCCCAG |
| *Apple* | *fruit* | /5AmMC6/GAACTTATTCGG<br>TATCTGTGCCGCT | *red* | /5AmMC6/AGCGCTTGGGGA<br>CACGTGAAGTAAC | *seeds* | /5AmMC6/ACGTCACGTCGC<br>CTATGGCGTTATT |
| *Banana* | *fruit* | /5AmMC6/GAACTTATTCGG<br>TATCTGTGCCGCT | *seeds* | /5AmMC6/ACGTCACGTCGC<br>CTATGGCGTTATT | *yellow* | /5AmMC6/TAATGTGGCTTG<br>GCTCACCGCTAGG |
| *Canoe* | *man-made* | /5AmMC6/ATGGATGCACGT<br>CCACAAGAAGCAG | *water* | /5AmMC6/CTGGTTTGATCC<br>GACACATTGATTC | *oars* | /5AmMC6/GTTTCCGCATAA<br>ACTCAGGGGAGTC |
| *Cat1* | *cat* | /5AmMC6/AACGATTGTTAT<br>GCCCCTAACTCAG | *domestic* | /5AmMC6/TCTTAACAAAGG<br>ATGGGCAGGTCGC | *black & white* | /5AmMC6/TTCAGGGTGGAA<br>GTACCTCCCAGAT |
| *Cat2* | *cat* | /5AmMC6/AACGATTGTTAT<br>GCCCCTAACTCAG | *domestic* | /5AmMC6/TCTTAACAAAGG<br>ATGGGCAGGTCGC | *orange* | /5AmMC6/CTGAATACTACA<br>CGCCGTGGTGAAG |
| *Cat3* | *cat* | /5AmMC6/AACGATTGTTAT<br>GCCCCTAACTCAG | *domestic* | /5AmMC6/TCTTAACAAAGG<br>ATGGGCAGGTCGC | *brown* | /5AmMC6/ATCTATCTGTTG<br>GAGTTAACGTACC |
| *Cherry* | *fruit* | /5AmMC6/GAACTTATTCGG<br>TATCTGTGCCGCT | *red* | /5AmMC6/AGCGCTTGGGGA<br>CACGTGAAGTAAC | *pit* | /5AmMC6/GAGCCGATTTAG<br>TAGCAGTGTCCAA |
| *Dog1* | *dog* | /5AmMC6/AAAAGCAAGGTC<br>GTTACATGGAGTT | *domestic* | /5AmMC6/TCTTAACAAAGG<br>ATGGGCAGGTCGC | *brown* | /5AmMC6/ATCTATCTGTTG<br>GAGTTAACGTACC |
| *Dog2* | *dog* | /5AmMC6/AAAAGCAAGGTC<br>GTTACATGGAGTT | *domestic* | /5AmMC6/TCTTAACAAAGG<br>ATGGGCAGGTCGC | *black & white* | /5AmMC6/TTCAGGGTGGAA<br>GTACCTCCCAGAT |
| *Flower* | *plant* | /5AmMC6/TAAGCAATGGGT<br>TCCACACTACGTA | *white* | /5AmMC6/TTTTATGCCGTG<br>TTGTTGCGCGTAC | *yellow* | /5AmMC6/TAATGTGGCTTG<br>GCTCACCGCTAGG |
| *House* | *man-made* | /5AmMC6/ATGGATGCACGT<br>CCACAAGAAGCAG | *building* | /5AmMC6/GTAGTTCGGGGT<br>GCATACTACCTGA | *wood* | /5AmMC6/GGGCGCAGAAGT<br>CTCTATTCTAGAA |
| *Lincoln* | *human* | /5AmMC6/CATCGTAGGAAT<br>GCGGCCGAGAATC | *19th century* | /5AmMC6/CGATGTAGTCAT<br>CCCGATGTGCTGG | *president* | /5AmMC6/ATGGACGACTTG<br>GGACGGGTATCAA |
| *Lion* | *cat* | /5AmMC6/AACGATTGTTAT<br>GCCCCTAACTCAG | *wild* | /5AmMC6/ACTCCGAGGAAC<br>TTCGTGCTTAGTG | *yellow* | /5AmMC6/TAATGTGGCTTG<br>GCTCACCGCTAGG |
| *Sailboat* | *man-made* | /5AmMC6/ATGGATGCACGT<br>CCACAAGAAGCAG | *water* | /5AmMC6/CTGGTTTGATCC<br>GACACATTGATTC | *sails* | /5AmMC6/CTTACTTTCCTA<br>CTCACTTCTCCAC |
| *Skyscraper* | *man-made* | /5AmMC6/ATGGATGCACGT<br>CCACAAGAAGCAG | *building* | /5AmMC6/GTAGTTCGGGGT<br>GCATACTACCTGA | *steel* | /5AmMC6/TAGTGTGTGCCC<br>ACTGTAGCCGTGA |
| *Tiger* | *cat* | /5AmMC6/AACGATTGTTAT<br>GCCCCTAACTCAG | *wild* | /5AmMC6/ACTCCGAGGAAC<br>TTCGTGCTTAGTG | *orange* | /5AmMC6/CTGAATACTACA<br>CGCCGTGGTGAAG |
| *Tree* | *plant* | /5AmMC6/TAAGCAATGGGT<br>TCCACACTACGTA | *tree* | /5AmMC6/GATCAGAATCTA<br>CTCGCATAGCCTC | *moon* | /5AmMC6/AGTTAAATGTCC<br>CAGGCTTGTCACC |
| *Washington* | *human* | /5AmMC6/CATCGTAGGAAT<br>GCGGCCGAGAATC | *18th century* | /5AmMC6/GCAGTAAAGCTC<br>GGTCCGATCTTCA | *president* | /5AmMC6/ATGGACGACTTG<br>GGACGGGTATCAA |
| *Wolf* | *dog* | /5AmMC6/GAGTATCCGTTT<br>GATTTGTCGTCGC | *wild* | /5AmMC6/AGTCGTCCGAAA<br>TATTGCATTCTTG | *black & white* | /5AmMC6/TTCAGGGTGGAA<br>GTACCTCCCAGAT |

**Chemical attachment of DNA barcodes on DNA capsules.** Using all the DNA capsules from the previous step, a mass of 5 mg of β-azido acetic acid *N*-hydroxysuccinimide ester and 5 µL *N*,*N*-diisopropylethylamine as catalyst were added and the mixture was stirred for 2 hours, at 1000 rpm, and at 25 °C using a thermal mixer. The azide-modified DNA capsules were washed repeatedly for 5 times by re-dispersing the particles with 1 mL of *N*-methyl-2-pyrrolidone, sedimenting the azide-modified DNA capsules with a centrifuge at 2,000 × g for 3 minutes, and removing the supernatant. After the final wash, the azide-modified DNA capsules were re-dispersed in 1 mL of *N*-methyl-2-pyrrolidone. A mass of 2-mg of DBCO-PEG13-NHS ester was added and the mixture was stirred for 30 minutes, at 1,000 rpm, and at 25 °C using a thermal mixer. The particles were washed repeatedly for 5 times by re-dispersing the PEG-modified DNA capsules with 1 mL of *N*-methyl-2-pyrrolidone, sedimenting the PEG-modified DNA capsules with a centrifuge at 2,000 × g for 3 minutes, and removing the supernatant. After the final wash, the PEG-modified DNA capsules were re-dispersed in 200 µL of *N*-methyl-2-pyrrolidone with 30-seconds of vortex mixing and 1 minute sonication (Cole Parmer; Vernon Hills, IL). A volume 10 µL of each ssDNA barcode (500 µM in nuclease-free water) and 200 µL of PEG-modified DNA capsules were added to 770 µL of 0.1 M bicarbonate buffer (pH 9.2) in a 1.5 mL Eppendorf LoBind tube. The mixture was stirred for 2 hours, at 1000 rpm, and at 25 °C using a thermal mixer to produce the final form of our data blocks or "files". The files were washed repeatedly for 5 times by re-dispersing the particles with 1 mL of saline Tris-acetate-EDTA buffer with surfactants (40 mM Tris, 20 mM acetate, 2 mM EDTA, 1.0% Tween-20, 1.0% sodium dodecyl sulfate, and 500 mM NaCl), sedimenting the particles with a centrifuge at 2,000 × g for 3 minutes, and removing the supernatant. After the final wash, the particles were re-dispersed in 500 µL of saline Tris-acetate-EDTA (40 mM Tris, 20 mM acetate, 2 mM EDTA, 1.0% Tween-20, 1.0% sodium dodecyl sulfate, and 500 mM NaCl) with 30 seconds of vortex mixing and 1 minute sonication. All the files were then pooled together which forms the file pool or molecular file database with an estimated final concentration of 2.0 mg $mL^{-1}$ in 10.0 mL of 40 mM Tris, 20 mM acetate, 2 mM EDTA, 1.0% Tween-20, 1.0% sodium dodecyl sulfate, and 500 mM NaCl.

### S5. Estimating plasmid copy numbers in DNA files

We took different volumes, 10 µL, 50 µL, 100 µL, 150 µL, and 200 µL, from three different files that have particle concentrations at approximately 1 mg $mL^{-1}$. The particles were centrifuged for 10,000 × g for 1 minute and the supernatant was carefully removed using a pipette. The residue particles were dissolved using 45 µL of 5:1 buffered oxide etch and incubating the mixture for 5 minutes. The mixture was vortexed for 5 seconds to re-suspend the pellet and the mixture was statically incubated at room temperature for 5 minutes. A volume of 5 µL of 1 M phosphate buffer (0.75 M $Na_2HPO_4$; 0.25 M $NaH_2PO_4$; pH 7.5 at 0.1 M) was then added, vortexed for 1 second, and desalted twice through an Illustra MicroSpin S-200 HR column (GE Healthcare) before analyzing the samples through qPCR.

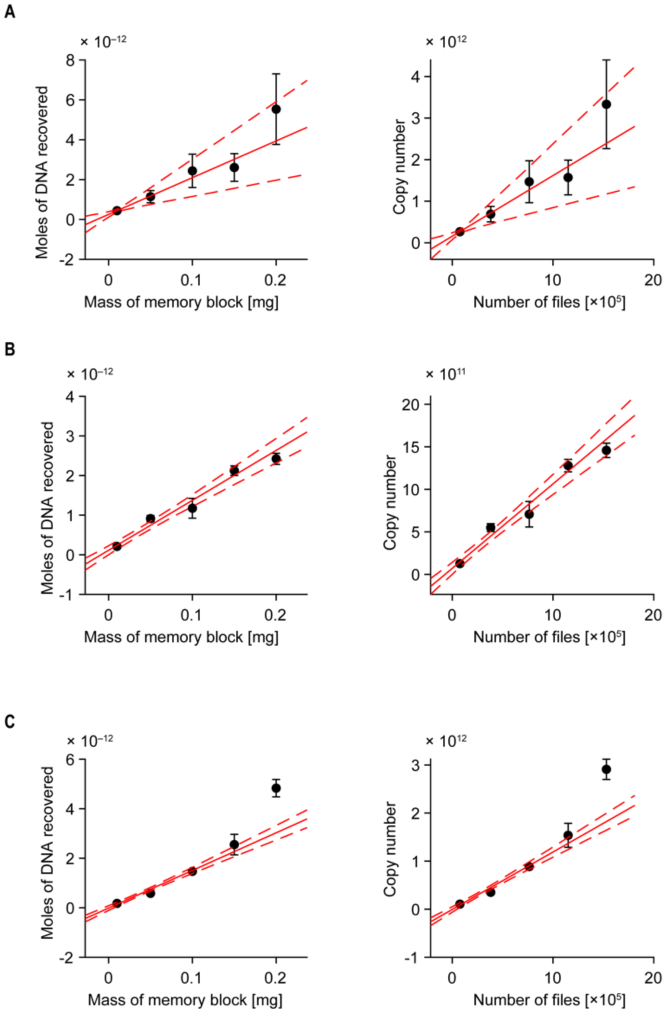

**Supplementary Fig. 5. Concentration and copy number of plasmid DNA for different memory files.** (**A**) *Cat2*, (**B**) *Sailboat*, (**C**) *Washington*. Solid red line denotes the best linear fit determined using Deming regression. Broken red lines are fit uncertainties.

**Supplementary Table 3. Fit results for Supplementary Fig. 5.**

| File | Slope [moles of DNA recovered per mass of files] | Slope [copy number per file] |
|---|---|---|
| *Cat2* | $1.84 \times 10^{-11} \pm 1.04 \times 10^{-11}$ | $1.46 \times 10^{6} \pm 0.83 \times 10^{6}$ |
| *Sailboat* | $1.26 \times 10^{-11} \pm 0.18 \times 10^{-11}$ | $0.99 \times 10^{6} \pm 0.14 \times 10^{6}$ |
| *Washington* | $1.52 \times 10^{-11} \pm 0.17 \times 10^{-11}$ | $1.20 \times 10^{6} \pm 0.13 \times 10^{6}$ |

### S6. Data density estimation

A hallmark of DNA as a data storage medium is the higher storage density compared to conventional memory systems such as flash and Blu-ray optical disc data storage[4]. For example, assuming 2 bits per base, $10^{-21}$ grams per base, and a density of double-stranded DNA of 1.7 grams per cubic centimeter, DNA has a theoretical volumetric density limit of $10^{27}$ bytes per $m^3$, a ~$10^9$-fold increase in data density over hard disk storage and ~$10^3$-fold increase in data density over flash storage. One means of DNA data access and retrieval is the use of digital microfluidics based on **e**lectro-**w**etting-**o**n-**d**ielectric (EWOD). Assuming the following properties of that system[5], namely an electrode area of $(2.7 \times 10^{-3}\ \text{m})^2$, a device thickness of $0.5 \times 10^{-3}$ m, and total bytes of data per electrode of $1 \times 10^{12}$ bytes, the total DNA data density dried on an electrode is

$$\frac{1\times10^{12}\,\text{bytes}}{\left(2.7\times10^{-3}\text{m}\right)^2\times0.5\times10^{-3}\text{m}} = 2.7\times10^{20}\ \text{bytes}/\text{m}^3$$

However, the use of an EWOD device leads to a reduction of data storage on DNA by a factor of ~$10^7$. In contrast, encapsulating DNA within silica capsules, as performed in the present work, assuming a plasmid copy number per silica particle of $10^6$ (**Supplementary Table 3**), the number of base pairs (bp) per plasmid ~3000, bits per bp = 2, and radius of the silica particle (**Figure 2**) = $3 \times 10^{-6}$ m, yields a total data density per silica particle of

$$\frac{10^6\ \text{plasmids}/\text{particle}\times3000\ \text{bp}/\text{plasmid}\times2\ \text{bits}/\text{bp}}{\frac{4}{3}\times\pi\times\left(3\times10^{-6}\ \text{m}\right)^3} = 5.3\times10^{25}\ \text{bits}/\text{m}^3 \approx 6.6\times10^{24}\ \text{bytes}/\text{m}^3$$

Our system is approximately four orders of magnitude greater in data storage density than the current EWOD implementation for data access and retrieval, but three orders of magnitude lower in density compared to when simply using bare DNA. Compared to conventional data storage systems, encapsulation of DNA within silica capsules has ~$10^6$-fold increase in data storage density over hard disk storage and ~$10^3$-fold increase in data storage density over flash storage. Thus, while encapsulation and barcoding leads to a ~$10^3$-fold reduction in density over bare DNA, our file system still has higher data storage density compared to EWOD-based retrieval. Further our approach provides a transformative gain in ability to sort and select data subsets efficiently through metadata labeling and barcoding for scalable random access and retrieval.

### S7. Query probes

An external table of probe sequences was generated associating each of these content descriptors to a DNA sequence database of reverse complements from the sequences displayed on the files and truncated to 15 nucleotides to maintain approximately 50 °C annealing temperatures using the IDT OligoAnalyzer tool (https://www.idtdna.com/pages/tools/oligoanalyzer). A 50 °C annealing temperature was chosen as the target temperature to easily de-hybridize the probe strands. A full-length 25-mer sequence would require annealing down from 95 °C and maintaining a temperature of 72 °C as designed for orthogonality and would introduce non-specific interactions at room temperature annealing, which would complicate sorting. Orthogonality of the truncated probe strands was tested computationally using the Nucleic Acids Package (NUPACK) [6-8].

**Synthesis and purification of fluorophore-labelled single-stranded DNA query probes.** A volume of 200 µL of a 1 mM (200 nmol) solution of hexylamine-modified single-stranded DNA reconstituted in 100 mM sodium bicarbonate buffer (pH 9.2) and 20 µL of 50 mM stock solution of either TAMRA or AFDye 647 NHS ester in DMSO (10,000 nmol, 50 equivalents) were added sequentially in a 1.5-mL LoBind Eppendorf tube. The reaction mixture was mixed at 25 °C using a thermal mixer for 2 hours after which it was passed through an Illustra NAP-5 gel filtration column (GE Healthcare; Marlborough, MA) for desalting and removal of residual small molecules, such as NHS, unreacted dye NHS esters, or dye acids that were formed due to hydrolysis.

The desalted reaction mixture was further purified with ion-pairing, reverse-phase high performance liquid chromatography (IP-RP-HPLC) using a Waters (Milford, MA) Alliance HPLC e2695 system equipped with a Waters 2998 photodiode array detector, Waters Fraction Manager–Analytical, and an XBridge Oligonucleotide BEH C18 2.1 mm × 50 mm column with a particle size of 2.5 µm. The aqueous mobile phase for IP-RP-HPLC is composed of 0.1 M triethylammonium acetate in HPLC-grade water (Millipore Sigma) with pH 7.0 while the organic mobile phase is composed of 0.1 M triethylammonium acetate in 90:10 (w/w) acetonitrile and HPLC-grade water. A focused gradient was optimized and used for all purification methods (**Supplementary Table 4**). All purification runs were run at a flow rate of 1.0 mL $min^{-1}$.

**Supplementary Table 4. Focused gradient table for IP-RP-HPLC purification of DNA-dye conjugates.**

| Dye conjugate | Hold | Linear gradient | Hold | Clean-up linear gradient | Clean-up hold | Re-equilibration linear gradient | Re-equilibration hold |
|---|---|---|---|---|---|---|---|
| AFDye 647 | Aqueous: 87%, Organic: 13% Time: 0–1 minute | Aqueous: 87% → 85% Organic: 13% → 15% Time: 1–16 minutes | Aqueous: 85% Organic: 15% Time: 16–17 minutes | Aqueous: 85% → 50% Organic: 50% → 50% Time: 17–18 minutes | Aqueous: 50% Organic: 50% Time: 18–19 minutes | Aqueous: 50% → 87% Organic: 50% → 13% Time: 19–20 minutes | Aqueous: 87% Organic: 13% Time: 20–23 minutes |
| TAMRA | Aqueous: 83%, Organic: 17% Time: 0–1 minute | Aqueous: 83% → 81% Organic: 17% → 19% Time: 1–16 minutes | None | Aqueous: 81% → 50% Organic: 19% → 50% Time: 16–17 minutes | Aqueous: 50% Organic: 50% Time: 17–18 minutes | Aqueous: 50% → 83% Organic: 50% → 17% Time: 18–19 minutes | Aqueous: 83% Organic: 17% Time: 19–23 minutes |

The column was heated and maintained at 50 °C using a column temperature controller and thermostat. Absorption intensities at 260 nm and 550 nm for TAMRA-DNA conjugates and 260 nm and 647 nm for AFDye 647-DNA conjugates were measured and used to automatically determine when to collect fractions. Collected purified fractions were dried to pellet form using a SpeedVac SPD300 (Thermo Fisher). The pelleted dye-modified DNA strands were reconstituted in 50 µL of HPLC-grade water and characterized with matrix-assisted laser desorption/ionization time-of-flight mass spectrometry using a Bruker (Billerica, MA) microflex to confirm conjugation of the dye. Oligothymidine single-stranded oligonucleotide standards (Waters MassPREP OST) were used to calibrate the mass spectrometry measurements. The final concentrations of the dye-labelled single-stranded DNA probes were determined by measuring the absorbance spectra from 190–840 nm using a Nanodrop 2000 (Thermo Fisher) and using the molar absorption coefficients of the dyes at the maximum absorbance peak (AFDye 647: 270,000 $M^{-1}$ $cm^{-1}$; TAMRA: 92,000 $M^{-1}$ $cm^{-1}$) to calculate the concentrations.

**Supplementary Table 5. Sequences and mass characterization of dye-labelled query DNA probes.** MW = molecular weight.

| Query barcode | Sequence | Expected MW (Da) | Measured MW (Da) |
|---|---|---|---|
| *18th century*-AFdye 647 | AFDye647-C6-TGAAGATCGGACCGA | 5660.0 | 5660.2 |
| *cat*-AFDye 647 | AFDye647-C6-CTGAGTTAGGGGCAT | 5682.2 | 5688.1 |
| *flying*-AFDye 647 | AFDye647-C6-CTGGGCCGAACGAGG | 5677.2 | 5675.3 |
| *fruit*-AFDye 647 | AFDye647-C6-AGCGGCACAGATACC | 5605.2 | 5607.9 |
| *wild*-AFDye 647 | AFDye647-C6-CACTAAGCACGAAGT | 5604.2 | 5601.0 |
| *building*-TAMRA | TAMRA-C6-TCAGGTAGTATGCAC | 5183.6 | 5176.5 |
| *dog*-TAMRA | TAMRA-C6-AACTCCATGTAACGA | 5136.6 | 5134.7 |
| *president*-TAMRA | TAMRA-C6-TTGATACCCGTCCCA | 5079.5 | 5077.1 |
| *yellow*-TAMRA | TAMRA-C6-CCTAGCGGTGAGCCA | 5169.6 | 5167.1 |

**Computational and experimental barcode characterization.** The orthogonality of barcode and probe sequences was confirmed using NUPACK[6-8] to estimate the energetic favorability of binding between a single-stranded barcode and a single-stranded probe molecule in solution. These estimates do not account for the surface effects of having multiple barcodes densely packed on the surface of the silica bead. Using NUPACK's complexes function, the difference in energy between the double-stranded probe-barcode complex and the two single-stranded complexes at room temperature (25 °C) was calculated as the free energy of binding between the two DNA sequences. The binding energy for all probe-barcode pairs used during our experiments is shown in **Supplementary Fig. 6a**. Overall, these results show that the binding affinity is much stronger between correct probe-barcode pairs as compared to incorrect probe-barcode pairs, which should be orthogonal to one another.

We assayed the orthogonality of our truncated probes using flow cytometry (BD Aria III). Briefly, we added 1 µL of 500 µM of a fluorescently-labelled probe to each individual file (20 µL of 2 mg $mL^{-1}$ in 40 mM Tris, 20 mM acetate, 2 mM EDTA, 1.0% Tween-20, 1.0% sodium dodecyl sulfate, and 500 mM NaCl) that are physically separated into wells in a 96-well plate. The plates were heated at 80 °C for 5 minutes using a thermal cycler (Thermo Fisher QuantStudio 6) then mixed at 1000 rpm using a thermal mixer (Thermo Fisher). The plate was centrifuged at 2,000 ×g for 2 minutes and the supernatant was removed. A volume of 100-µL of 40 mM Tris, 20 mM acetate, 2 mM EDTA, 1.0% Tween-20, 1.0% sodium dodecyl sulfate, and 500 mM NaCl was added to wash the beads through five repeated cycles of centrifugation and removal of supernatant. The particles were finally dispersed with 200-µL of 40 mM Tris, 20 mM acetate, 2 mM EDTA, 1.0% Tween-20, 1.0% sodium dodecyl sulfate, and 500 mM NaCl then analyzed using flow cytometry. Gates were drawn in FlowJo (version 10) based on the fluorescence signal histogram of the negative control (silica beads only) (see **Supplementary Section S10** for additional details). Any particle with a fluorescence signal above the negative control threshold is considered positive for the specific probe being tested. This entire process was repeated for all the probes used for sorting in this manuscript (**Supplementary Fig. 6b**). In this flow cytometry assay, we observed very small non-specific binding of the probes to other barcodes that are not perfect complementary, consistent with computational predictions from NUPACK.

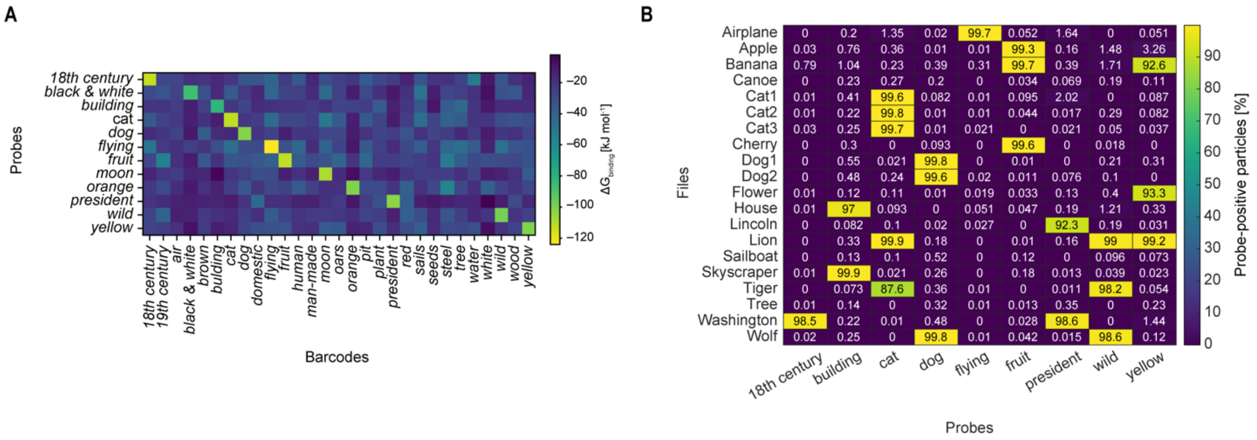

**Supplementary Figure 6. Characterization of probe orthogonality.** (**A**) NUPACK estimates of binding affinity at 25 °C between barcode-probe pairs. Affinities were estimated at 1M NaCl and without $MgCl_2$. Only probes actually used for sorting were tested. Binding affinity is strongest for correct pairs, although some interactions between non-orthogonal pairs exist. (**B**) Experimental validation of probes orthogonality against non-complementary barcodes on files. We used flow cytometry to analyze how many particles would contain the fluorescent probe.

### S8. Estimating surface-accessible DNA barcodes using DNA hybridization assay

DNA hybridization assay was used to estimate the number of surface-accessible DNA barcodes [9-11]. In a typical experiment, we add 1 µL of 500 µM of TYE705-modified single-stranded DNA probe to a 1.5 mL Eppendorf LoBind tube that contains 50 µL of 2 mg $mL^{-1}$ of files that has surface-attached barcodes that are complementary to the TYE705-modified DNA probe sequence. Negative controls, with sample volumes of 50 µL of 2 mg $mL^{-1}$ in 1.5-mL Eppendorf LoBind tubes, were measured simultaneously with the test files. These negative controls either have surface-attached barcodes that are orthogonal to the TYE705-modified DNA probe sequence or hydroxy-terminated silica surface. Upon addition of the TYE705-modified DNA probe sequence into the file solution, the mixtures were mixed at 70 °C at 1,200 rpm using a thermal mixer for 5 minutes. The mixtures were cooled to 20 °C at 1,200 rpm using thermal mixer over 20 minutes and then centrifuged at 10,000 × g for 1 minute. The supernatant was collected, and the pelleted particles were washed with 80 µL of 40 mM Tris, 20 mM acetate, 2 mM EDTA, 1.0% Tween-20, 1.0% sodium dodecyl sulfate, 500 mM NaCl. The sedimentation and washing process was repeated for five additional times while collecting the supernatant each cycle and pooling all the collected supernatant. A calibration curve using the TYE705-modified single-stranded DNA probe was used to determine the concentration of unhybridized TYE705-modified single-stranded DNA probe that remained in the supernatant solution (**Supplementary Fig. 7**).

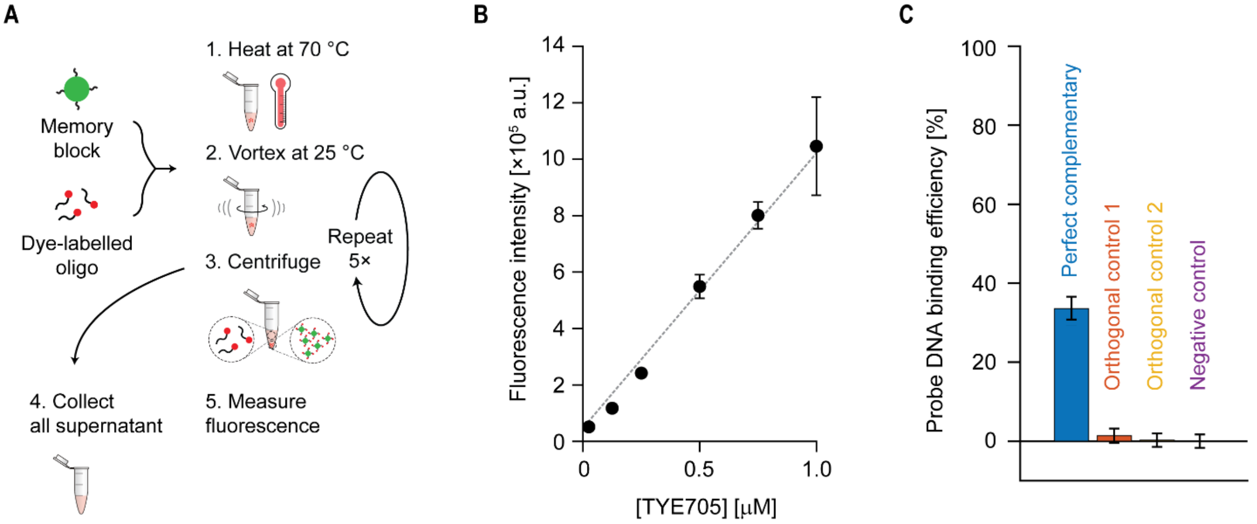

**Supplementary Figure 7. Hybridization assay to estimate the number of surface-accessible DNA barcodes.** (**A**) Sampling and analysis workflow. (**B**) Calibration curve used to determine the concentration of remaining unbound TYE705-modified DNA probes which encodes for the *black & white* barcode. Buffer for all dilutions: 40 mM Tris, 20 mM acetate, 2 mM EDTA, 1.0% Tween-20, 1.0% sodium dodecyl sulfate. (**C**) Probe DNA binding efficiency determined from the concentration of unbound TYE705-modified DNA probes that are left in the supernatant. Error bars are standard deviations from three independent replicates. Perfect complementary: File (*Dog2*) that has a *black & white* barcode. Orthogonal control 1: *Cat1*. Orthogonal control 2: *Airplane*. Negative control: silica core with hydroxy-terminated surface.

We used the density of silica and mass of silica that was sampled to estimate the number of silica particles in solution, and subtract the difference of initial TYE705-DNA concentration and the concentration of TYE705-DNA that remained in the supernatant to determine the concentration of hybridized TYE705-DNA. The number of TYE705-DNA that were hybridized onto the file surface can be calculated by taking the product of the volume of the supernatant, the concentration of the hybridized TYE705-DNA, and Avogadro's number ($6.022 \times 10^{23}$ objects $mole^{-1}$). Assuming that the hybridization efficiency is unity, the ratio of the number of hybridized TYE705-DNA and number of particles in solution provides the estimate of the number of surface-accessible barcodes per particle. The calculation is outlined as follows:

| | |
|---|---|
| **Number of silica particles** | Concentration of silica particles ($C_{\text{silica}}$) = 2.0 mg mL$^{-1}$<br>Volume sampled ($V_{\text{silica}}$) = 0.050 mL<br>Mass of silica particles in solution ($M_{\text{silica}}$ )= $C_{\text{silica}} \times V_{\text{silica}}$ = 1 × 10$^{-4}$ g<br><br>Density of a silica particle ($D_{\text{particle}}$) = 2.0 g mL$^{-1}$<br>Volume of a silica particle ($V_{\text{particle}}$) = 4 × π × (2.5 × 10$^{4}$ cm)$^{3}$ / 3 = 6.5 × 10$^{-11}$ cm$^{3}$<br>Mass of a silica particle ($M_{\text{particle}}$) = $D_{\text{particle}} \times V_{\text{particle}}$ = 1.3 × 10$^{-10}$ g<br><br>Number of silica particles ($N_{\text{particle}}$) = $M_{\text{silica}}$ / $M_{\text{particle}}$ = 7.6 × 10$^{5}$ particles |
| **Number of hybridized TYE705-DNA** | Initial concentration of TYE705-DNA ($C_{\text{initial}}$) = 500 μM<br>Volume of TYE705-DNA used ($V_{\text{used}}$) = 1 μL<br>Moles of TYE705 used ($m_{\text{initial}}$) = $C_{\text{initial}} \times V_{\text{used}}$ = 5 × 10$^{-10}$ moles<br><br>Measured concentration of TYE705-DNA from calibration curve ($C_{\text{supernatant}}$) = 0.740 ± 0.05 μM (mean ± s.d., n = 8)<br>Total volume of supernatant ($V_{\text{supernatant}}$) = 450 ± 11 μL (mean ± s.d., n = 8)<br>Moles of TYE705-DNA in supernatant ($m_{\text{supernatant}}$) = $C_{\text{supernatant}} \times V_{\text{supernatant}}$ = 3.3 ± 0.2 × 10$^{-10}$ moles (propagated uncertainty: mean $m_{\text{supernatant}}$ × √ (s.d. $V_{\text{supernatant}}$ / mean $V_{\text{supernatant}}$)$^{2}$ + (s.d. $m_{\text{supernatant}}$ / mean $m_{\text{supernatant}}$)$^{2}$)<br><br>Moles of hybridized DNA ($m_{\text{hybridized}}$) = $m_{\text{initial}} - m_{\text{supernatant}}$ = 1.7 ± 0.2 × 10$^{-10}$ moles (propagated uncertainty: √ (s.d. $m_{\text{supernatant}}$)$^{2}$)<br><br>Number of hybridized DNA ($n_{\text{hybridized}}$) = $m_{\text{hybridized}}$ × 6.022 × 10$^{23}$ objects mole$^{-1}$ = 1.0 ± 0.1 × 10$^{14}$ hybridized DNA (propagated uncertainty: mean $n_{\text{hybridized}}$ × √ (s.d. $m_{\text{hybridized}}$ / mean $m_{\text{hybridized}}$)$^{2}$) |
| **Number of surface-accessible DNA barcodes** | Number of surface-accessible DNA barcodes per particle ($n_{surface}$) = $n_{\text{hybridized}}$ / $N_{\text{particle}}$ = 1.3 ± 0.1 × 10$^{8}$ surface-accessible DNA barcodes per particle (propagated uncertainty: mean $n_{surface}$ × √ (s.d. $m_{\text{hybridized}}$ / mean $m_{\text{hybridized}}$)$^{2}$) |

### S9. Sequencing analysis

Three methods were used for verification of retrieval of DNA, which included quantitative PCR (qPCR), next-generation sequencing, and bacterial transformation and Sanger sequencing. For qPCR, standard curves were generated for each of the 20 plasmids using the 20 hash barcode primer pairs for 100 fg, 1 pg, and 10 pg of each, as judged by dilution from absorbance at 260 nm using the Nanodrop.

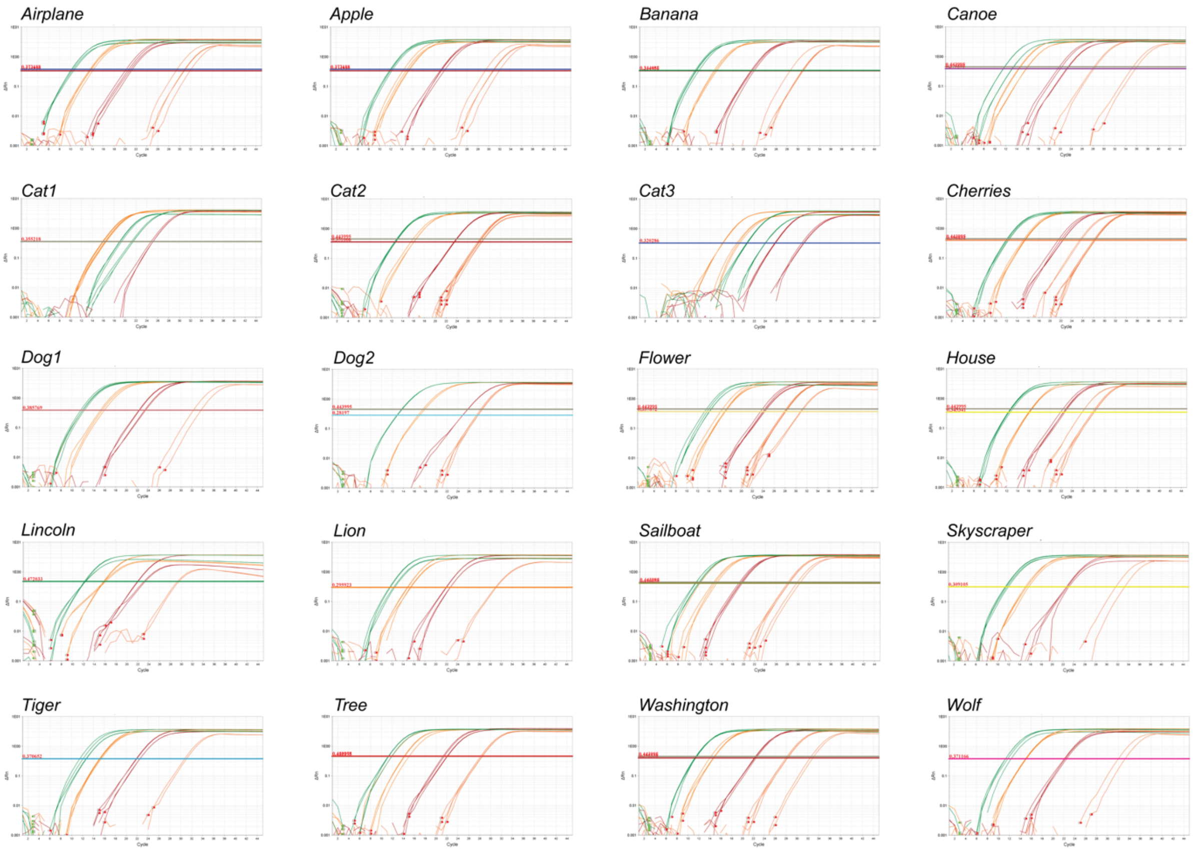

**Supplementary Figure 8. Plasmid standard curves.** PCR amplification curves (ΔRn vs. Cycle, log scale) shown for each plasmid DNA, for both master and hash UID primers, with 100 fg (red), 10 pg (orange), and 100 pg (green) of the indicated plasmid.

Gram weights were converted by molecular weight to moles to copy number, and the stand curve was generated as the log of the amounts vs. the threshold cycle. Fit curves were generated and the slopes are reported in **Supplementary Fig. 9**. Each file copy number approximately doubling per cycle. A simple plasmid mixture composed of equimolar amounts of *Cat2*, *Lincoln*, and *Washington*, and 5× enriched of each of the 3 plasmids, with qPCR using the hash barcodes for amplification, showing capabilities of qPCR in isolating enriched populations.

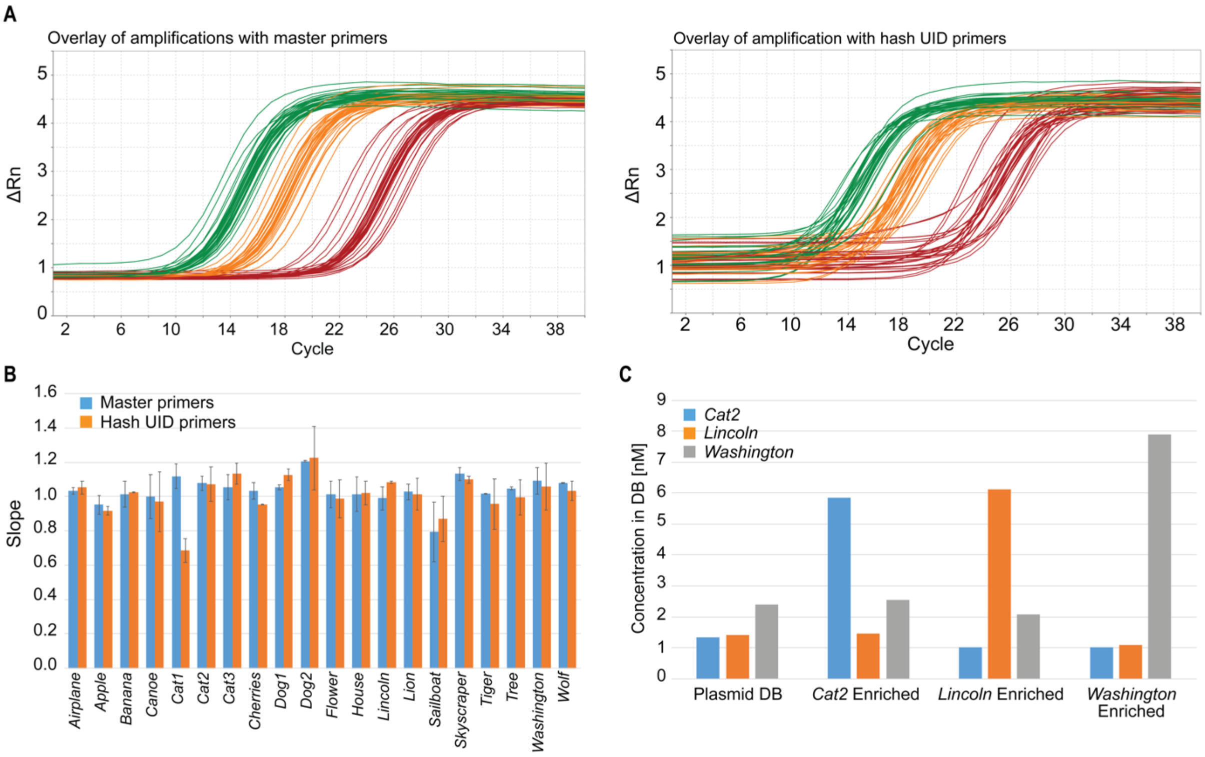

**Supplementary Figure 9. Quantitative PCR analysis of plasmid.** (**A**) Overlay of 20 plasmids individually amplified with master primers (left) and hash UID primers (right), with 100 fg (red), 10 pg (orange), and 100 pg (green). (**B**) Calculated slopes from each standard curve from each master (blue) and hash UID (orange) primer amplification series, error bars are standard deviation of triplicate measurements. (**C**) Plasmid database with approximately equimolar concentrations, or with *Cat2*, *Lincoln*, or *Washington* 5 × enriched, as shown, with qPCR with hash UID primers were used to quantify each of the three memory plasmids for each of the four databases.

For Illumina MiniSeq and MiSeq sequencing, the master primer pair with 5′ extensions matching Illumina Nextera sequencing adapters were used to amplify all plasmids simultaneously (**Supplementary Fig. 9**). Template amounts were adjusted based on concentrations determined with Qubit fluorescence assay (Thermo Fisher) or qPCR. If required, the amplification was simultaneously followed by qPCR and enough cycles were used to rise above the Ct, or alternatively obtain a final concentration of 2 ng $\mu L^{-1}$. Dual sequence indices were then added to the adaptor-modified inserts at the 5′ and 3′ ends, associating the sequencing lane with a particular logic-gated pull, which was followed by SPRIbead cleanup. A 25 µL PCR reaction amplified the material over 8–10 cycles using Kapa HiFi polymerase with 1 ng of template and 1 µM forward and reverse primers. After amplification, this was combined with 20 µL of SPRIselect beads, mixed, and let stand for 5 min. The mix was then separated by magnetic plates, and washed twice with 150 µL 80% ethanol, dried for 2 min, and eluted in 20 µL Qiagen TE buffer. Samples were quantified using the Qubit fluorescence assay with the provided high-sensitivity buffer and standards. A sequencing pool was generated to approximately equimolar amount per index pair. Illumina MiniSeq with 150 × 150 read lengths was used to read out the start and end of each sequence. Sequences were demultiplexed, and sequence clustering was used to count the number of occurrences of each image.

**Supplementary Table 6. Primer sequences for sequence adaptor addition**. Master primer sequence is underlined.

| | |
|---|---|
| Mem_R1 | TCGTCGGCAGCGTCAGATGTGTATAAGAGACAGC<u>GTCGTCGTCCCCTCAAACT</u> |
| Mem_R2 | GTCTCGTGGGCTCGGAGATGTGTATAAGAGACAGGCAG<u>ATTGATGCCACCTTTTCAGC</u> |

MiniSeq and MiSeq reads were clustered through a two-pass procedure that focused solely on the “hash” sequences flanking the image encoding internal regions of each sequence, because each hash sequence provided a unique identifier distinct between all sequences in our test set. UID’s were extracted from each read and clustering was subsequently performed with the following algorithm:

1) Create an empty list to store all observed clusters.
2) For each read do the following:
    a. Check if the Hamming distance between the read’s UID and any cluster UID is at or below a predetermined threshold of 5. This distance threshold was determined as significantly below the Hamming distance between any pair of correct UIDs corresponding to one of the expected sequences (see **Supplementary Fig. 10**).
    b. For any clusters satisfying this criterion, update the nucleotide counts observed at each position in the UID. Update the consensus UID of this cluster by a simple majority vote of the base identity at each position.
    c. Periodically perform the following cluster clean-up check: If a cluster UID has a Hamming distance to one or more other clusters at or below the distance threshold, merge these clusters and update their nucleotide counts at each position and the consensus sequence. For all analyses, this was performed every 20,000 reads.
3) After processing all reads, remove any clusters that make up less than 0.02% of the total reads observed.
4) For each read, assign it to all clusters for which its UID satisfies the Hamming distance threshold. Some reads may not be assigned to any clusters or to multiple clusters, although in practice the latter occurrence was quite rare. Do not update any clusters during this step.
5) Each cluster’s UID was compared against the correct hash for each of the expected sequences and assigned to the one with the lowest Hamming distance, if this distance was less than the threshold. These assignments were used as the counts for each file in that sample.

As a control, clustering on some samples was also performed using an internal region of each sequence rather than the hash sequence, with minimal change to the results (data not shown). Code used to perform clustering is available on Github (https://github.com/lcbb/DNA-Memory-Blocks/). The sort probability for a file into a particular fraction was calculated as the count associated with that file divided by the sum of the counts for that file over all fractions generated from an initial sample. This metric is also referred to as *enrichment* throughout the text. In **Figs. 3–5** in the main text, the enrichment of each file is indicated by the percent opacity of the images displayed on the grid.

**Image reconstruction with Illumina MiSeq sequencing.** Reconstruction of the original images was carried out using Illumina MiSeq ran with 300 × 300 read lengths, which span the plasmids sequences that encode images completely except for *Cat1*, which was 649 nucleotides in length. For this image, the clustering statistics were used although the image shown is not reconstructed from the sequencing results. To perform reconstruction, each pair of forward and reverse reads were aligned relative to each other to obtain a complete sequence for that read. From all sequences within each cluster, a consensus sequence was generated by determining the nucleotide identity that was most commonly observed at each nucleotide position. From this consensus sequence, the image was reconstructed by reversing the encoding process described in **Supplementary Section S2**. With the exception of *Cat1*, all images were reconstructed successfully. Error rates were low enough that image reconstruction was almost always possible with as few as three reads. Results of the image reconstruction are shown in **Supplementary Fig. 10**.

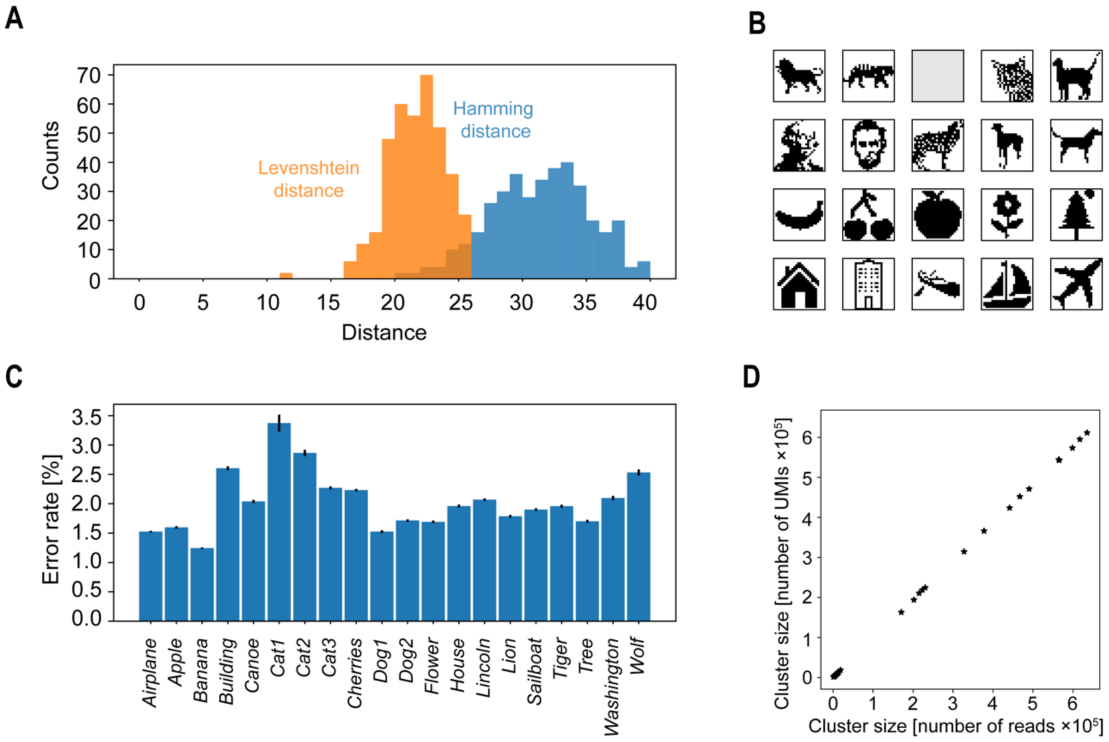

**Supplementary Figure 10.** (**A**) Hamming and Levenshtein distances between all pairs of hashes taken from file DNA sequences. Hamming distance was used rather than Levenshtein distance during clustering to reduce processing time. The minimum Hamming distance between any pair was 20, indicating that the distance threshold of 5 used during clustering is sufficient to avoid clustering correct sequences. **(B)** Image reconstructions from Illumina MiSeq 300 × 300 sequencing, performed on a pool containing all files. Images were successfully reconstructed for all templates with the exception of *Cat1*, whose length (649 nucleotides) prevented full sequencing. (**C**) Sequencing error rates per base for each template. Error rates ranged from 1% to 3.5%, which is consistent with previous literature on sequencing of DNA de-encapsulated from silica particles [12]. Error bars on the error rates show standard errors of the mean. (**D**) To determine if PCR amplification bias could affect the relative counts of each file sequence during sequencing, universal molecular identifiers (UMIs) were added to some samples prior to PCR amplification. The size of each cluster was recalculated as the number of unique UMIs in that cluster. UMIs were 12-nt long random sequences added to the 3' and 5' ends of the file sequences, and two UMIs with a Levenshtein distance less than or equal to 1 were considered equivalent. The data show that the number of UMIs in each cluster was scaled linearly with the number of reads in that cluster, indicating that UMIs were not necessary for accurately measuring relative cluster size.

### S10. Fluorescence sorting of files

**Querying molecular file database using fluorescently-labelled probes.** The molecular file database was vortexed for 10 seconds, sonicated for two minutes, and re-vortexed for another 10 seconds to re-disperse the settled particles. A volume of 100-µL of the molecular database (2 mg $mL^{-1}$) is added into a 1.5 mL Eppendorf LoBind tube. Dye-labelled probes for querying the molecular file database were added such that the final concentration of the DNA-dye single-stranded DNA in solution is 5 µM. The resulting mixtures were mixed at 70 °C at 1,200-rpm using a thermal mixer for 5 minutes. The mixtures were then cooled to 20 °C at 1,200 rpm using a thermal mixer over 20 minutes and then centrifuged at 10,000 × g for 1 minute. The supernatant was discarded, and the pelleted particles were washed with 500-µL of 40 mM Tris, 20 mM acetate, 2 mM EDTA, 1.0% Tween-20, 1.0% sodium dodecyl sulfate, 500 mM NaCl. The sedimentation and washing process was repeated for five additional times to remove non-specifically bound dye-DNA. The particles are finally re-suspended in 500 µL of 40 mM Tris, 20 mM acetate, 2 mM EDTA, 1.0% Tween-20, 1.0% sodium dodecyl sulfate, 500 mM NaCl.

**Fluorescence-activated sorting.** All fluorescence-activated sorting (FAS) experiments were performed on a BD FACSAria III flow cytometer. Samples were filtered through a Corning® 70-µm cell strainer (Fisher Scientific) prior to particle sorts. Samples are flowed into the instrument with 1× PBS as sheath fluid at a flow rate that maintains an events detection rate of 1,200 events per second and below. We found that performing sorting at a flow rate that exceeds this events rate clogged the FAS instrument intermittently. All sorts were accomplished with a standard 70 µm nozzle. The sample was held at room-temperature and agitated periodically every 5 minutes by stopping the sort and vortexing the sample vigorously with a vortex mixer. We note that the internal agitator in the flow cytometer with a 300 rpm agitation speed was not sufficient to prevent the silica particles from sedimenting over time and we found that periodically agitating the sample tube every 5 minutes with a vortex mixer was more effective. Since all files must contain a fluorescein core, all particles were gated by default using the 'FITC' laser and detector settings, which is defined by gating the majority population in the 'FITC-A' channel histogram, in addition to standard FSC and SSC gates to minimize sorting of doublets (**Supplementary Fig. 11**). All FAS experiments were performed at room-temperature.

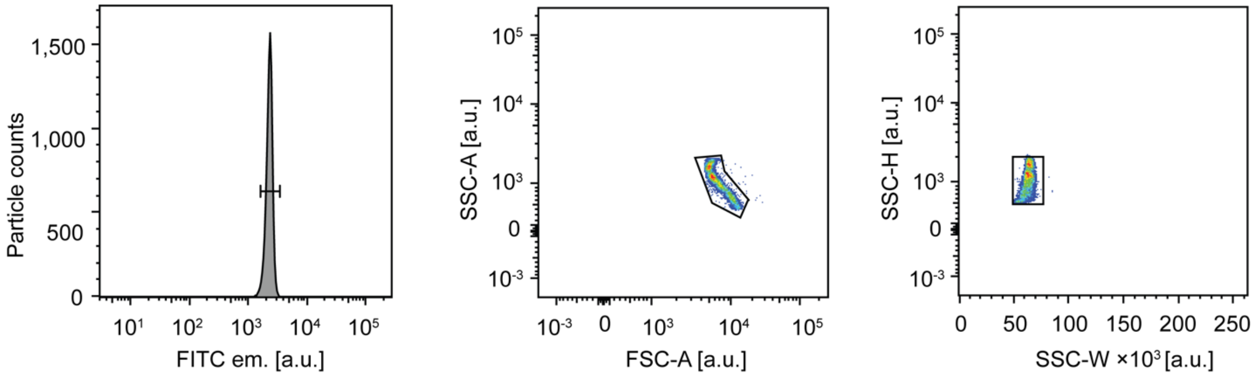

**Supplementary Figure 11. Example of a standard set of gates used in all particle sorts.** Colors indicate number density.

We also ran positive controls for every dye before every FAS experiment to validate that there is no significant spectral crosstalk during the sorting process and to validate that we have a distinguishable fluorescence signal in the presence of other fluorescent dyes. For example, because all our files have FITC, we validated that there is no fluorescence spillover of FITC in the TAMRA (PE-Texas Red channel), AF647 (APC channel), or TYE705 (Alexa Fluor 700 channel) that would otherwise make it difficult to distinguish the different particle populations. **Supplementary Fig. 12** summarizes the fluorescence traces of single and two-color controls.

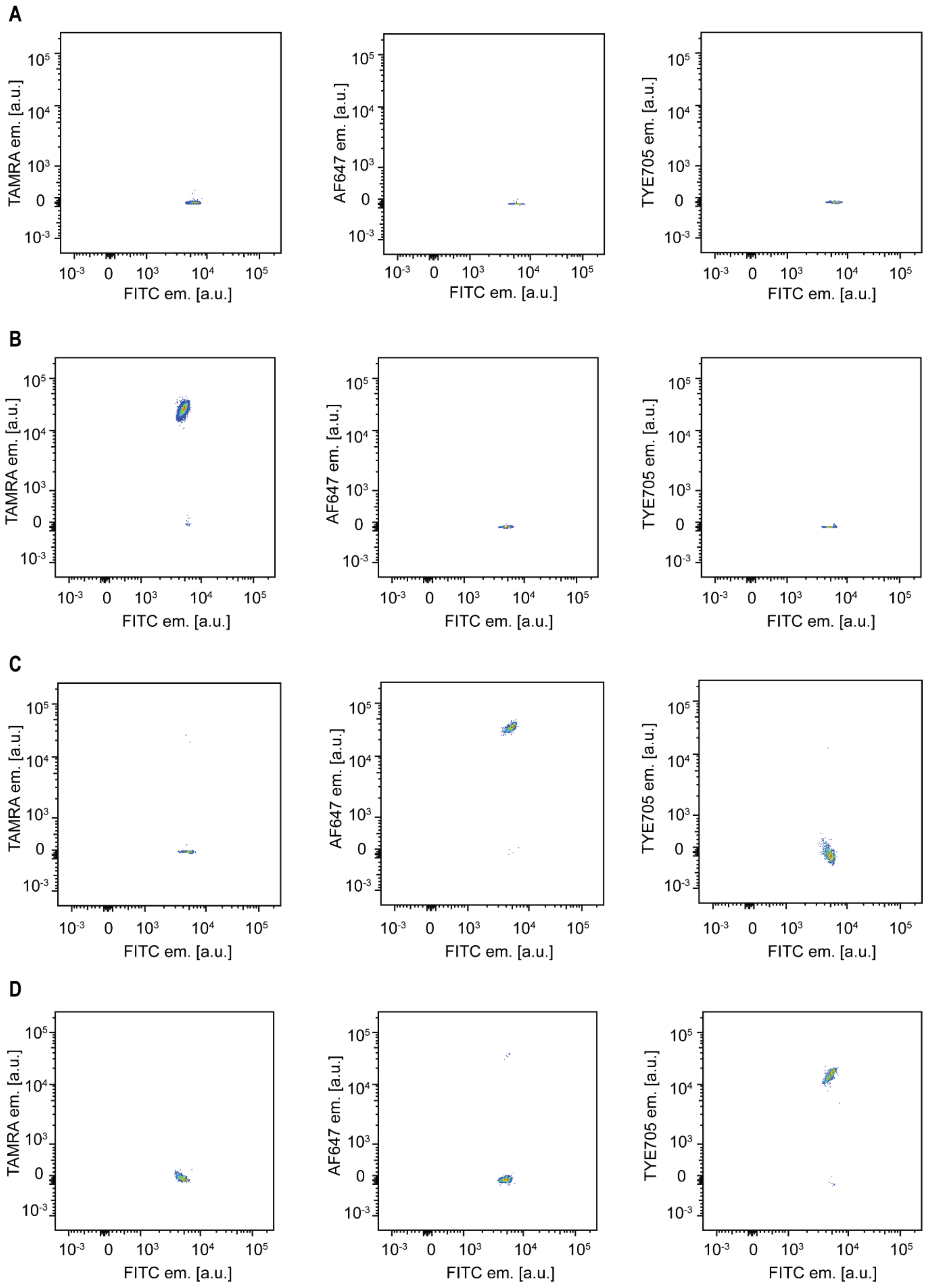

**Supplementary Figure 12. Single and two-color barcode positive controls.** (**A**) FITC only, (**B**) FITC + TAMRA, (**C**) FITC + AFDye 647, and (**D**) FITC + TYE705. Colors indicate number density.

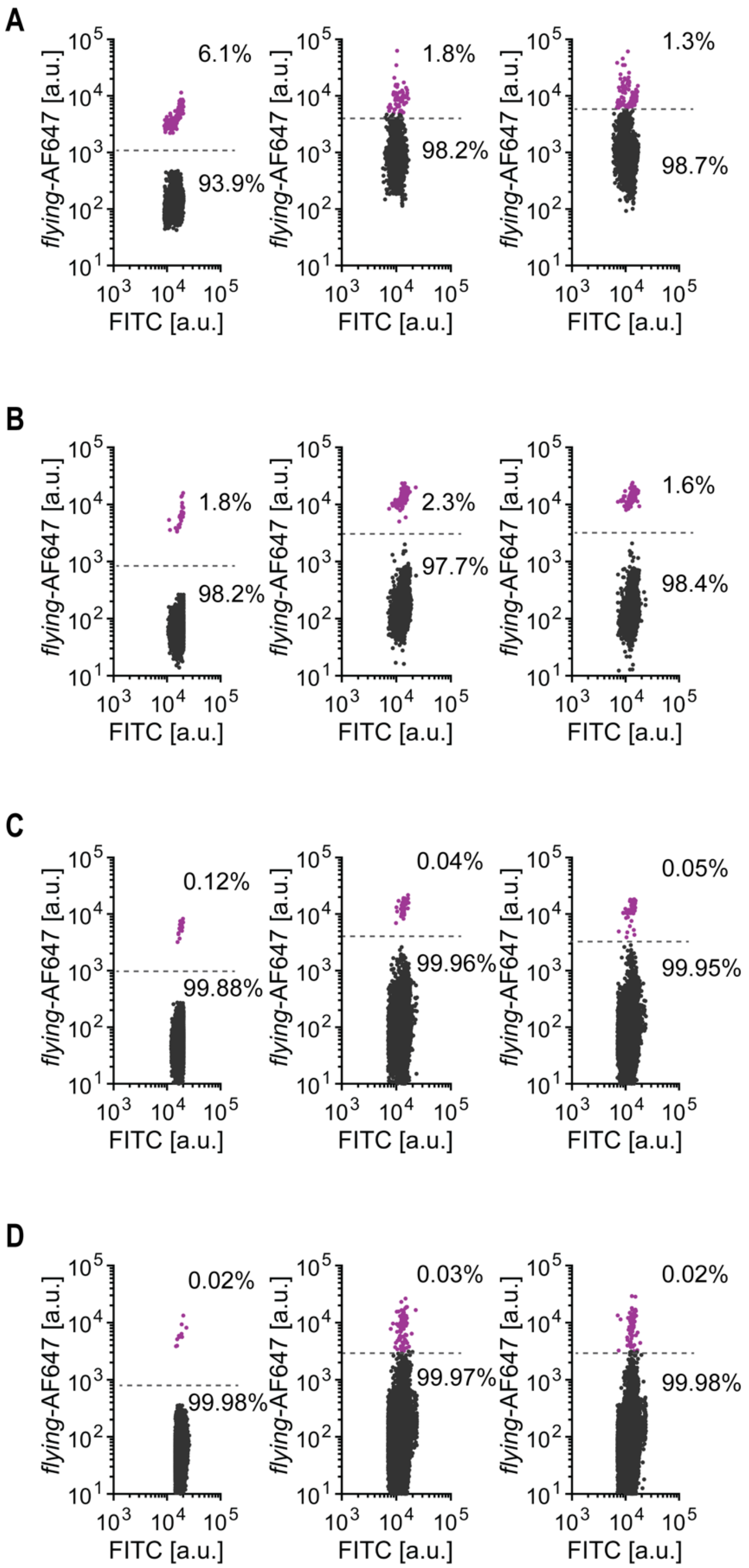

**Supplementary Figure 13. Raw data of FAS replicates from single barcode *flying* selections at different relative abundance of Airplane files compared to the other nineteen files.** (**A**) 1:1 ratio of *Airplane* to each other file, (**B**) $1:10^2$ ratio, (**C**) $1:10^4$ ratio, (**D**) $1:10^6$ ratio.

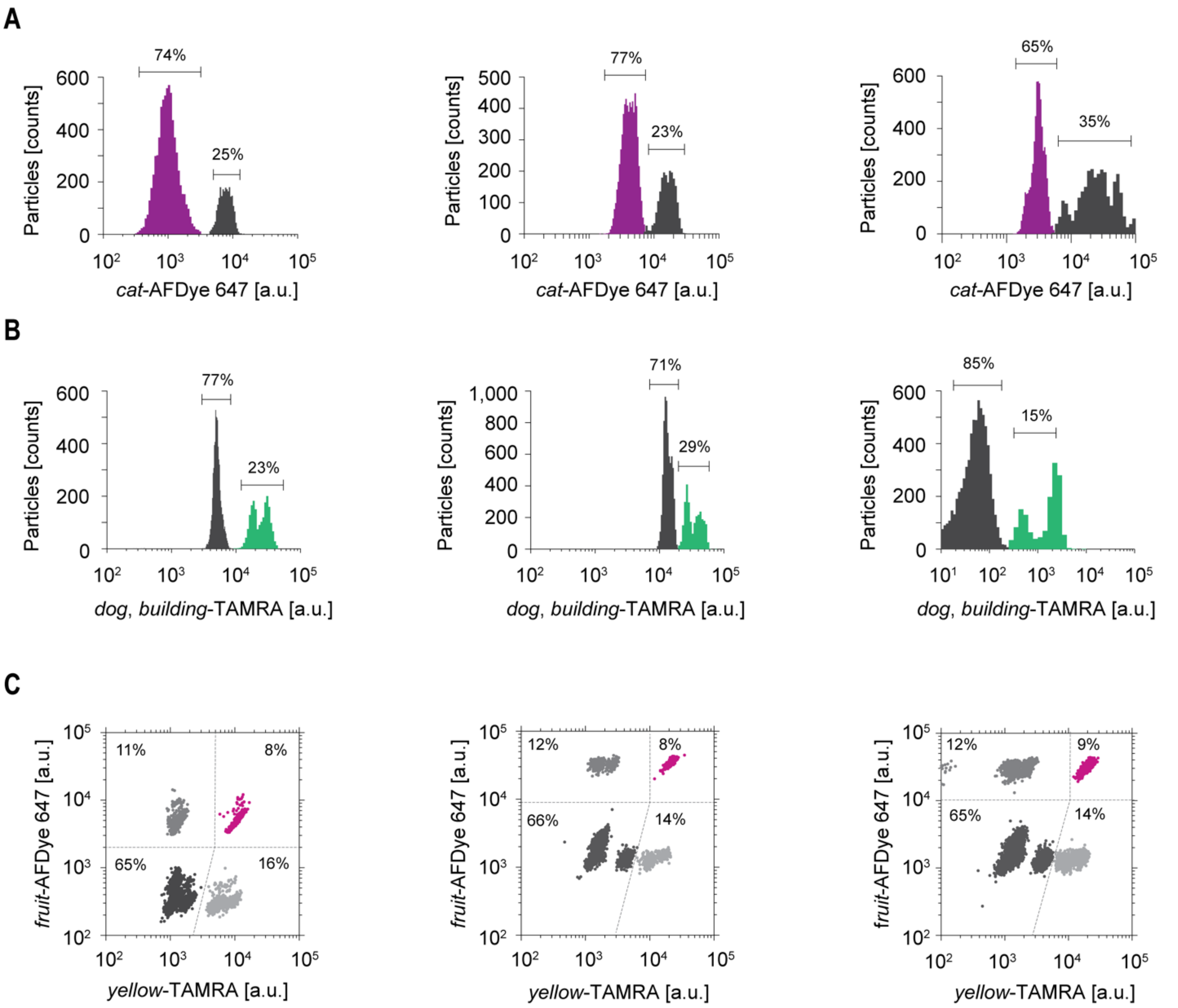

**Supplementary Figure 14. Raw data of FAS replicates from Boolean logic selections.** (**A**) `NOT cat`, (**B**) `dog OR building`, (**C**) `fruit AND yellow`.

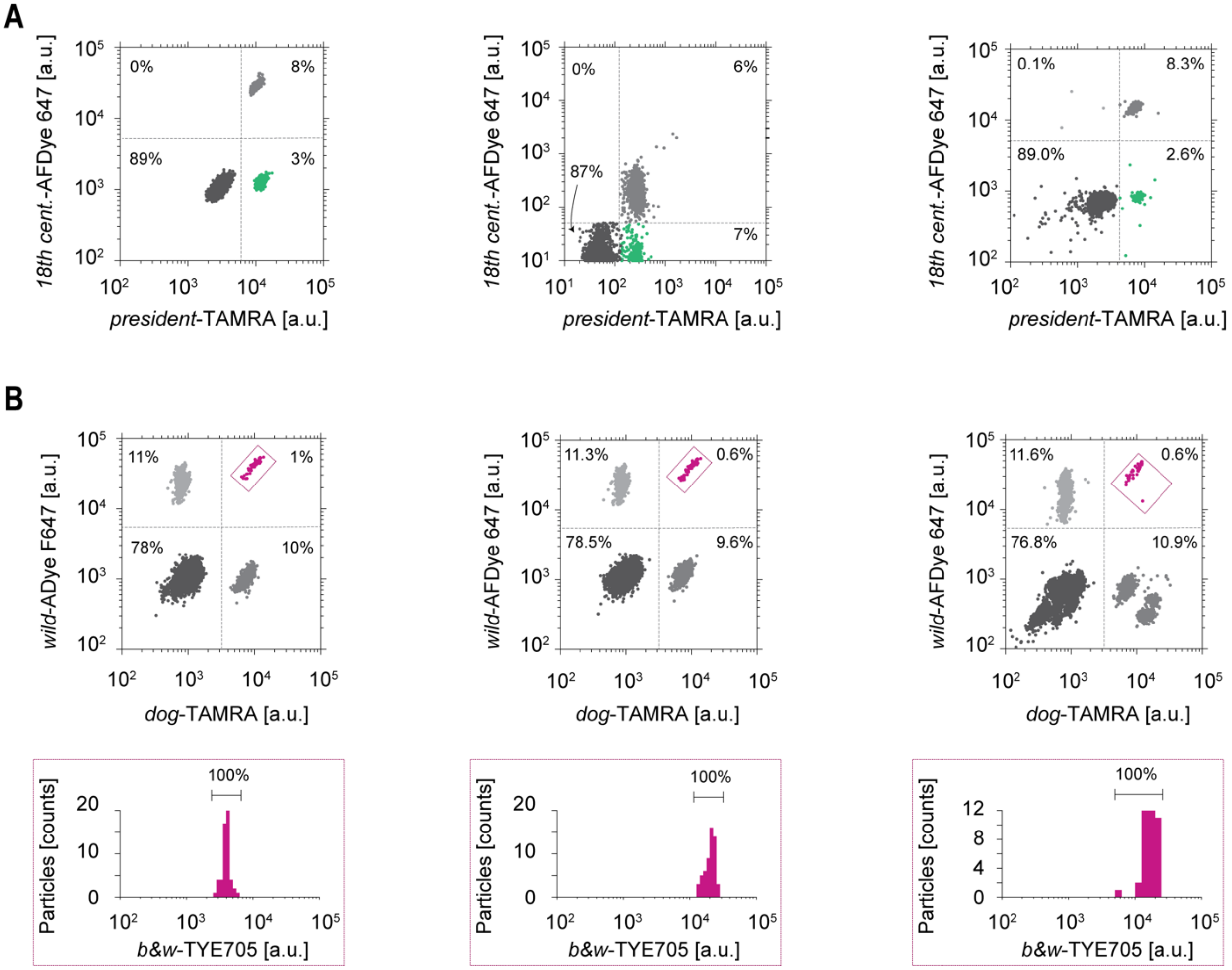

**Supplementary Figure 15. Raw data of FAS replicates from combined Boolean logic operations.** (**A**) `president AND (NOT 18`$^{th}$` century)`. (**B**) `dog AND wild`.

### S11. Verification of DNA retrieval from sorted sequences

**Release of DNA from sorted files.** Sorted populations were centrifuged at 10,000× g for 1 minute. The supernatant was carefully removed with a pipette to avoid disturbing the silica pellets. A volume of 45 µL of CMOS-grade 5:1 buffered oxide etch (Avantor; Visalia, CA) was then added. The mixture was vortexed for 5 seconds to re-suspend the pellet and the mixture was statically incubated at room temperature for 5 minutes. A volume of 5 µL of 1 M phosphate buffer (0.75 M $Na_2HPO_4$; 0.25 M $NaH_2PO_4$; pH 7.5 at 0.1 M) was then added, vortexed for 1 second, and desalted twice through an Illustra MicroSpin S-200 HR column, which we found to be most effective compared to other clean-up methods.

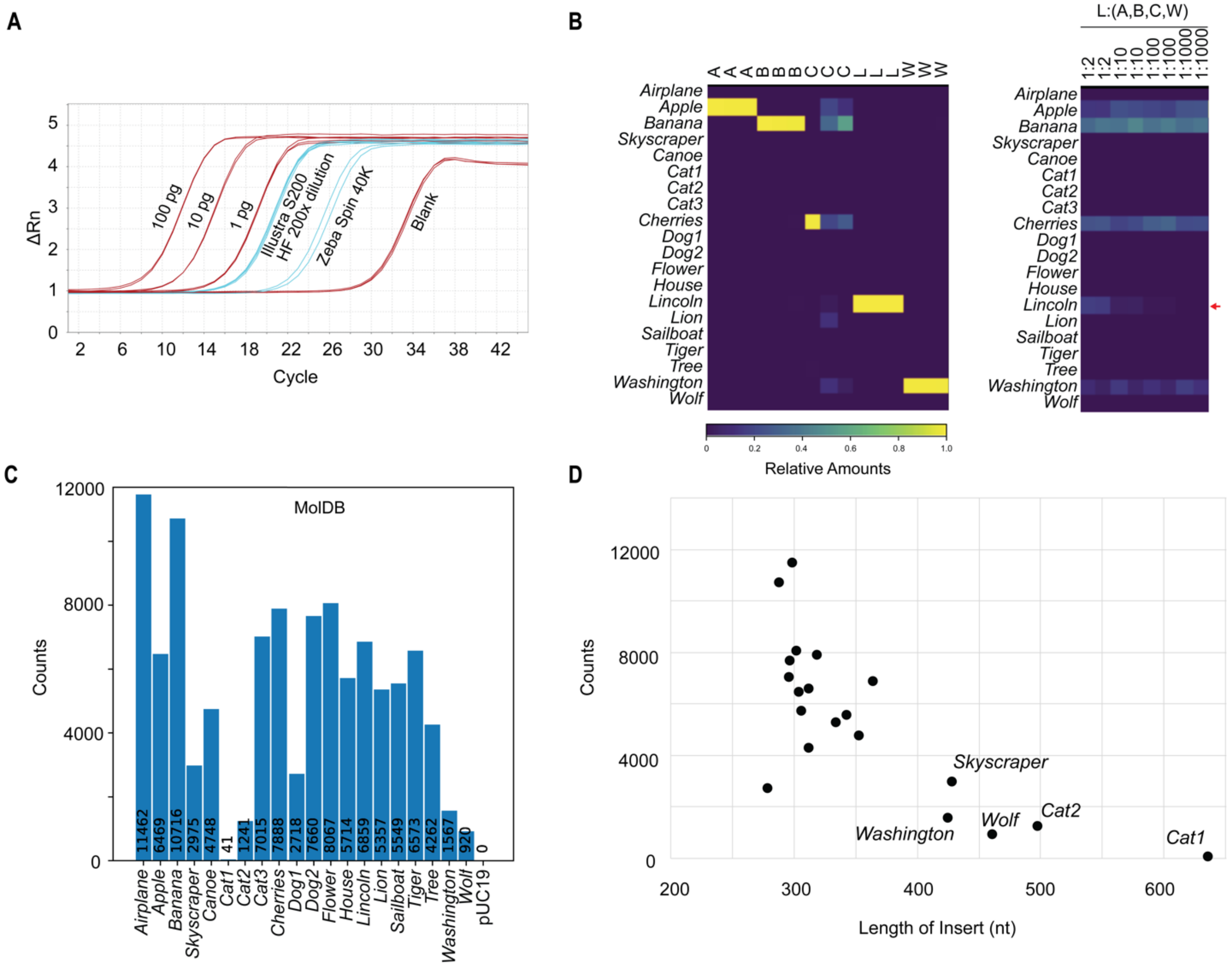

**Supplementary Figure 16. Cleanup and quality control on the release of DNA from the silica.** (**A**) Cleanup of release solution and additional salts from the release were tested by several methods of buffer exchange. Dilution of the released DNA mixture by 200-fold allowed for qPCR detection, as did cleanup by Illustra MicroSpin S-200 HR (GE Healthcare) and PCR Kleen Purification spin columns (Bio-Rad) with minimal sample loss. Zeba Spin 40K MWCO did not yield as much DNA. Illustra MicroSpin S-200 HR was used for all subsequent cleanup. (**B**) Release and purification of a subset of the molecular file database containing *Apple* (A), *Banana* (B), *Cherries* (C), *Lincoln* (L), and *Washington* (W) characterized from individual releases (left), and when part of a pool with *Lincoln* diluted (right), quantified by amplification, barcoding, and Illumina MiniSeq sequencing. (**C**) Release and purification of the entire molecular file database followed by amplification and barcoding, and sequencing from MiniSeq shows broadly similar profile of counts, with low detection of *Cat1*, and moderately low amplification of *Cat2*, *Dog1*, *Skyscraper*, *Washington*, and *Wolf*. (**D**) Low sequencing counts from the plasmid database can be explained by the variable length of the inserts being assayed. *Cat1*, *Cat2*, *Skyscraper*, *Washington*, *Wolf* are longer than 400 nucleotides.

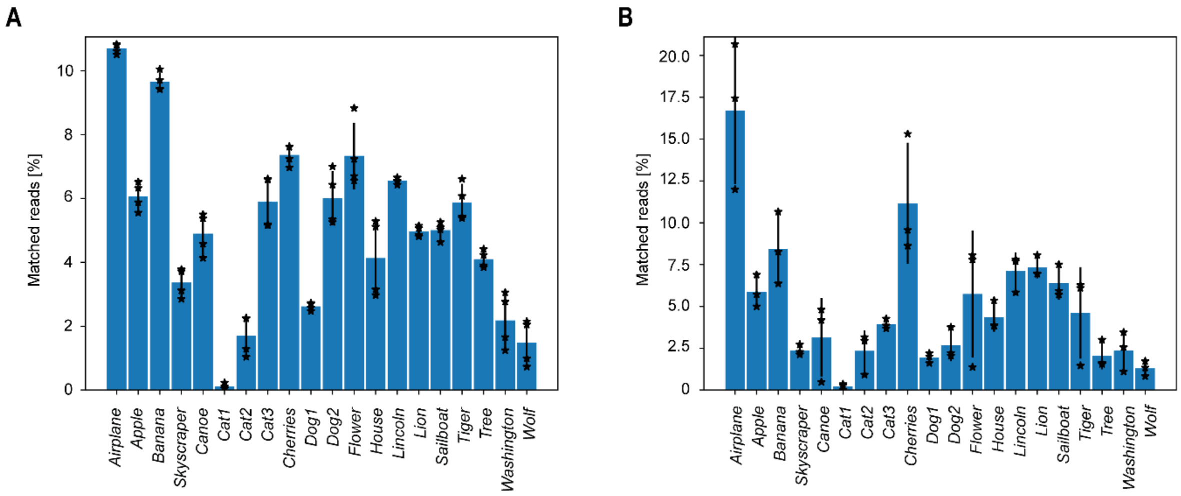

**Supplementary Figure 17. Count and sort probability statistics of sequencing reads from molecular file database.** (**A**) Molecular file database that did not pass through the FAS and were released directly. Mean and standard deviations were calculated from four independent replicates. (**B**) Molecular file database that was sorted into the FAS instrument using 'FITC' as the only sorting gate (all files contain FITC by default). Mean and standard deviations were calculated from three independent replicates. Matched reads are the number of reads matching each template divided by the number of reads matched to any template.

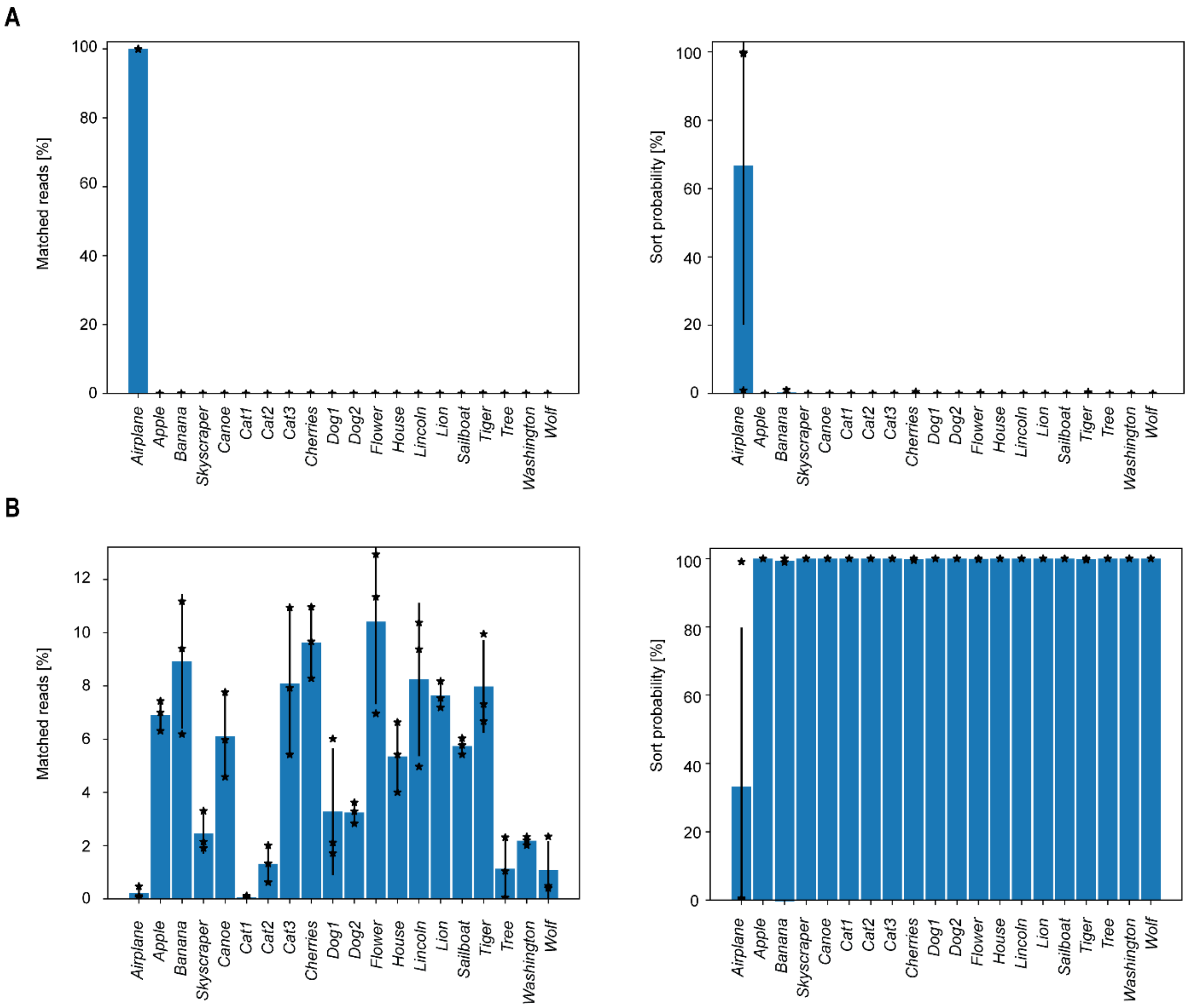

**Supplementary Figure 18. Count and sort probability statistics of sequencing reads from single-barcode sorts of *Airplane* in molecular database containing a 1:1 ratio of *Airplane* to each other file (i.e. equimolar concentration).** (**A**) Sorted populations from the `flying` gate. Left: matched reads, Right: sort probabilities. (**B**) Sorted populations from the `NOT flying` gate. Left: matched reads, Right: sort probabilities. Matched reads are the number of reads matching each template divided by the number of reads matched to any template. Mean and standard deviations were calculated from four independent replicates.

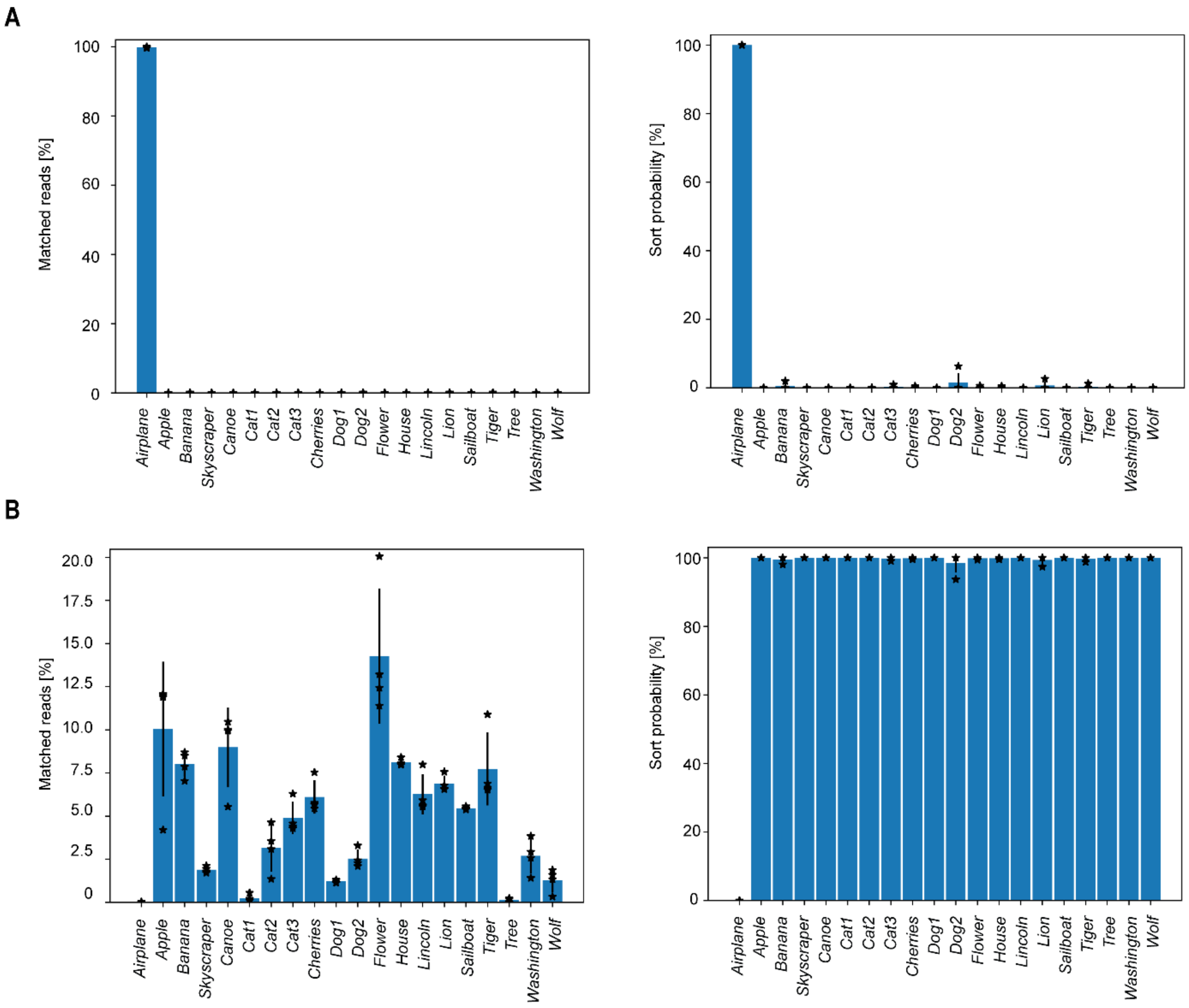

**Supplementary Figure 19. Count and sort probability statistics of sequencing reads from single-barcode sorts of *Airplane* in molecular database containing a 1:100 ratio of *Airplane* to each other file.** (**A**) Sorted populations from the `flying` gate. Left: matched reads, Right: sort probabilities. (**B**) Sorted populations from the `NOT flying` gate. Left: matched reads, Right: sort probabilities. Matched reads are the number of reads matching each template divided by the number of reads matched to any template. Mean and standard deviations were calculated from four independent replicates.

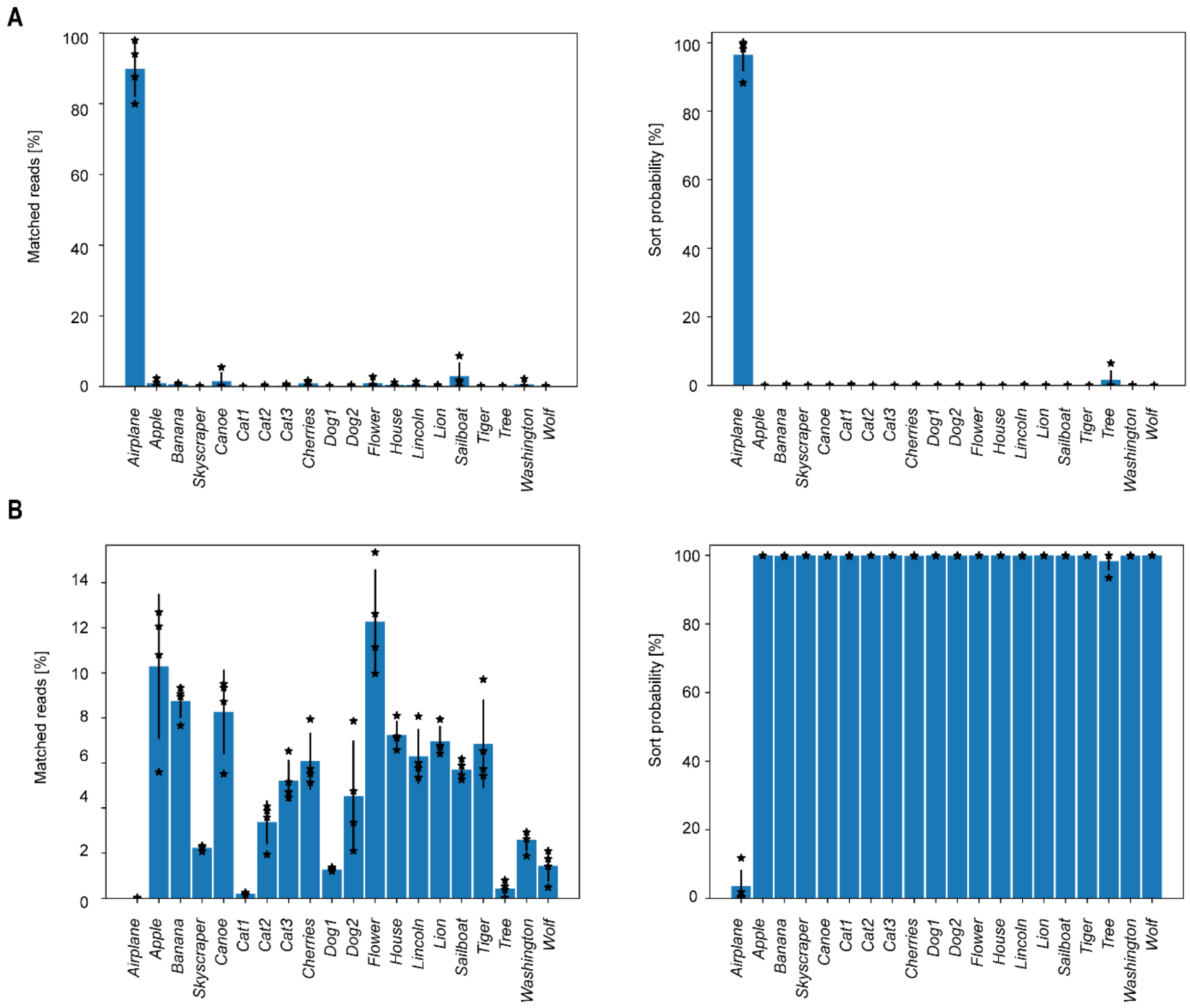

**Supplementary Figure 20. Count and sort probability statistics of sequencing reads from single-barcode sorts of *Airplane* in molecular database containing a 1:10,000 of *Airplane* to each other file.** (**A**) Sorted populations from the `flying` gate. Left: matched reads, Right: sort probabilities. (**B**) Sorted populations from the `NOT flying` gate. Left: matched reads, Right: sort probabilities. Matched reads are the number of reads matching each template divided by the number of reads matched to any template. Mean and standard deviations were calculated from four independent replicates.

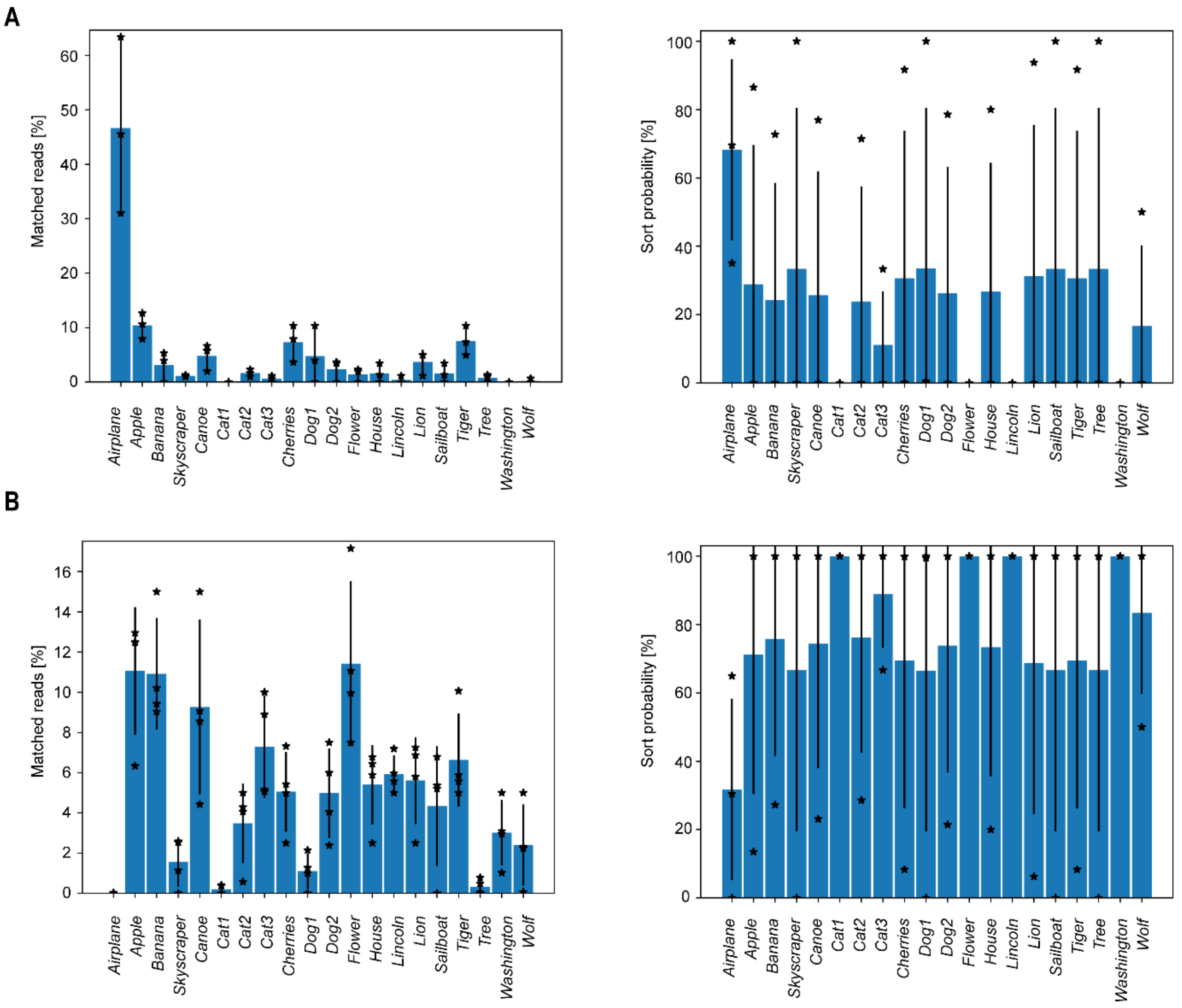

**Supplementary Figure 21. Count and sort probability statistics of sequencing reads from single-barcode sorts of *Airplane* in molecular database containing a 1:1,000,000 ratio of *Airplane* to each other file.** (**A**) Sorted populations from the `flying` gate. Left: matched reads, Right: sort probabilities. (**B**) Sorted populations from the `NOT flying` gate. Left: matched reads, Right: sort probabilities. Matched reads are the number of reads matching each template divided by the number of reads matched to any template. Mean and standard deviations were calculated from three independent replicates.

A

B

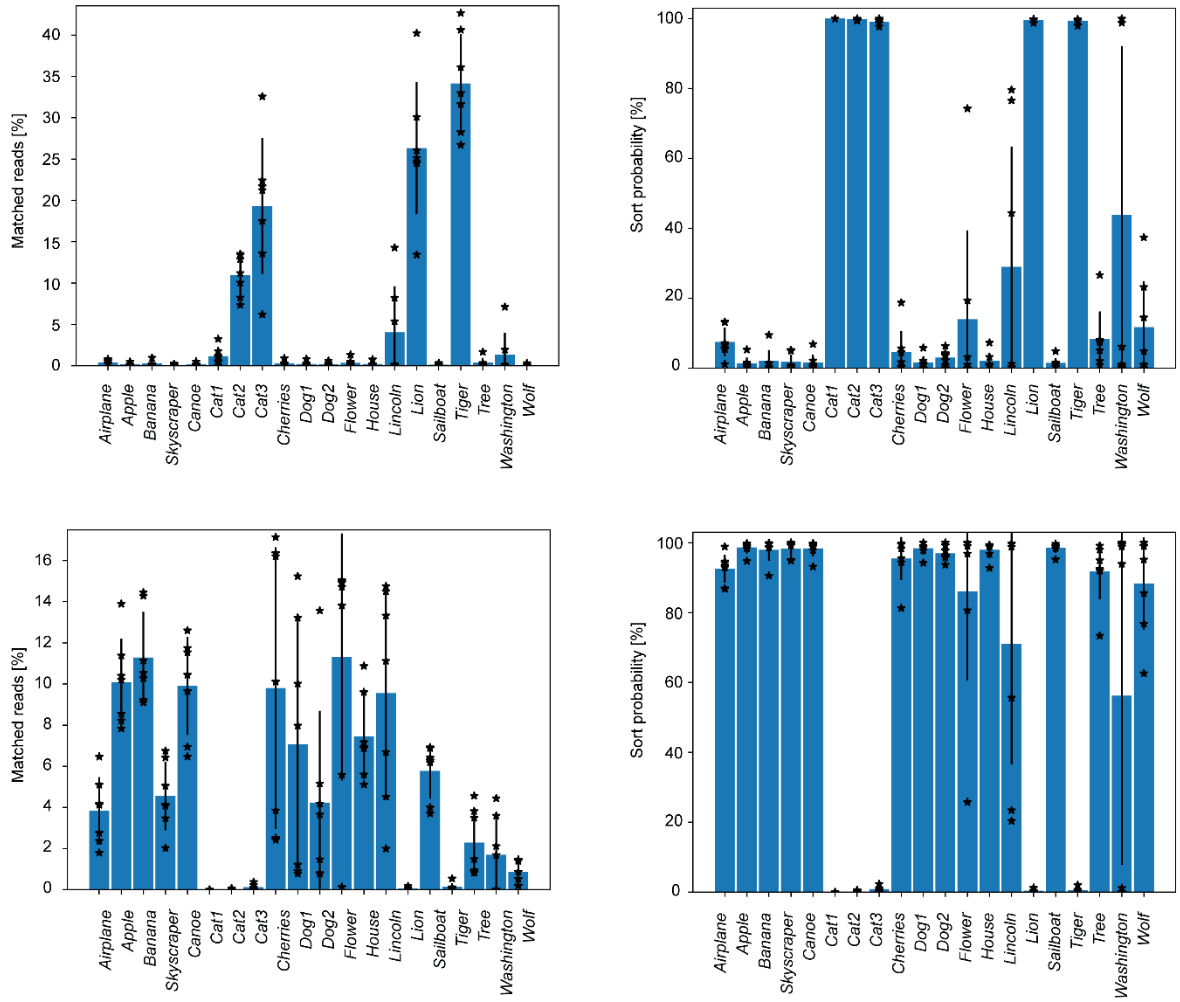

**Supplementary Figure 22. Count and sort probability statistics of sequencing reads from `NOT cat` sorts.** (**A**) Sorted populations from the `cat` gate. Left: raw counts, Right: sort probabilities. (**B**) Sorted populations from the `NOT cat` gate. Left: matched reads, Right: sort probabilities. Matched reads are the number of reads matching each template divided by the number of reads matched to any template. Mean and standard deviations were calculated from seven independent replicates.

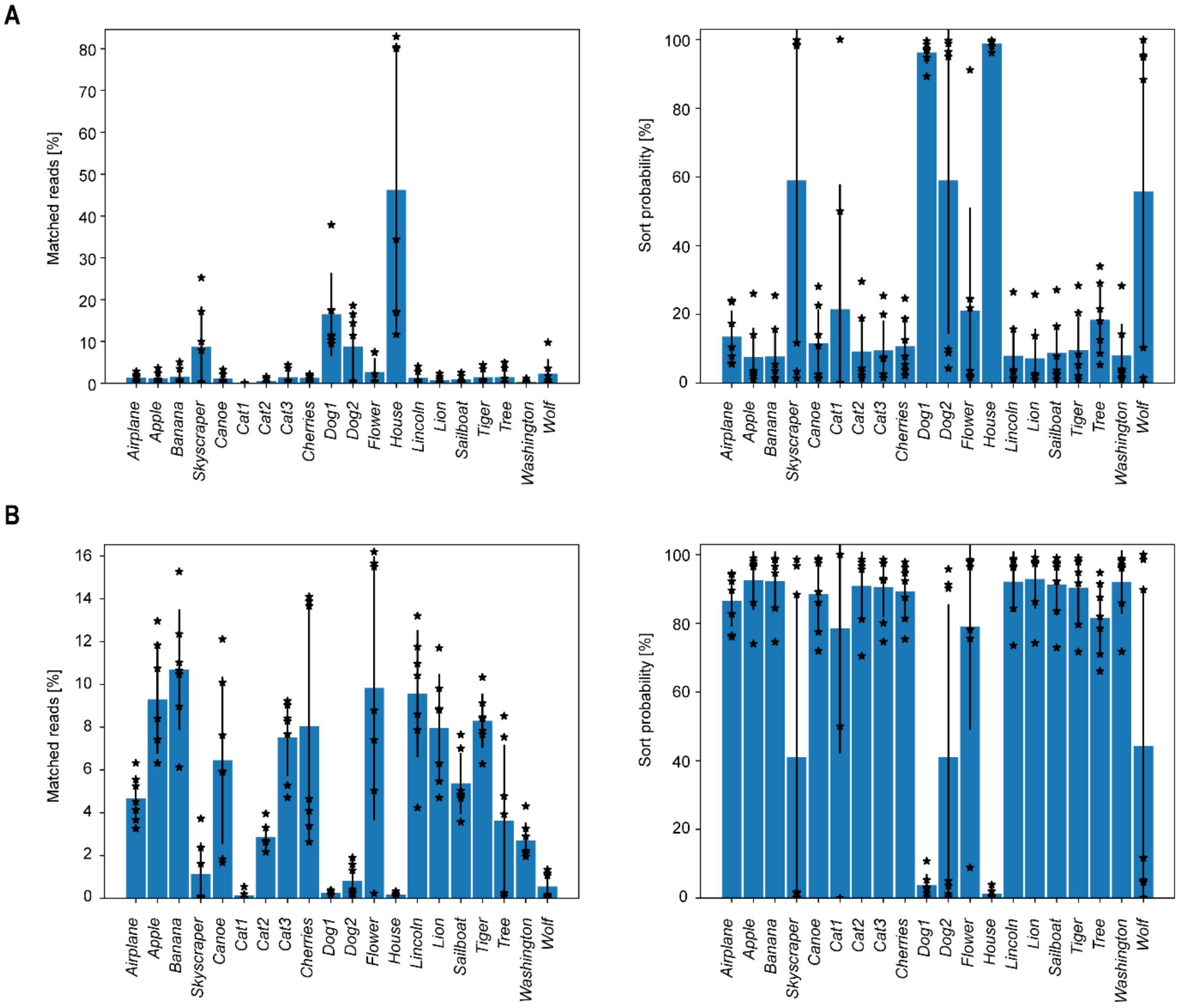

**Supplementary Figure 23. Count and sort probability statistics of sequencing reads from `dog OR building` sorts.** (**A**) Sorted populations from the `dog OR building` gate. Left: raw counts, Right: sort probabilities. (**B**) Sorted populations from the `NOT (dog OR building)` gate. Left: matched reads, Right: sort probabilities. Matched reads are the number of reads matching each template divided by the number of reads matched to any template. Mean and standard deviations were calculated from seven independent replicates.

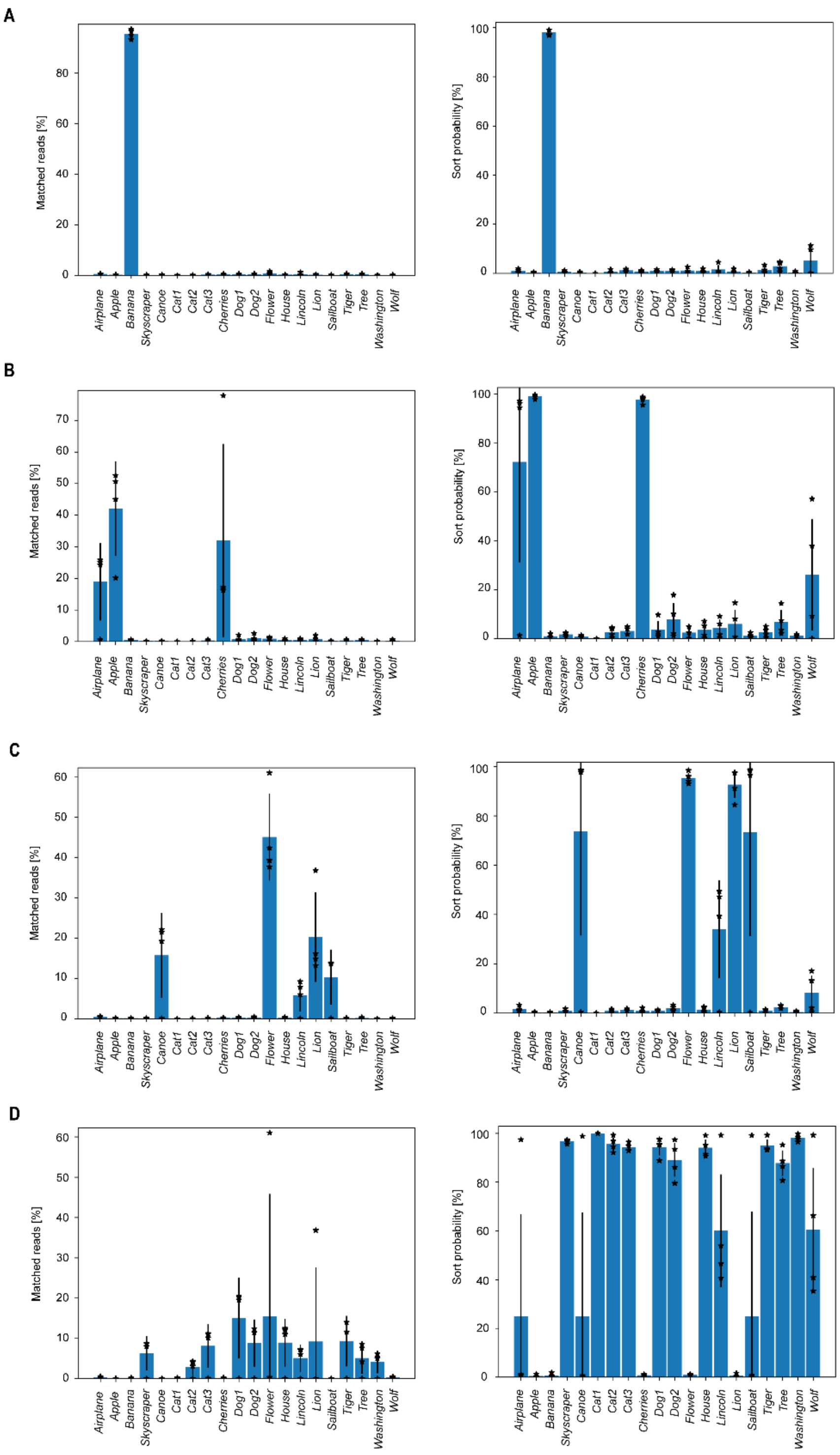

**Supplementary Figure 24. Count and sort probability statistics of sequencing reads from `yellow AND fruit` sorts.** (**A**) Sorted populations from the `yellow AND fruit` gate. Left: matched reads, Right: sort probabilities. (**B**) Sorted populations from the `NOT yellow AND fruit` gate. Left: matched reads, Right: sort probabilities. (**C**) Sorted populations from the `yellow AND NOT fruit` gate. Left: matched reads, Right: sort probabilities. (**D**) Sorted populations from the `NOT fruit AND NOT yellow` gate. Left: matched reads, Right: sort probabilities. Mean and standard deviations were calculated from four independent replicates. Matched reads are the number of reads matching each template divided by the number of reads matched to any template.

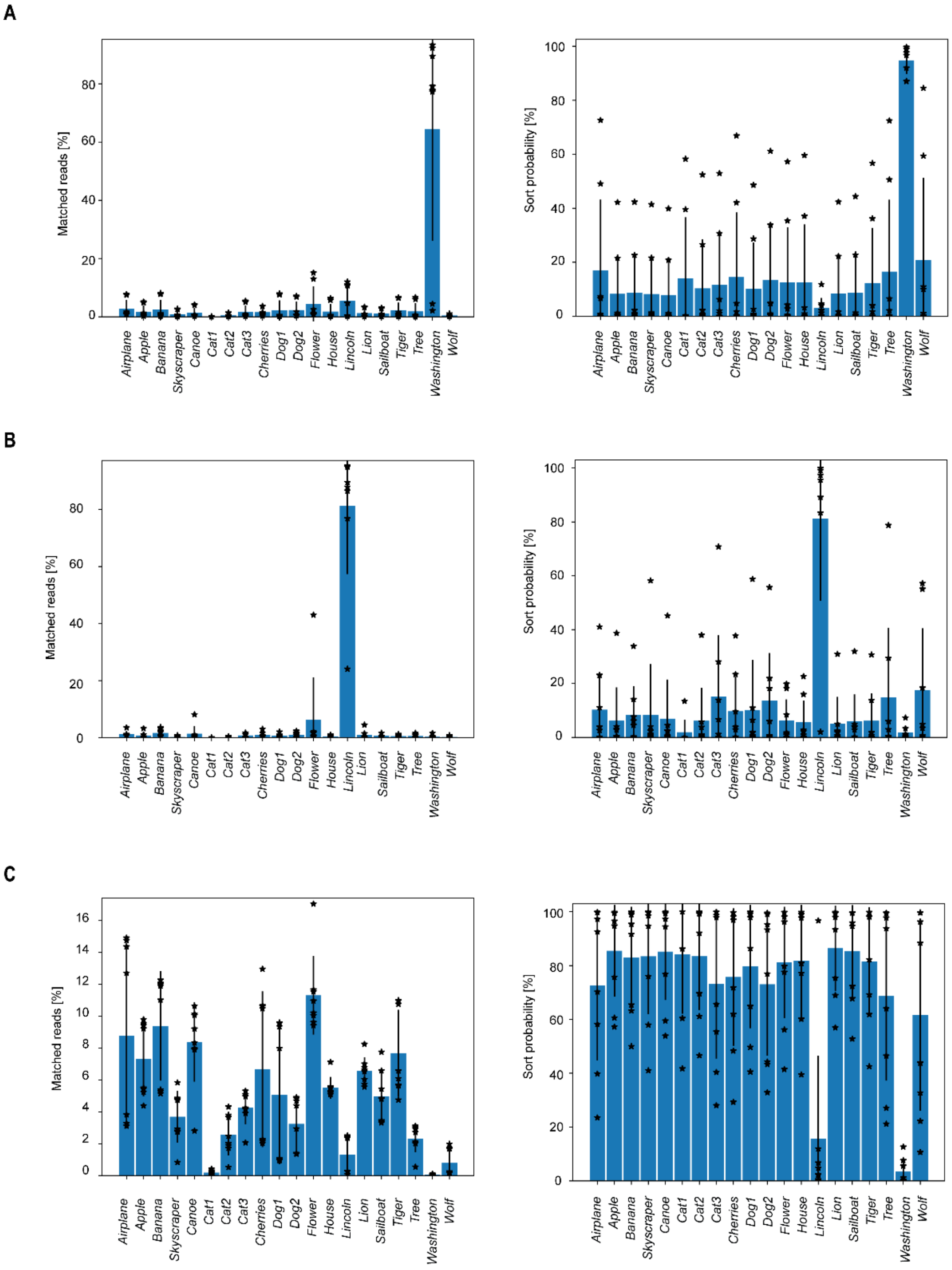

**Supplementary Figure 25. Count and sort probability statistics of sequencing reads from `president AND 18th century` sorts.** (**A**) Sorted populations from the `president AND 18th century` gate. Left: matched reads, Right: sort probabilities. (**B**) Sorted populations from the `president AND NOT 18th century` gate. Left: matched reads, Right: sort probabilities. (**C**) Sorted populations from the `NOT president` gate. Left: matched reads, Right: sort probabilities. Matched reads are the number of reads matching each template divided by the number of reads matched to any template. Mean and standard deviations were calculated from eight independent replicates.

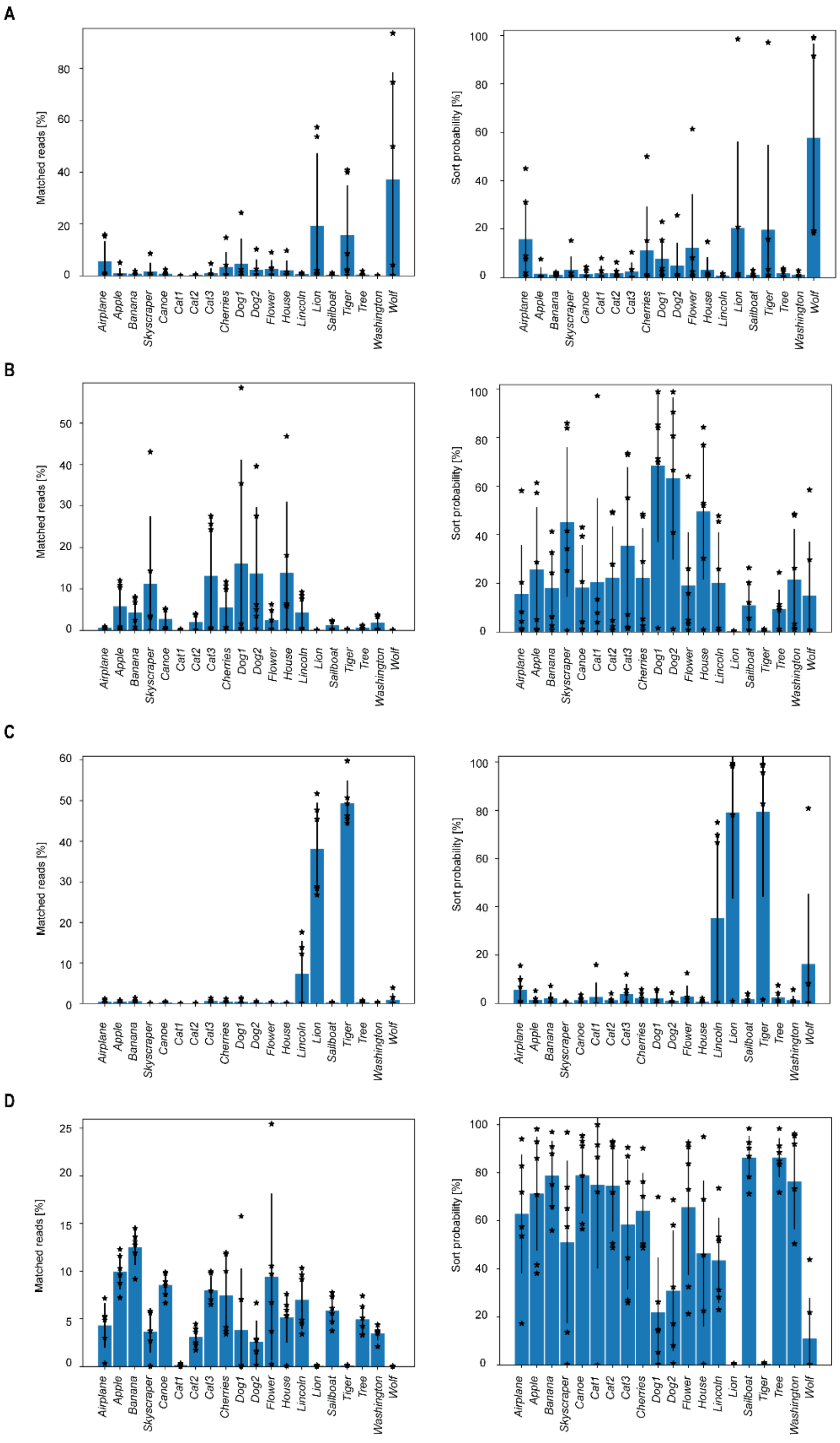

**Supplementary Figure 26. Count and sort probability statistics of sequencing reads from `dog AND wild` sorts.** (**A**) Sorted populations from the `dog AND wild` gate. Left: matched reads, Right: sort probabilities. (**B**) Sorted populations from the `dog AND NOT wild` gate. Left: matched reads, Right: sort probabilities. (**C**) Sorted populations from the `NOT dog AND wild` gate. Left: matched reads, Right: sort probabilities. (**D**) Sorted populations from the `NOT dog AND NOT wild` gate. Left: matched reads, Right: sort probabilities. Matched reads are the number of reads matching each template divided by the number of reads matched to any template. Mean and standard deviations were calculated from six independent replicates.

### S12. Bacterial transformation of sorted sequences

Samples that were sorted to single populations (`yellow AND fruit`: *Banana*; `president AND 18th century`: *Washington*; `president AND NOT 18th century`: *Lincoln*) were transformed to chemically competent *E. coli* 10β cells (NEB). Of the transformed cells, three colonies from each were grown in 4-mL LB overnight at 37 °C and Qiagen miniprep spin purification was used to retrieve the plasmid. Each of the three plasmids, as well as the PCR amplified release from each of the three populations, were sent for Sanger sequencing. All three amplicons showed primarily expected DNA sequences and each of the three images were retrieved from the sequencing of the sorts. Of the transformed colonies, 2 out of the 3 of the *Lincoln* colonies were verified to be *Lincoln* plasmids (one *Canoe* colony was also retrieved from this sample), while 3 out of the 3 *Banana* and *Washington* colonies were recovered. Sanger sequencing of the nine colonies showed no errors in results and all images were retrieved by inverse DNA to image processing. The bacterially amplified DNA was pure and readily available for re-encapsulation in a closed read-write cycle.

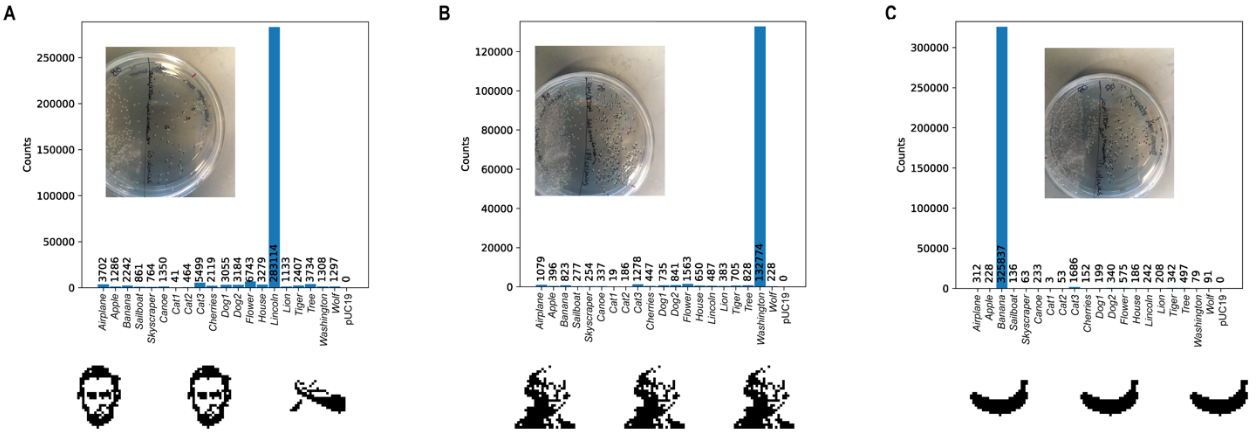

**Supplementary Figure 27. Bacterial transformation with sorted and cleaned DNA.** Cleanup of DNA release solution and additional salts away from the DNA allowed for transformation of NEB DH10β cells with the purified DNA. DNA sorted to single populations from (**A**) `president AND NOT 18`$^{th}$` century`. (**B**) `president AND 18`$^{th}$` century`, and (**C**) `yellow AND fruit` were transformed and grown to single colonies, and 3 colonies were selected and grown for DNA preparation and Sanger sequencing was applied to each. The expected Lincoln sort yielded two positive colonies and one colony encoding *Canoe* (**A**, bottom), with *Washington* (**B**, bottom) and *Banana* (**C**, bottom) sorts showed all three colonies returning the expected encoded image.
